# Enrichment of Cysteine S-palmitoylated peptides using Sodium Deoxycholate Acid Precipitation - SDC-ACE

**DOI:** 10.1101/2023.07.09.548252

**Authors:** Peter T. Jensen, Giuseppe Palmisano, Christopher J. Rhodes, Martin R. Larsen

## Abstract

S-palmitoylation is a poorly understood post-translational modification that is gaining more attention as an essential regulator of cellular processes. The reversible nature of S-palmitoylation allows for fine-tuned control of cellular events and adaptation to stimuli. The enrichment of S-palmitoylated proteins and peptides includes the Acyl-Biotin Exchange (ABE) method, Acyl resin-assisted Capture (Acyl-RAC), metabolic labelling, and derivatives thereof. We present a novel method of enrichment of S-palmitoylated peptides termed *SDC Acid Precipitation Enrichment* (SDC-ACE). Here, S-palmitoylated peptides are enriched by taking advantage of their co-precipitation with Sodium-Deoxycholate (SDC) under acidic conditions, allowing easy and fast separation of lipidated peptides from the sample suspension. We applied our novel method for the characterization of the mouse brain, providing an in-depth analysis of S-palmitoylation events within the brain and comprehensive profile of the mouse brain S-palmitoylome. Furthermore, we applied our method for mapping mouse tissue-specific S-palmitoylation, highlighting the extensive role of S-palmitoylation throughout various organs in the body. Finally, we applied our methods for studying the brain palmitoylome of diabetic db/db mouse, uncovering alterations in the palmitoylation related to obesity and type 2 diabetes. The SDC-ACE method allows fast and easy enrichment of S-palmitoylated peptides, providing a valuable tool for exploring the dynamics and function of S-palmitoylation in diverse biological systems.

## Introduction

Protein lipidation is an important post-translational modification (PTM) that plays critical roles in regulating protein location, function, and activity within cells. This process involves the covalent attachment of lipid molecules, such as fatty acids, to specific amino acids in proteins. The significance of protein lipidation lies in its ability to dynamically modulate various aspects of protein behaviour, such as binding affinities, stability, folding patterns, cellular localization, and interaction with other macromolecules[1]. Given its central role in numerous cellular functions — including trafficking, migration, and signal transduction — protein lipidation is a key focus of biomedical research. Importantly, abnormalities in protein lipidation are linked to various diseases such as cancer (reviewed in [2]), neurodegenerative conditions (reviewed in [3]), and metabolic disorders [4–6], making protein lipidation an area of keen interest for biomedical research and highlighting its potential as a target for therapeutic intervention.

Among the different types of protein lipidation, Cysteine S-acylation is the most prevalent. This modification involves a two-step enzymatic addition of a fatty acid moiety to the sulphur atom of cysteine residues in a protein via a thioester linkage [7–9]. In contrast to most protein lipid modifications, S-acylation is reversible allowing for the dynamic regulation of protein function, activity, and cellular localization, thereby making it a critical player in cellular signalling pathways and other physiological processes [10, 11]. Although different long chain acyl moieties can be conjugated to proteins’ appropriately exposed Cysteine, S-acylation is more commonly referred to as S-palmitoylation (or palmitoylation) since the majority of S-acylation modifications commonly utilize the 16-carbon fatty acid palmitate. The enzymes responsible for S-palmitoylation are protein acyltransferases, a family of 23 genes in humans [12] and 24 in mouse [13], also termed the S-palmitoylation “writers”. These enzymes are integral membrane proteins and feature a highly conserved aspartate-histidine-histidine-cysteine (DHHC) motif housed within a zinc-binding cysteine-rich domain (zDHHC), which is vital for their catalytic activity [7–9, 14, 15]. Several zDHHC palmitoyl-transfrerases (PATs) are themselves S-palmitoylated by other PATs, regulating their membrane association and activity [16, 17]. One example is the regulatory interplay between zDHHC6 and zDHHC16, where the protein stability and catalytic activity of zDHHC6 is regulated through zDHHC16-catalyzed palmitoylation of three cysteines on the cytosolic C-terminal tail of zDHHC6 [16]. In contrast to the relative high number of S-palmitoylation “writers”, only few proteins have been identified as S-palmitoylation “erasers”, including the Acyl-protein thioesterase 1 and 2 (APT1/2), protein palmitoyl thioesterase (PPT) and members of the Alpha/beta hydrolase domain-containing (ABHD) protein family [3].

Studying S-palmitoylation is a complex analytical task due to the attachment of the fatty acid modifying group which bind to surfaces and chromatographic resin, complicating the isolation and identification of intact S-palmitoylated peptides. Therefore, most methods developed for studying S-palmitoylation sites in proteins are relying on sequential reduction of the S-palmitoyl group combined with specialized affinity-based techniques and identification of formerly S-palmitoylated peptides using liquid chromatography coupled with tandem mass spectrometry (LC-MS/MS). These methods include acyl-biotin exchange (ABE) [18, 19] and methods related to ABE. ABE is the most widely used method that involves the complete alkylation of free cysteines on the target proteins to avoid false positive identifications, followed by the removal of the S-palmitoyl group on cysteines using high molar concentrations of hydroxylamine (HA) [20]. After the S-palmitoyl group is removed, the newly freed cysteine thiol groups are reacted with a biotin molecule and subsequently, these biotinylated, formerly S-palmitoylated proteins are enriched by capture on streptavidin/avidin beads. This approach was first adapted by Roth et al., for global analysis of protein S-palmitoylation by releasing proteins from the biotin moity, followed by cysteine alkylation and tryptic digestion prior to LC-MS/MS to identify and quantify the formerly S-palmitoylated protein/peptides [21]. Yang et al., reported a different adaptation, named palmitoyl protein identification and site characterization (PalmPISC) using the ABE method with affinity enrichment of biotinylated S-palmitoylated proteins at the peptide-level, allowing the identification of 398 S-acylated proteins and 168 S-acylation sites, making the approach more suitable for large-scale characterization of S-palmitoylated sites [22]. Acyl resin-assisted capture (Acyl-RAC), developed by Forrester et al., is another affinity-based approach utilizing methanethiosulfonate (MMTS) to alkylate free cysteines followed by hydroxylamine treatment and capture of formerly S-palmitoylated cysteines with thiol-reactive thiopropyl Sepharose resin [23]. Collins et al., further developed on the ABE approach, reporting a new method named site-specific ABE (ssABE), utilizing a molecular weight-based cut-off column concentrator to reduce protein loss and eliminate labour-intensive protein precipitation, ensuring the removal of chemical reagents, which can interfere with subsequent reactions [24]. Using this approach, they identified 906 putative S-palmitoylation sites in 641 proteins from mouse forebrain.

For ABE and related methods that has been developed over the years, researchers have used either Tris buffer as a control, which theoretically should not release the S-palmitoyl group from cysteines.

As an alternative method to affinity-based techniques for characterization of S-palmitoylation, metabolic labelling approaches employ bio-orthogonal probes [25], such as 17-octadecyonic acid (17-ODYA) [26], has been used. The 17-ODYA labelled sites can subsequently be identified or visualized using click chemistry with biotin-probes combined with LC-MSMS analysis, Western blotting, or imaging (e.g., [26]).

Here we present a simple and fast method for enriching and characterizing formerly S-palmitoylated peptides using sodium deoxycholate (SDC) liquid-solid separation combined with acid precipitation and dithiothreitol (DTT) reduction, termed SDC Acid Precipitation Enrichment (SDC-ACE). We optimized the method on synthetic S-palmitoylated peptides and applied it to different biological settings, including mouse organs and obesity linked diabetic mouse brains. In total, we identified more than 33,000 formerly S-acylated peptides from mouse brain membrane preparations. We highlight tissue-specific S-palmitoylation events in mice organs, including synaptic signaling in brain, ion channels and transporters in kidney, and metabolic processes and bile secretion in liver. We identified S-palmitoylation of many enzymes involved in regulation of other PTMs, including kinases and phosphatases, strongly suggesting PTM crosstalk between S-palmitoylation and other PTMs. We also identify extensive S-palmitoylation on zDHHC enzymes, which could serve as a reservoir for palmitoyl moieties for substrate S-palmitoylation. Finally, we identified several significantly regulated S-palmitoylated peptides in obesity linked diabetic mouse brain, including S-palmitoylation belonging to proteins involved in insulin and leptin signaling pathways, further linking S-palmitoylation with metabolic disorders.

## Results

### Development of the SDC Acid Precipitation Enrichment (SDC-ACE) method for S-palmitoylation

SDC is a frequently used detergent in proteomics for extracting proteins from cells and tissues, since proteases, such as trypsin, retain activity in SDC [27]. SDC is easily removed using either phase transfer extraction with a water-immiscible organic solvent such as ethyl acetate [28] or acid precipitation followed by centrifugation [29]. Whereas phase transfer extraction is difficult and often result in contamination due to the top ethyl acetate phase, the acid precipitation is fast and easy. However, acid precipitation results in a loss of peptides to the SDC pellet. This is illustrated by the tryptic digestion of an aliquot of proteins extracted from HeLa cells using 1% SDC. The content of peptides was measured using Amino Acid Composition analysis before and after acid precipitation and centrifugation (**Supplementary fig. S1A**). The acid precipitation resulted in almost 30% loss of peptide amount to the SDC pellet. When washing the SDC pellet by re-solubilization in pH 8-8.5 buffer followed by acid precipitation, more peptides can be recovered from the SDC pellet [30]. This is illustrated here by the tryptic digestion of proteins extracted from a mouse brain membrane preparation using 5% SDC (**Supplementary fig. S1B**). Here the recovery of primarily increasingly hydrophobic peptides could be detected in repeating washes of the SDC pellet (**Supplementary fig. S1B**). This together with the relatively large amount of remaining material in the SDC pellet, indicate that very hydrophobic peptides bind tightly to the precipitated SDC. We therefore hypothesized that lipid modified peptides are enriched in the SDC pellet.

To investigate this hypothesis, three synthetic S-palmitoylated peptides (peptide sequence A, B and C) were generated using TFA and palmitoyl chloride [31] (**Supplementary fig. S2A-C**). To test if S-palmitoylated peptides co-precipitate with the SDC after acidification and centrifugation, the three S-palmitoylated peptides (A-, B-, and C-Palm) were combined and analyzed by MALDI-MS (**Fig 1Aa**). SDC was added to the mixture to a final concentration of 1% w/v followed by acidification and centrifugation. MALDI-MS analysis of the supernatant revealed their complete removal (**Fig. 1Ab**), indicating efficient enrichment of the S-palmitoylated peptides with SDC acidic precipitation. Analysis of the intact S-palmitoylated peptides from the SDC pellet was not possible by LC-MS analysis due to the high amount of SDC present.

**Figure 1:**
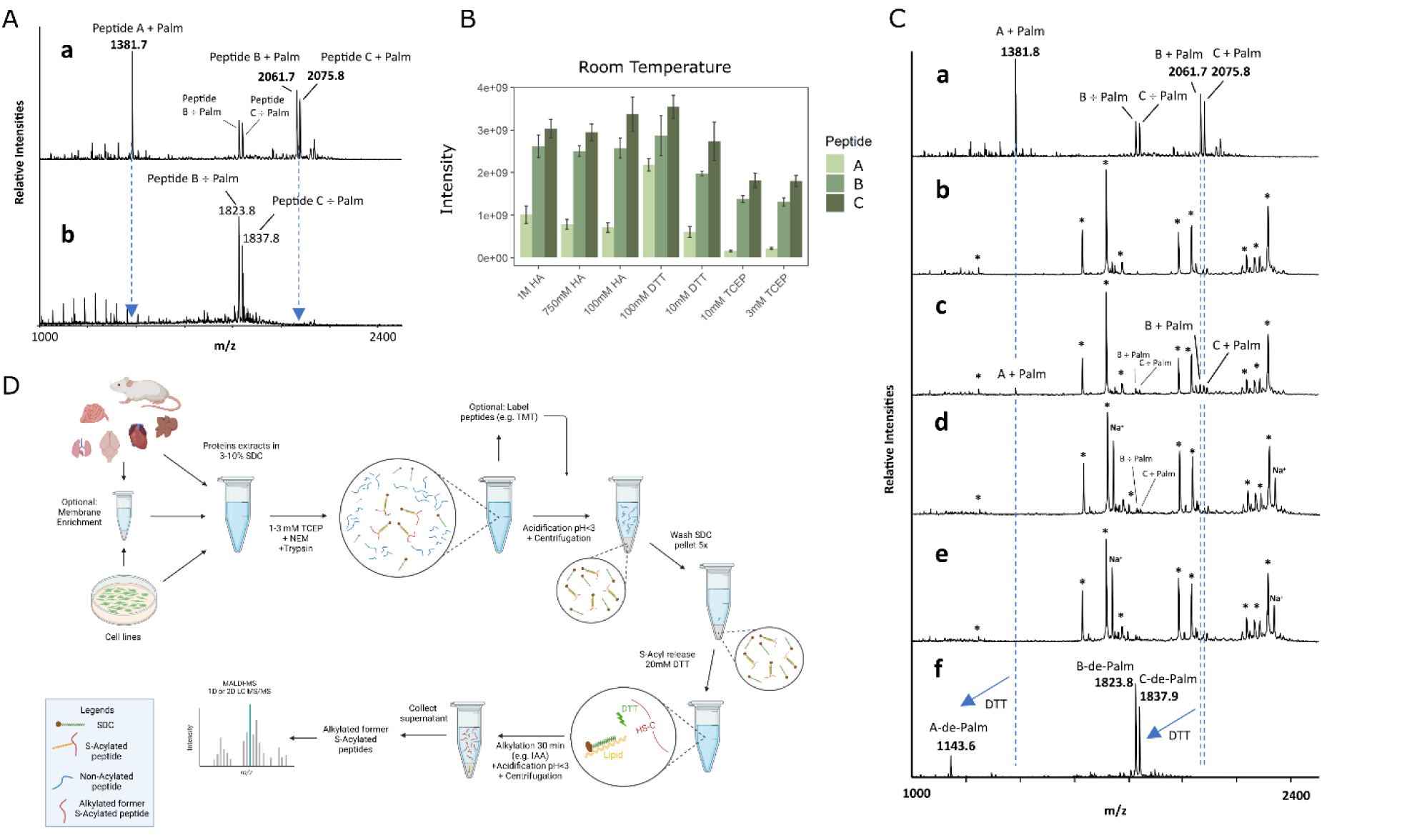
Development of the SDC-ACE method for enrichment of S-palmitoylated peptides. **A**) (a) MALDI MS analysis of synthetic S-palmitoylated peptides (b) MALDI MS analysis of supernatant after SDC precipitation. **B**) Validation of the most efficient reducing reagent for releasing S-palmitoylation from the synthetic S-palmitoylated peptides. DTT was comparably more efficient at releasing S-palmitoylation compared to HA and TCEP. **C**) Specificity of the SDC-ACE method demonstrated by mixing synthetic S-palmitoylated peptides (a) with lactoglobulin derived peptides (b). The mix of peptides (c) shows both lactoglobulin peptides (*) and synthetic Palm/÷Palm peptides. S-palmitoylated peptides are retained in the SDC pellet as shown from precipitation and washing supernatants (d-e). (f) efficient release of de-palmitoylated peptides from the SDC fraction using DTT. **D**) Schematic representation of the SDC-ACE workflow using SDC acid precipitation for efficient S-palmitoylated peptide enrichment. Schematic representation was created with BioRender.com.

The three peptides were subsequently incubated with common reagents used for reduction of reversibly modified cysteines (e.g., disulfide bonds) and thioesters; Tris(2-carboxyethyl)phosphine (TCEP), DTT and HA, to test their efficiency in releasing the S-palmitoylation. Traditionally, HA has been used for decades for the selective release of palmitoyl groups from cysteines [20], however with very little evidence for its selectivity. Efficient and selective enrichment of formerly depalmitoylated peptides using the ABE and Acyl-RAC methods require that HA is very selective in releasing the palmitoyl group as these methods subsequently link biotin or any other tag to newly generated free SH group followed by enrichment and very sensitive LC-MS/MS analysis. However, the ability of HA to reduce abundant disulfide bonds has to our knowledge never been assessed. We therefore tested the HA specificity on bovine serum albumin, where the free cysteines were blocked with N-ethylmaleimide prior to precipitation of the protein. After tryptic digestion, we assessed the reduction of the disulfide bonds left in the protein using DTT, TCEP and various concentrations of HA. From the result of the experiment (**Supplementary fig. S3**), it is evident that higher concentrations of HA, as the ones used in most ABE and Acyl-RAC experiments, leads to a reduction of a small amount of disulfide bonds in BSA, resulting in subsequent contamination of formerly S-palmitoylated peptides with abundant disulfide-bonded cysteines in previous studies using the conventional methods for identifying de-palmitoylated peptides.

To test the reducing reagents’ ability to reduce the thioester bond, the synthetic S-palmitoylated peptides were incubated with different concentrations of HA, DTT and TCEP for 60 min and de-palmitoylated peptides were quantified using LC-MS/MS (**Fig. 1B**). Surprisingly, HA (100 mM) resulted in a small increase in peptide de- palmitoylation compared to 750 mM or 1000 mM. DTT was the most efficient of the three reducing reagents and we therefore chose 20 mM DTT as the optimum DTT concentration for release of de-palmitoylation peptides from the SDC pellet. In addition, this concentration shows efficient de-palmitoylation and does not require high concentration of subsequent alkylating reagent (**Supplementary fig. S4**). For reduction of disulfide bonds in our method we chose 1-3 mM TCEP as this resulted in minimal release of S-palmitoylation.

The specificity of the SDC enrichment of the S-palmitoylation peptides in a background of peptides derived from bovine lactoglobulin was examined by MALDI-MS analysis (**Fig. 1Ca-c**). After adding SDC to a final concentration of 1% followed by acidic precipitation and centrifugation, the supernatant (S1) was analyzed and signals originating from the S-palmitoylated peptides were absent, suggesting their precipitation with the SDC (**Fig. 1Cd**). Washing the precipitated SDC pellet only recovered remnant peptides from lactoglobulin, as expected (**Fig. 1Ce**). De-palmitoylation using 20 mM DTT on the re-solubilized SDC pellet resulted in the release of all three de-palmitoylated peptides (**Fig. 1Cf**), illustrating high selectivity of the method.

### Large scale SDC-ACE strategy

We designed the optimized SDC-ACE strategy for enriching formerly S-palmitoylated peptides using SDC acid precipitation to be applied to cells and tissues (**Fig. 1D**). Proteins are first extracted using 3-10% SDC (optimal analysis is obtained using a crude membrane preparation [32]), reduced using 1-3 mM TCEP which is adequate for most applications and alkylated using 10 mM N-ethylmaleimide. The proteins are then proteolyzed using trypsin or Lys-C/trypsin. After proteolytic digestion, the peptide solution is subsequently acidified and SDC pelleted by centrifugation. The SDC pellet is subsequently washed 3 times with washing buffer and twice with H_2_O, all buffered to pH 8-8,5 using NaOH. Finally, reduction of S-palmitoylation is performed using 20 mM DTT in 50 mM HEPES, pH 8 and subsequent alkylation of de-palmitoylated cysteines using Iodoacetamide. De-palmitoylated peptides are collected from the supernatant following SDC acid precipitation and centrifugation and they are subsequently analyzed by 1D or 2D LC-MSMS after desalting. A small volume of SDC sample without DTT can serve as a LC-MS/MS background control for the de- palmitoylation step.

### The mouse brain S-palmitoylome

The SDC-ACE method was applied to a crude membrane preparation from mouse brain in replicates using a total of 2 mg protein as starting material per replicate, which is comparable with other studies assessing formerly S-palmitoylation sites in cells or tissues. The enriched de-palmitoylated peptides were separated into 12 concatenated fractions using high pH reversed-phase separation. From the two replicates a total of 36,125 unique cysteine-containing peptides (**Supplementary table S1-3**) were identified from 6,721 proteins (**Fig. 2A**). A small aliquot of the SDC pellet analyzed with and without DTT revealed a “false discovery rate” in isolation of cysteine-containing peptides estimated to be below 0.49% (**Supplementary fig. S5**). A total of 81.6% of the de-palmitoylated peptides contained one cysteine, whereas 18.4% of the de-palmitoylated peptides contained two or more closely spaced cysteines, generating highly hydrophobic hubs on protein surfaces likely for interaction with lipid membranes, proteins, or other biomolecules (**Fig. 2A**).

**Figure 2:**
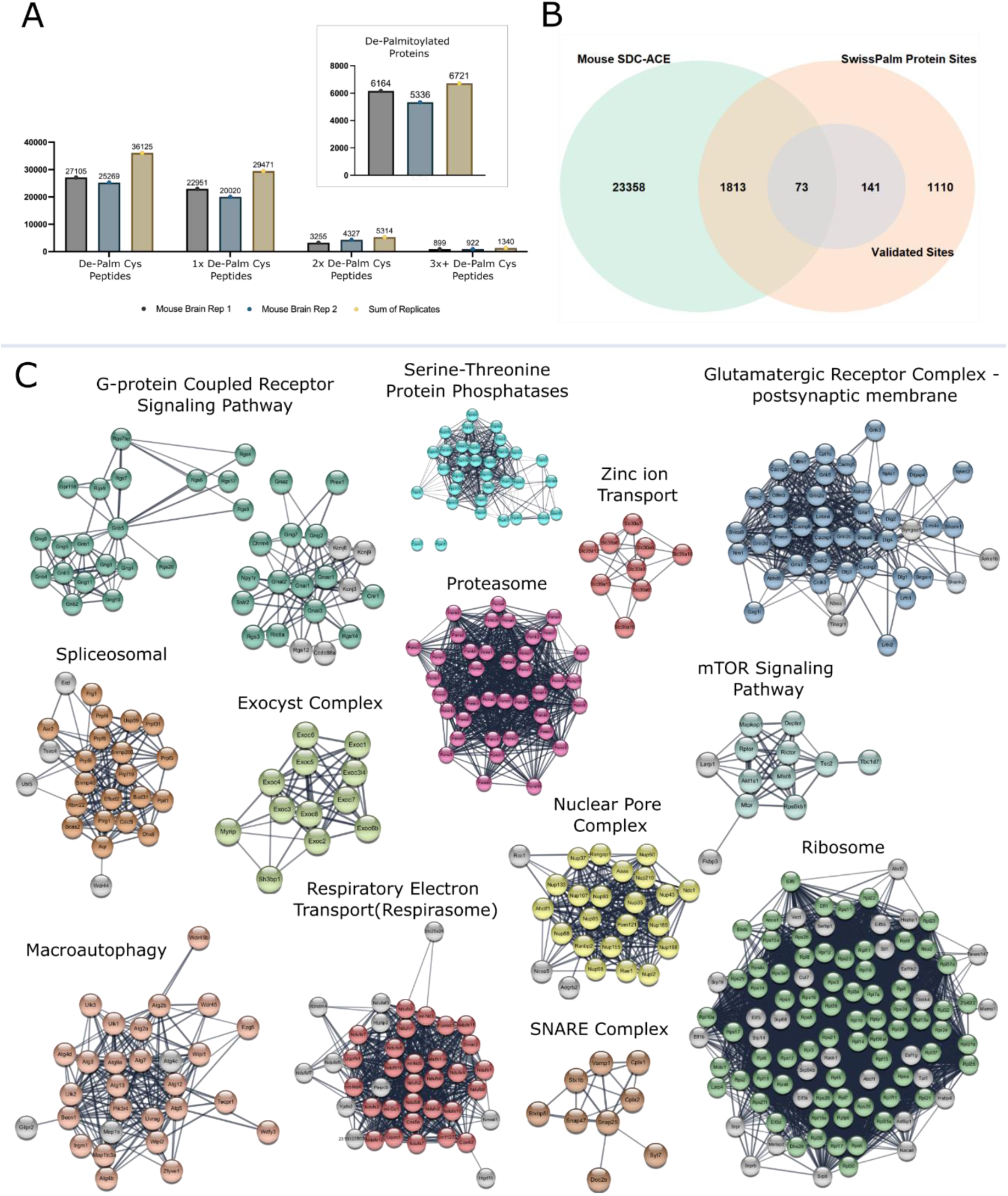
**The S-palmitoylation from mouse brains. A**) The number of identified de-palmitoylated peptides and proteins in each of the two replicates and the sum of both. Peptides are separated based on number of S-palmitoylation modification sites in a tryptic peptide, ranging from one to 3+. **B**) Comparison of mouse brain S-palmitoylation sites identified by the SDC-ACE method with all unique S-palmitoylated sites, including experimental validated unique sites in the SwissPalm database. **C**) Enrichment of GO and KEGG-pathway terms of the SDC-ACE enriched mouse brain S-palmitoylation proteins using the Cytoscape StringApp. Multiple protein complexes are illustrated.

As also noticed by Collins et al [24], we observed that some of the identified S-palmitoylation sites in our data mapped to cysteines annotated as situated in disulfide bonds (experimentally determined and predicted) in the Uniprot database. S-palmitoylation and disulfide bonds are not mutually exclusive and therefore both reversible cysteine modifications could occupy the same sites. We have previously observed that SDC is capable of co-precipitating proteins (data unpublished) and therefore larger disulfide-linked peptides could potentially pellet with the SDC. Therefore, to increase the confidence in our list of de-palmitoylated peptides, we filtered out all identified peptides with cysteine sites annotated as situated in disulfide bonds according to the UniProt annotation. This included multi-de-palmitoylated peptides even if only a single cysteine was annotated in the UniProt disulfide database. We found that 2,794 sites from our mouse brain data mapped to cysteines annotated as disulfide bond forming cysteines in the UniProt database, corresponding to 9.9% of all identified sites (28,256 sites in total) in the mouse brain data. A total of 128 non-disulfide cysteine sites were filtered out, as they were located on multi-de-palmitoylated peptides with at least a single disulfide annotated cysteine. This result is close to the 6.5 % that was observed by Colins et al. [24]. However, It is worth noting that S-palmitoylation has previously been reported to occur on cysteines involved in disulfide bond formation [33]. Thus, these sites may very well be S-palmitoylated in our samples.

Our list of mouse brain de-palmitoylated peptides share 1,886 of the 3,137 unique S-palmitoylation sites from predictions, compiled palmitoyl-proteome and validation studies in the mouse SwissPalm database [34, 35] (60.12%), including 73 of the only 214 validated unique sites in the database (**Fig. 2B**). Sites from SwissPalm were filtered for only SwissProt curated proteins. Some of the validated sites were not identified due to the tryptic cleavage specificity generating less likely detectable peptides in the LC-MS/MS analysis and UniProt disulfide annotation. Considering the use of HA in previous studies for “selective” reduction, and the difficulty in predicting S-palmitoylation by bioinformatics tools, we assume that a significant number of false de- palmitoylation sites is present in the SwissPalm database, contributing to a reduced overlap between sites in our dataset.

Gene Ontology (GO) and Kyoto Encyclopedia of Genes and Genomes (KEGG) pathway analysis of proteins containing peptides with S-palmitoylation sites was highly associated with known larger protein complexes, such as the ribosome, spliceosome, proteosome, SNARE-complex, Exocyst complex, mTOR-complex 1, Complex I-V and ATPase Complexes (mitochondrial respiratory system, respirasome) (**Fig 2C**). S-palmitoylation has previously been associated with some of these complexes [4, 34, 36], however, the present results strongly suggest that S-palmitoylation may be of much broader importance for formation, interaction and regulation of protein complexes and protein-protein interaction in the cell.

A total of 1,369 peptides from 768 proteins, resulting in 1,340 peptides from 756 proteins after disulfide correction contained 3 or more cysteines located in the same tryptic peptide sequence, indicating the presence of hydrophobic clusters in short regions in these proteins (**Fig. 2A**). Many well-known multi-S-palmitoylated proteins were identified, such as SNAP25a/b (annotated spectra for the de-palmitoylated peptides in **Supplementary fig. S6A** and **S6B**), involved in the SNARE complex involved in synaptic vesicle exocytosis and PSD-95, a postsynaptic scaffolding protein required for synaptic plasticity [1]. A peptide from the N-terminal, extracellular part of Serine incorporator 5 (SERC5)(Uniprot: Q8BHJ6), an important protein in viral infection and neurological disease [37], was identified containing 10 de-palmitoylated sites in a single tryptic peptide (**Supplementary fig. S7**). Furthermore, S-palmitoylation sites were identified on multiple enzymes involved in regulatory processes in the cells including E3 ubiquitin-protein ligases, protein kinases, protein phosphatases, acetyl-transferases, deacetylase, GTPases and surface receptors, strongly suggesting that this modification may be considerably more involved in regulatory processes in the cell than previously believed.

More than 350 kinases were identified in our dataset with potential S-palmitoylation sites, including small molecule kinases and protein kinases. These include well characterized important kinases in health and disease, such as ribosomal protein S6 kinases, Mitogen-activated protein kinases, Glycogen synthase kinase-3A/B, Cyclin-dependent kinase 5, Calcium/calmodulin-dependent protein kinases and various tyrosine kinases such as epidermal growth factor receptor, Tyrosine-protein kinase Fyn/Lyn/UFO/TYRO3/JAK1/JAK2/ Fer/CSK/ALB2/FLT3 and many others. Some of these protein kinases were potentially heavily S-palmitoylated. For example, we identify a total of 21 de-palmitoylation sites in Protein Kinase C beta type, which is a kinase involved in several signaling pathways in the cell and is activated by calcium and diacylglycerol. This protein has two zinc finger domains (phorbol-ester/DAG-type 1 and 2) in the regions aa 36-86 and aa 101-151 [38]. However, we identify the cysteines in these regions to be S-palmitoylated in our data set (Cys 50, 53, 67, 70, 71, 78, 86, 115, 118, 132, 135, 143 and 151). The presence of these peptides in the SDC pellet due to conjugation with zinc is highly unlikely as these peptides were released by DTT reduction, which will not interfere in zinc conjugated peptides. To support this observation, we washed a fraction of a SDC pellet containing peptides from mouse brain membrane preparation with 10 mM EDTA in 50 mM HEPES, pH 8.5 for 30 min at 37 °C followed by acid precipitation. Subsequent LC-MSMS analysis of the released peptides did not result in the identification of cysteine-containing peptides (data not shown). The N-terminal domain of Protein Kinase C beta type, illustrated by a red circle in the 3D structure (**See Supplementary fig. S8**), contains several loops and disordered regions and heavy S-palmitoylation of this region could drive the localization of this kinase to the plasma membrane. Protein kinase C isoforms have previously been shown to be recruited to the plasma membrane after glucose-stimulation of pancreatic β-cells, but this translocation mechanism is unknown [39].

Phosphatases, responsible for dephosphorylation, were also found to be S-palmitoylated in our mouse brain dataset (e.g., Fig. 2C), including the abundant brain protein phosphatase calcineurin. Interestingly, several dual specificity protein phosphatases and tyrosine-protein phosphatases were found to be potentially modified by S-palmitoylation on the cysteine residue in the active site, in the C(X)5R motif [40, 41], a site that must be free for the enzyme to be active. This suggests that S-palmitoylation may shield dephosphorylation activity for these tyrosine phosphatases in the cell.

The S-palmitoylation of multiple enzymes responsible for the dynamics of reversible PTMs in cells strongly suggest a crosstalk between S-palmitoylation and other PTMs. Such a PTM crosstalk has previously been suggested for protein phosphorylation [42, 43], S-nitrosylation [44–46] and ubiquitination [47, 48].

### Palmitoylation of the mouse nuclear pore complex in the brain

The nuclear pore complex (NPC) is a critical structure that functions as a gateway between the nucleus and the cytoplasm in cells. It is composed of multiple proteins known as nucleoporins, which together form a large, octagonal complex[49]. This complex regulates the selective transport of molecules, such as RNA and proteins, in and out of the nucleus. In mice, as in other eukaryotes, the NPC is essential for maintaining cellular function and gene expression[49]. Structurally, it features a central channel surrounded by a ring of proteins, facilitating the bidirectional flow of cellular materials. Many of the proteins in the NPC are closely associated with the nuclear membrane[49].

Among the proteins of the nuclear pore complex [49, 50] S-palmitoylation sites were identified on 25 in our study (**Fig. 3**). These included proteins from the inner and outer rings and proteins situated in the nucleoplasmic basket, transmembrane ring, and the cytoplasmic filaments. This suggest that S-palmitoylation may be central for assembling and/or dynamic of the nuclear protein complex and therefore important for cytoplasmic - nuclear transportation of proteins and mRNA molecules.

**Figure 3:**
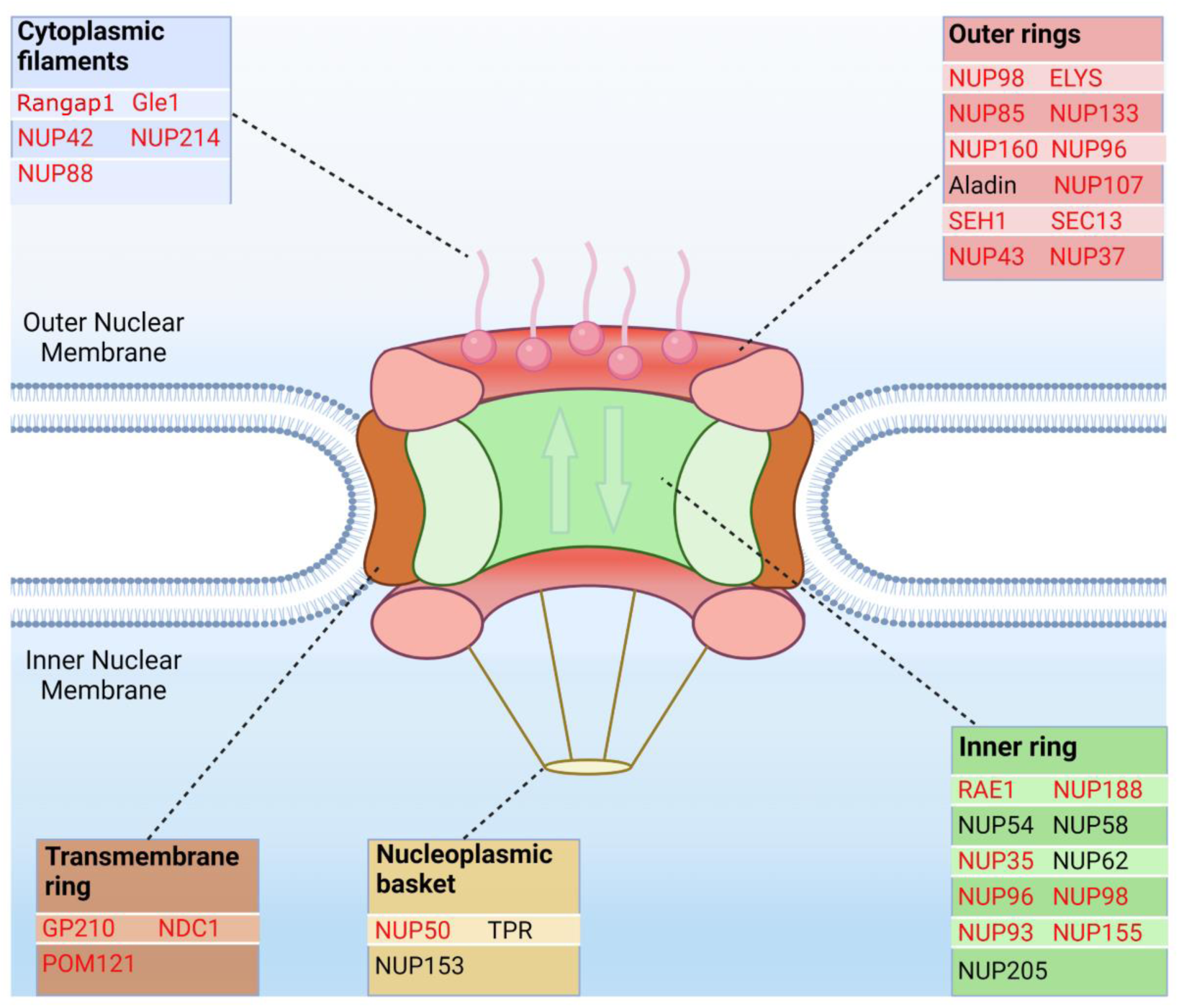
S-palmitoylated components of the nuclear pore complex identified from the mouse brain. Schematic illustration of the nuclear pore complex architecture that spans the nuclear envelope. Enriched de-palmitoylated peptides from the mouse brain could be identified from several proteins (highlighted in red) related to the nuclear pore complex. A total of 25 proteins known in the nuclear pore complex were identified as S-palmitoylated. These included proteins from the inner and outer rings and proteins situated in the nucleoplasmic basket, transmembrane ring and the cytoplasmic filaments. The figure was created with BioRender.com.

### Palmitoylation of palmitoyl-transferases in the mouse brain

As mentioned above, 24 zDHHC PATs exist in the mouse genome and several PATs are published to be S-palmitoylated by other PATs. A common feature of the zDHHC PATs is a transmembrane region and a cysteine rich domain (CRD-domain), that contain several down-stream cysteines and the catalytic DHHC cysteine, illustrated with the 3D structure for zDHHC9 obtained from Alphafold [51–53] in **Fig. 4A**. According to the 3D structures of all the zDHHC proteins, derived by Alphafold, the downstream cysteines are localized in sequential “stacks” close to the active site cysteine, as illustrated for zDHHC9 in **Fig. 4A-C**. Some of these cysteines have been suggested to participate in zinc binding as described for the zDHHC3 [15]. Within the CRD amino acid sequence downstream of the proposed active zDHHC site, several tryptic cleavages sites are located (K/R), and the presence of lipids in this area could compromise the tryptic cleavage specificity. Therefore, to investigate the S-palmitoylation of the CRD cysteines we searched the mouse brain de- palmitoylation data in a limited database containing only the zDHHC sequences using up to 4 missed cleavages in the database search criteria in Proteome Discoverer (PD). Since we blocked the free cysteines with NEM and labeled the formerly palmitoylated cysteines with iodoacetamide we used both modifications as variable modifications. The identified peptides were validated and filtered for 1% FDR using the percolator software [54] in PD. The database search result for zDHHC9 is shown in **Fig. 4B**. We identified all the 24 zDHHCs in the mouse brain palmitoylome. The result of the database search for the other zDHHCs can be seen in **Supplementary File “zDHHC 3D structures 1 and 2”**, where the sequence coverage from the database search is illustrated for each zDHHCs next to the 3D structures obtained from Alphafold. As evident from the peptide coverage, all the down-stream cysteines in all zDHHCs are potentially S-palmitoylated at some point in the mouse brain, but not fully S-palmitoylated, as illustrated by the differential presence of NEM on several cysteines. A cysteine located 6 amino acids downstream from the active site, conserved in all zDHHCs, was also found to be differentially S-palmitoylated in most of the zDHHCs in our data. Since we cannot calculate any stoichiometry from our data, we cannot speculate on the level of S-palmitoylation on these cysteines.

**Figure 4:**
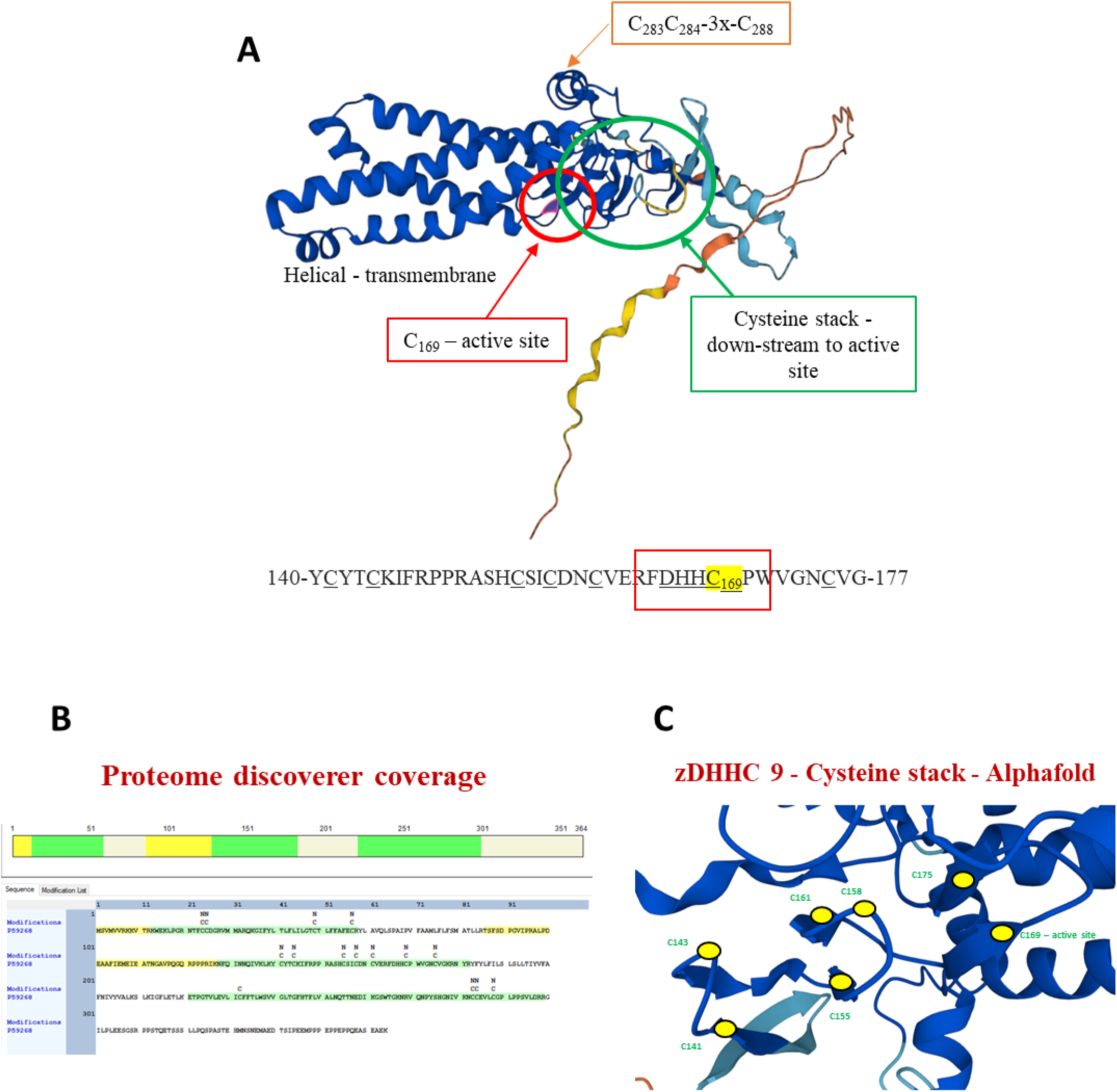
**3D structure and S-palmitoylation of zDHHC9. A**) Alphafold 3D structure of zDHHC9 highlighting the cysteine stack (green circle), the active site (red circle) and C-terminal S-palmitoylation. **B**) Sequence coverage of zDHHC9 from Proteome Discoverer, highlighting de-palmitoylated and free cysteines (de-palmitoylated cysteines - carbamidomethyl and free cysteines - ethylmaleimide). **C**) Alphafold 3D structure illustration of the “Cysteine stack”, the active cysteine site and the upstream cysteine (yellow dot illustrate the individual cysteines).

Since previous research have shown that the cysteines within the CRD could be conjugated to zinc [15], we, as described above, washed an SDC pellet containing peptides from mouse brain membrane preparation with EDTA and subsequent LC-MS/MS analysis of the released peptides did not result in the identification of cysteine-containing peptides, such as the ones covering the CRD cysteines. This suggest that the CRD cysteines are not conjugated to zinc in the SDC pellet. However, this does not rule out that a fraction of these cysteines could be conjugated to zinc at some point during the lifetime of cells.

These results strongly indicate that the conserved cysteines in the CRD and the up-stream cysteine are sub- stoichiometric S-palmitoylated. The presence of S-palmitoylated cysteines in very close proximity to the active site cysteine (illustrated in **Fig. 4C**) could have several reasons. The up- and downstream cysteines to the active site are very closely situated in the 3D structure (**Fig. 4C and supplementary File “zDHHC 3D structures”**) and therefore could serve as a reservoir of palmitate that can easily be transferred to the cysteine in the active site for S-palmitoylation of substrates. Alternatively, up- and downstream cysteines could be part of a larger conserved “active site pocket” where they all participate actively in the palmitoyl-transferase process, capable of transfer palmitate to the substrates. The latter is supported by the fact that mutations in the DHHC motif of yeast S-acyltransferases Swf1 and Pfa4 only partially abolish their activities [55].

### Mouse tissue specific S-palmitoylation

We further applied our method to investigate S-palmitoylation characteristics of different mouse tissues using TMT based quantitative proteomics in combination with SDC-ACE. After membrane enrichment, proteolysis, and TMT labeling, we enriched de-palmitoylated peptides from mouse brain, brown adipose tissue (BAT), white adipose tissue (WAT), heart, liver, and kidney. The enriched de-palmitoylated TMT labeled peptides were separated into 20 concatenated fractions using high pH reversed-phase separation, and each fraction was analyzed on LC-MS/MS. We identified a total of 18,575 de-palmitoylated peptides from 5,925 proteins across all tissues.

Removing cysteines potentially involved in disulfide bonds, yielded 16,823 de-palmitoylated peptides. As seen in the PCA analysis (**Fig. 5A**), the tissues are grouped into three separate clusters. Clustering de-palmitoylated peptides revealed distinct patterns for most tissues (**Fig. 5B**), reflecting the physiological differences between these tissues. Duplicate samples from each tissue grouped together similar in the PCA plot, collectively reassuring the specificity and reproducibility of our method. The mouse brain palmitoylome is somehow significantly different from the other tissues, most likely due to the very special functions of neurons as rapidly excitable cells, requiring abundant intracellular transport of vesicles and secretion of proteins and small molecules, processes known to be regulated by S-palmitoylation [56–58]. Many of the unique S-palmitoylated proteins in the mouse brain are, as stated above, present in the nerve-terminals and axons where multiple transportation and secretion mechanisms exist. The kidney and liver group close to each other in the PCA plot, which is driven by S-palmitoylation of common metabolic pathways in these tissues.

**Figure 5:**
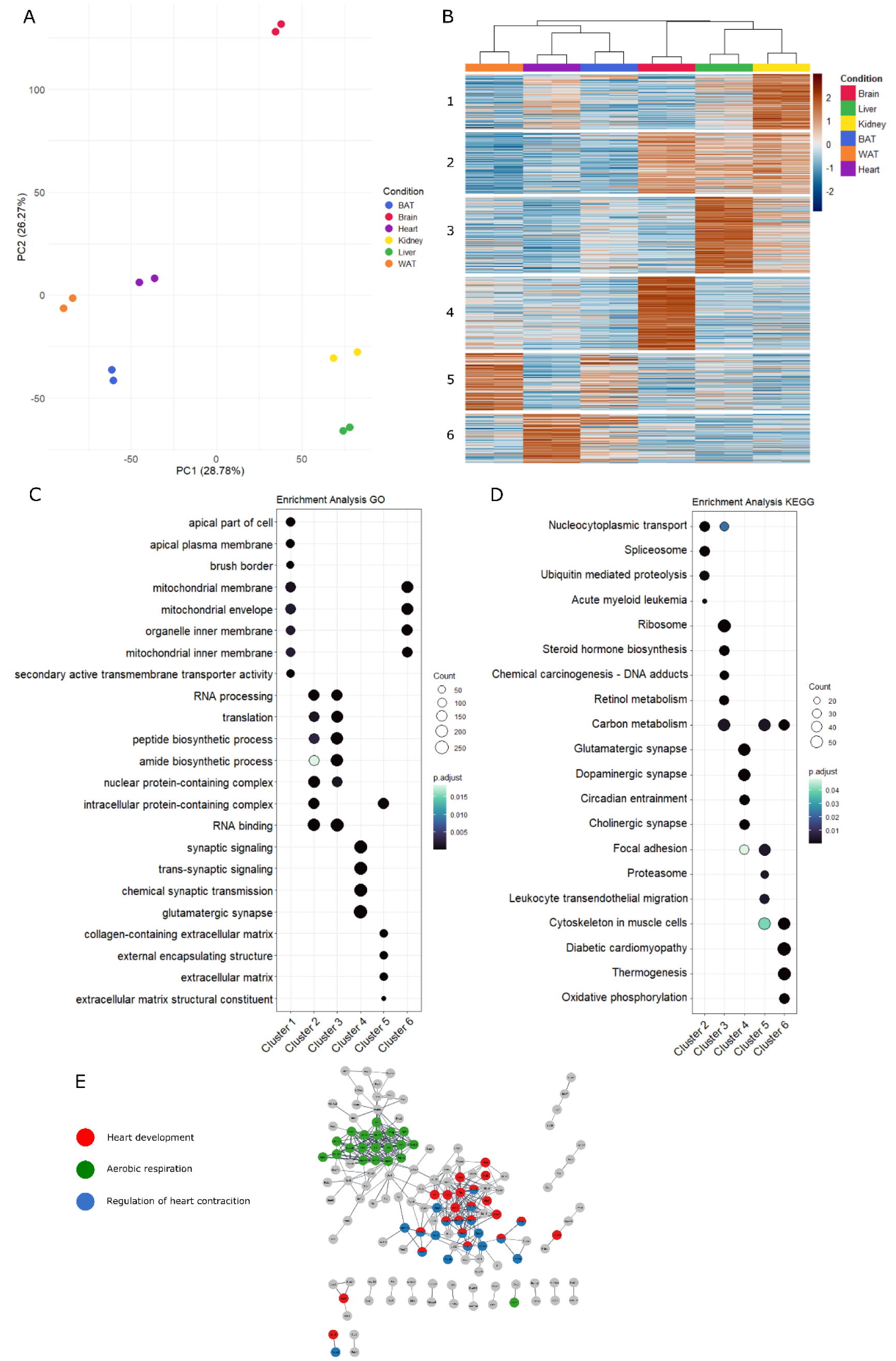
**Characterization of mouse tissue palmitoylomes. A**) PCA analysis of mouse organ tissue palmitoylation. **B**) Heatmap of kmeans clustered mouse organ tissues. Each organ includes 2 replicates. Scale bar refers to scaled log_2_ abundances for each replicate. Cluster numbers are listed on the left of the heatmap. **C + D**) GO/KEGG enrichment analysis of clustered mouse tissue. E) StringApp enrichment of GO/KEGG terms for mouse heart tissue following protein filtering for upregulated features with a log_2_ value above 2 in at least one replicate across all conditions.

GO-term and KEGG pathway enrichment of cluster 4 revealed terms mainly associated with the brain, such as synaptic signaling/transmission, agreeing with the increased enrichment of these de-palmitoylated peptides in the brain tissue and previous experiments on mouse brains [24] (**Fig. 5C and D**). Similarly, in cluster 3, we enriched terms related to metabolism, such as amide/peptide biosynthetic processes and proteins involved in bile metabolism, showing a predominant association with the liver (**See Supplementary fig. S9**), and we observed substantial S-palmitoylation on key proteins involved in small molecule transporters and ion channels in cluster 1, associated with the kidney. Cluster 6 represented S-palmitoylation abundant in the heart revealed GO/KEGG pathways such as heart development, regulation of heart contraction and aerobic respiration (**Fig. 5E**). The impact of S-palmitoylation on cardiac physiology is poorly understood with only few known S-palmitoylated proteins with known roles in the heart (reviewed in[59]). We identified several potential modification sites on Ryanodine receptor 2 (E9Q401) and S-palmitoylation of cysteine 1189 in the Voltage-dependent L-type calcium channel subunit Alpha 1C (Q01815), proteins heavily involved in optimal myocardial function. These data emphasize the extensive role of S-palmitoylation in cellular processes across the entire body. The StringApp network GO/KEGG analyses for liver, brain and kidney can be seen in **Supplementary Figures S9-S11**.

### Palmitoylation in mouse brain tissue from obesity linked type 2 diabetes (T2D) in mice

We have previously profiled the proteome and phosphoproteome in relation to pancreatic β-cell adaptive restoration of insulin secretory capacity in obese diabetic db/db mouse islets, in which key molecular events in the secretory pathway reverted to normal upon euglycemic treatment [60]. Considering that S-palmitoylation is heavily related to trafficking/sorting events within the cell, we decided to apply our method to study the S-palmitoylome in brains from db/db mice. The db/db diabetic mouse model carries a mutation in the leptin receptor gene, inactivating the receptor and leading to hyperphagia, severe obesity, hyperglycemia and glucose intolerance [61]. The model is widely applied for studying early phenotypes of obesity linked T2D. The appetite-suppressing and weight-reducing effects of leptin is believed to be exerted primarily in the hypothalamus. Leptin receptor signals through a multitude of signaling pathways, including JAK2-STAT3, PI3K-AKT-mTOR, SHP2-ERK and more (reviewed in [62]). Out of the 15,421 potential S-palmitoylated peptides identified in the analysis, we identified 198 significantly regulated S-palmitoylated peptides (**Fig. 6A**). These sites were found in proteins such as Tyrosine-protein phosphatase non receptor type 11 (Ptpn11, SHP-2), WD repeat and FYVE domain-containing protein 1 (WDFY1), Calcium-activated potassium channel subunit alpha-1 (Kcnma1), PI3K regulatory subunit beta (p85β) and protein kinase C delta type (PKC-δ) (**Fig. 6B**), proteins associated with signaling pathways of both insulin and leptin. WDFY1 belong to a large family of WD40 repeat domain containing proteins, one of the most common protein interaction domains in the human proteome [63]. This domain consists of a central pore formed by circularly arranged beta-propeller folds, often acting as interaction scaffold proteins within multiprotein complexes [63]. All identified S-palmitoylated cysteines in our analysis are localized to the WD40 domain (**Fig. 6C**). WDFY1 has been reported to localize to endosomes through its FYVE domain and act as an adaptor molecule in the Toll-like receptor (TLR) activation of NF-κB by TLR3/TLR4 signaling [64]. The role of S-palmitoylation of WDFY1 in diabetic db/db mice is unknown.

**Figure 6:**
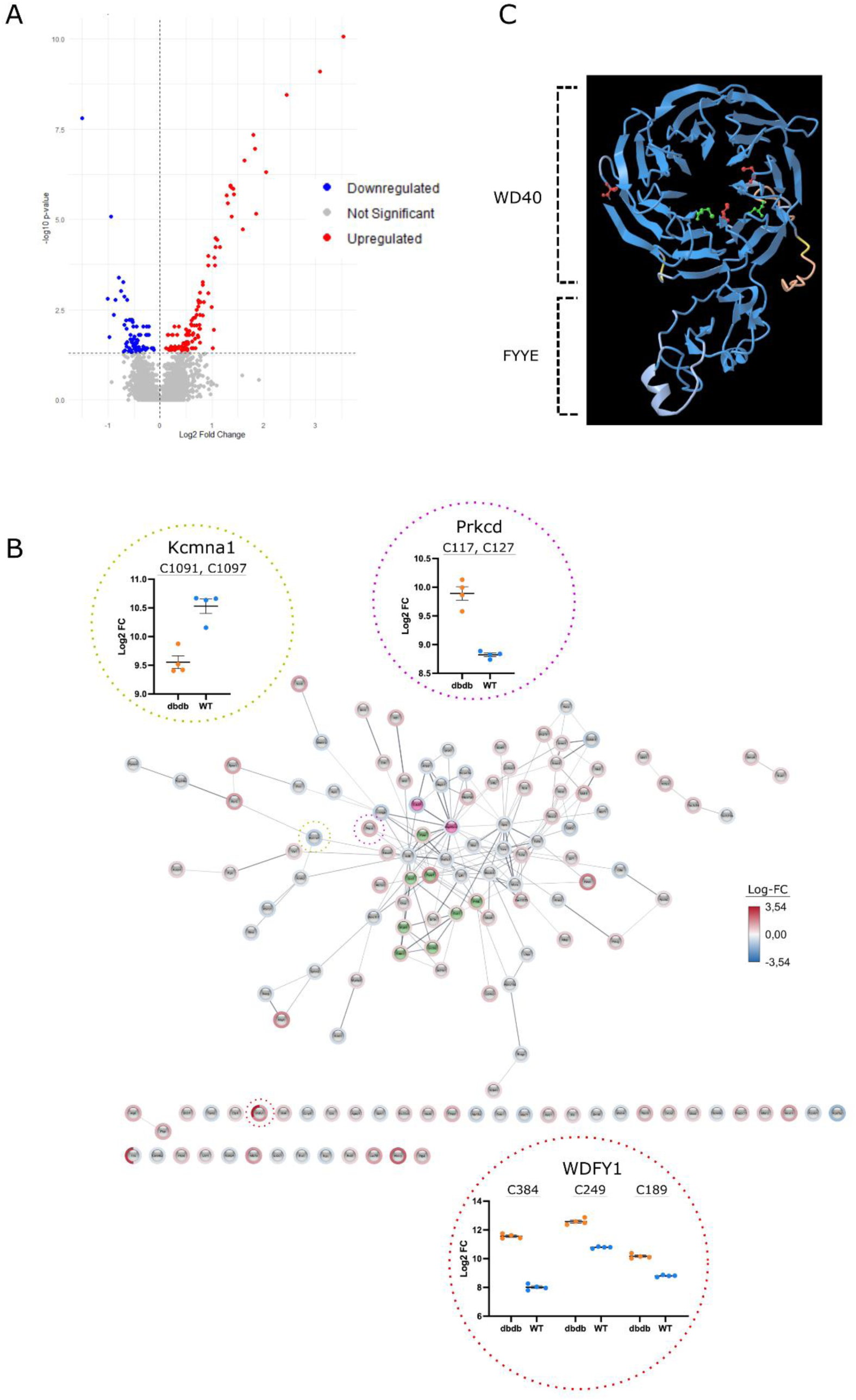
**Characterization of db/db mouse brain palmitoylome. A**) Volcano plot of differentially regulated S-palmitoylated peptides enriched from Wildtype (WT) and db/db mouse brains. Significantly upregulated and downregulated S-palmitoylated peptides are shown in red and blue, respectively. Non-significantly regulated peptides are shown in grey. **B**) StringApp protein-protein interaction enrichment of proteins with identified significantly regulated S-palmitoylated peptides in the db/db mouse brain. Scatter diagram of selected protein/peptides show relative abundance between identified peptide sites of WT and db/db mice. Scatter diagram of selected protein/peptides is highlighted in the StringApp enrichment network by colorized circles, red circles: WDFY1, yellow circle: Kcnma1 and purple circle: Prkcd. **C**) Graphical illustration of WDFY1 showing the WD40 repeat domain and FYYE domain. Significantly regulated palmitoylation sites are illustrated and colorized in red. Non-significantly regulated palmitoylation sites identified in our experiment is shown and colorized in green.

Activation of hypothalamic PKC-δ by lipid modification has previously been reported to lower hepatic glucose production and regulate insulin-sensitivity by glucagon-like peptide 1 (GLP-1) [65, 66]. In addition, PKC-δ^+^ neurons in the central amygdala have been implicated in regulating food intake. Interestingly, a recent paper found PKC-δ S-palmitoylated in hypothalamic microglia under high-fat diet-induced microglial neuroinflammation [67]. Here, S-palmitoylation of PKC-δ was suggested to induce PKC-δ activation by facilitating T505 phosphorylation, leading to subsequent downstream signaling and contributing to microglial activation. Inhibiting PKC-δ S-palmitoylation using a derivative of artemisinin, artemether, suppressed neuroinflammation and alleviated hepatic lipid metabolism disorder in mouse model [67]. We found PKC-δ significantly upregulation on the doubly S-palmitoylated peptide carrying Cys117 and Cys127 in db/db mice (FDR<0.05, Log_2_-FC: 1.07), located just upstream of the two zinc finger domains, as mentioned previously, and pseudo-substrate domain of novel PKC isoforms. The zinc finger domains, also known as C1a and C1b, is known as the binding site of novel PKCs to diacylglycerol (DAG) at the plasma membrane, a key step in the downstream signaling of novel PKCs[68]. In addition, zDHHC5, a PAT mainly localized at the plasma membrane, were proposed as the PAT for PKC-δ S-palmitoylation [67]. This indicates that PKC-δ is S-palmitoylated at the plasma membrane, potentially after recruitment and interaction with plasma membrane DAG, facilitating stable interaction with the plasma membrane or conformational changes for T505 phosphorylation.

## Conclusion

The new SDC-ACE method is an efficient and easy method for identification of S-palmitoylation sites. It is associated with fewer false identifications due to the efficient enrichment of the S-palmitoylated peptides with SDC prior to de-palmitoylation and the elimination of the use of hydroxylamine, which was here found to reduce abundant disulfide bonds at high concentrations resulting in false identifications. Applying our method to biological samples, enabled us to characterize in depth the mouse brain palmitoylome, map tissue-specific S-palmitoylation differences between mouse organs and characterize the db/db mouse brain palmitoylome. It is realized that this novel SDC-ACE method has revealed a first-insight into S-palmitoylation PTMs, and the further validation studies will be needed to gain deeper mechanistic insight. Nonetheless, we highlight palmitoylation as an extensive PTM involved in many regulative processes and associate palmitoylation with several tissue-specific pathways, implying S-palmitoylation could well be a critical PTM for cellular processes and signaling pathways in biological systems. Moreover, identifying palmitoylation on several important cellular enzymes responsible for regulation of other PTMs, including kinases and phosphatases, indicates a coordination of distinct PTMs that may specifically localize signal transduction activity to particular compartments of the cell. Finally, in observing that PKC-delta S-palmitoylation as significantly upregulated in the brain of obese diabetic mice, may indicate altered neuronal signaling pathways that potentially contribute to disease pathogenesis in obesity-linked T2D.

## Supporting information

Supplementary zDHHC 3D structures

## Acknowledgements

We would like to thank Lene Andrup Jakobsen and Arkadiusz Miroslaw Nawrocki for excellent technical support and Sofie Blomberg Elmkvist and Andreas Abildskov Thomsen for discussions on data analysis at the Protein Research group of University of Southern Denmark. This research was financially supported by the Novo Nordic Foundation (No. 95-310-73303-01100), the Lundbeck foundation (R336-2020-1113) and the Villum Center for Bioanalytical Sciences at SDU. GP is supported by the São Paulo Research Foundation (FAPESP), grants processes n° 2018/18257-1 (GP), 2018/15549-1 (GP), 2020/04923-0 (GP) and Capes “bolsa de produtividade”.

## Methods

### Materials

Sodium deoxycholate and Dithiothreitol was obtained from Sigma Aldrich, Germany. Trifluoroacetic acid, Hydroxylamine (HA) and Formic acid was obtained from Merck, Germany. Tris(2-carboxyethyl)phosphine (TCEP) was obtained from Thermo Fisher Scientific, Germany.

### Animals and harvesting of tissues

Mice of strain NMR1 were euthanized at age 3 months and whole brains were excised rapidly, flushed in 0.1% saline, and snap frozen in liquid nitrogen. The various organs (liver, kidney, heart, brown fat and white fat) were dissected and snap frozen in liquid nitrogen. All organs were stored at -80 degrees until use. All animal work was performed in accordance with the guidelines of the University of Southern Denmark.

Wild type (WT) C57BL/6J and C57BL/6J^db/db^ mice were bred and purchased from The Jackson Laboratory (Bar Harbor, ME) and housed on a 12-hour light/dark cycle with free access to standard mouse food and water. Mice were sacrificed between 14-16 weeks of age and brains where isolated immediately after and stored at - 80°C until further processing.

### Synthetic peptides

The three tryptic synthetic peptides used for this study (See Table 1) contains a single free cysteine (highlighted in bold in the table below) and no serine and threonine residues, eliminating the possibility of S-palmitoylation of these residues using the palmitoylation method. All other cysteines were carbamidomethylated. The C-terminal arginine’s were all heavy labeled with Arg-10 ((^13^C6,^15^N4). The synthetic peptides were obtained as a gift from Prof. Phillip Robinson, Childrens Medical Research Institute, Westmead, Sydney, NSW, Australia.

**Table 1:** Synthetic peptides used for developing the SDC-ACE method. * denote carbamidomethylated cysteines.

| ID | Sequence |
| --- | --- |
| Peptide A | VAIHCHAGLGR |
| Peptide B | EEPGC*C*IAVHCVAGLGR |
| Peptide C | EEPGC*C*VAVHCVAGLGR |

### Generation of palmitoylated synthetic peptides

Synthetic S-palmitoylated peptides were generated essentially as described in [31]. Synthetic peptides were S-palmitoylated by dissolving the dried peptides in 10µL 100% trifluoroacetic acid (TFA) and subsequently adding 1µL palmitoyl chloride (Sigma Aldrich, Steinheim, Germany). The solution was incubated for 10 minutes at room temperature (RT). After incubation, the solution was diluted to 5% TFA and 20% Acetonitrile (ACN) and centrifuged at 14,000g for 10 minutes. S-palmitoylated peptides were purified by mixing the solution with Oligo R1 material (Poros®, Perseptive Biosystems, Framingham, USA) for 10 minutes at RT. After incubation, the beads were washed with 20% ACN, 0.1% TFA and the S-palmitoylated peptides were eluted with 60% ACN.

### MALDI MS of S-palmitoylated synthetic peptides

A small aliquot of the S-palmitoylated and non-palmitoylated synthetic peptides (0.5 µL), respectively, were mixed with 0.5µL 0.1% TFA and 0.5µL α-Cyano-4-hydroxycinnamic acid (4-HCCA) on a stainless steel MALDI target (Bruker) and the mixture was allowed to dry at room temperature. The peptides were analyzed using a Bruker Ultraflex (Bruker Daltonics, Germany) in positive reflector ion-mode. An average of 4000 laser shots were used for each peptide to obtain a MALDI MS spectrum. For the mixture of S-palmitoylated synthetic peptides and tryptic peptides from lactoglobulin the same procedure was used, except for the solution containing the wash of the SDC pellet. Here, the supernatant from the wash was concentrated on a Poros Oligo R2 microcolumn and the peptides were eluted directly onto the MALDI MS target using the MALDI MS matrix solution.

### Comparing de-palmitoylation of synthetic S-palmitoylated peptides using HA, DTT and TCEP

For comparing the de-palmitoylation efficiency of HA, DTT and TCEP, S-palmitoylated synthetic peptides were mixed and diluted in 50mM Tris containing 1% SDC, pH: 7.4. The pool of peptide solution was split into new tubes with equal volumes. Samples were then treated with either HA, DTT or TCEP at concentrations given in Figure 1B and incubated at RT for 1 hour. Following incubation, SDC was precipitated by adding 100% formic acid to a final concentration of 2% and the samples were centrifuged at 14,000g for 10 minutes to pellet the SDC.

Released de-palmitoylated peptides were then purified using Poros Oligo R3 reversed-phase (RP) microcolumns, by loading the acidified solution in equal volume to an equilibrated p200 stage tip containing C18 3M material disk (Fisher Scientific, US) and Oligo R3 material (Poros, Perseptive Biosystems, Framingham, US). Microcolumns were washed with 40 µL 0.1% TFA and peptides were eluted with 40 µL 60% ACN in 0.1% TFA. Eluted samples were dried by lyophilization prior to LC-MS/MS.

### Label-free quantification of de-palmitoylated synthetic peptides using LC-MS/MS

The purified de-palmitoylated samples were analyzed with a nano-Easy liquid Chromatography instrument (Thermo Fisher Scientific) coupled to a Q-Exactive HF mass spectrometer (Thermo Fisher Scientific). Dried samples were resuspended in 0.1% formic acid and loaded onto a 2cm 100 ID pre-column containing C18 material (Reprosil 3 µm, Dr. Maisch, Ammerbuch-Entringen, Germany) and eluted directly onto an analytical column with a gradient of 0% to 34% buffer B (90% ACN in 0.1% FA) over 23 minutes. All LC-MS/MS runs were performed with an analytical column of 18cm x 75µm inner diameter fused silica packed with C18 Reprosil 3 µm material (Dr. Maisch, Ammerbuch-Entringen, Germany). Mass spectrometry was performed using full MS scan with a resolution of 120,000 and a target value of 3 x 10^6^. For MS/MS maximum injection time was 100 ms and scan range 300-1300 m/z. Synthetic peptides were quantified based on the extracted ion chromatogram (XIC) intensity levels from raw data files.

### Enrichment of de-palmitoylated peptides from mouse brain membrane preparation

A brain from a C57BL/6J mouse (day 21 old) was homogenized using a Dounce homogenizer in 2 mL ice-cold 100 mM Sodium Carbonate (Na_2_CO_3_) for isolation of membrane proteins [69]. After homogenization, the solution was subjected to probe sonication for 3x20 sec. at 60% amplitude on ice, using a Q125 Sonicator (Qsonia). After sonication, the sample was incubated at 4°C with rotation for 1 h. After incubation, the sample was ultracentrifuge at 120.000 g for 60 min at 4°C. After centrifugation, the supernatant was transferred to another tube and stored. The pellet containing membrane proteins, membrane associated proteins and larger protein complexes were washed once with 50 mM HEPES, pH 8. The pellet was solubilized by sonication for 2x60 sec in 500 µL 50 mM HEPES, pH 8, containing 10% SDC and subsequently placed at 110 °C for complete inactivation of enzymes. After heat denaturation, the solution was diluted to 5% SDC and the sample was centrifuged at 20.000g for 20 min. The supernatant was transferred to a low binding Eppendorf tube (Sorensen Bioscience Inc. 1,7 mL) and the protein concentration was measured using an Implen NanoPhotometer N60 (Implen, Germany).

A total of 2 mg of the membrane enriched protein fraction was subjected to reduction of disulfide bridges using 3 mM TCEP for 30 min at RT and the free cysteines were subsequently alkylated using 10 mM N-Ethylmaleimide (NEM). After alkylation, the proteins were digested using Endoproteinase Lys-C (0.04AU) for 2 hours at 37 °C. After incubation, trypsin (homemade modified trypsin[70] (5% w/w)) was added and the sample was left overnight at 37 °C. After incubation, formic acid was added to a final of 2% and the sample was centrifuged at 20.000g for 20 min to pellet the SDC. After centrifugation, the supernatant was transferred to another low binding Eppendorf tube and the pellet was washed 3 times using 20% Acetonitrile, 50 mM HEPES and 1 M KCL. Briefly, the pellet was solubilized by sonication (1x20 sec at 40% amplitude) and NaOH was added to increase the pH to >8.0 for complete solubilization of the SDC. After incubation, the SDC was precipitated using formic acid (final of 2%) and centrifugation at 20.000g for 20 min. After the three washes, the SDC pellet was washed two times with H_2_O buffered to pH 8 by NaOH to solubilize the SDC in a similar way as described above. All washes were dried by lyophilization. After the last wash, the SDC pellet was solubilized in 600 µL 50 mM HEPES and adjusted to pH 8 using NaOH. An aliquot (20 µL) was transferred to another low binding Eppendorf tube and the remaining solution was added DTT to a final concentration of 20 mM. An aliquot (20 µL) was transferred to a low binding Eppendorf tube. All three samples were incubated at RT for 60 min. After incubation, formic acid was added to the two 20 µL aliquots to a final of 2% and the samples were centrifuged at 20.000g for 20 min to pellet the SDC. These two samples were subsequently analyzed by LC-MS/MS for illustrating the efficiency of releasing de-palmitoylated peptides from SDC. The third sample containing the remaining 560 µL de-palmitoylated sample was alkylated using iodoacetamide (40 mM) for 45 min at RT in the dark. After alkylation, formic acid was added to a final of 2% and the sample was centrifuged at 20.000g for 20 min to pellet the SDC. After centrifugation, the supernatant was subjected to Poros Oligo R3 RP microcolumn desalting and the peptides were subsequently lyophilized for high pH RP separation. The procedure was repeated twice to produce a technical duplicate.

### Analysis of de-palmitoylated peptides from mouse tissue using TMTpro quantitation

Membrane and membrane-associated proteins were isolated from mouse tissues (2 replicates for each tissue) using bead beating in 1 mL 100 mM ice-cold Na_2_CO_3_. The bead beating was done using a FastPrep-24 instrument (MP Biomedicals) with a total of 25 ceramic beads per sample in a 2 mL Eppendorf tube. After bead beating, the sample was transferred to a 1.5 mL Eppendorf tube and sonicated for 2x20 sec at 60 % amplitude on ice using a Q125 probe-sonicator (QSONICA). After sonication, the sample was centrifuged for 10 min at 14000 g and the supernatant was transferred to a Microtube WX 1.5 mL ultracentrifuge tube (Thermo Scientific Cat. No. 314352H01), incubated for 60 min a 4°C with rotation and subsequently the sample was centrifuged at 120.000g for 60 min at 4°C using a HITACHI micro ultracentrifuge CS 150NX. After centrifugation, the supernatant was removed and the pellet was washed with 100 mM HEPES, pH 8. The wash was discarded. The membrane protein pellet was solubilized in 200 µL 10% SDC in 50 mM HEPES, pH 8.5 by probe-sonication (2x20 sec at 40% amplitude using the Q125 probe-sonicator). After sonication, the sample was diluted to 5% SDC using 50 mM HEPES, pH 8.5 and the protein concentration was measured using a Implen NanoPhotometer N60 (Implen, Germany).

A total of 100 µg of protein from each tissue (six tissues with two replicates from each tissue) and a mixture of proteins from all tissues, was taken out and subjected to reduction using 3mM TCEP and alkylation using 10 mM NEM. After alkylation Endoproteinase Lys-C (0.04AU) was added for 1 hours at 37 °C and subsequently trypsin was added (5%) and incubated over-night at 37 °C. After proteolysis, TMTpro was added to each solution for labeling as following; Kidney (TMT126/127N); Heart (TMT127C/128N); Brain (TMT128C/129N); Liver (TMT129C/130N); BAT(TMT130C/131N); WAT(TMT131C/132N); Mixture (134N). After TMT labeling the samples were mixed and the SDC was precipitated using 2% formic acid and centrifugation at 14000g for 10 min. The SDC pellet was washed and de-palmitoylated peptides were isolated as described above. The purified de-palmitoylated peptides were lyophilized prior to high pH RP separation.

### Analysis of de-palmitoylated peptides from mouse WT and db/db brain using TMTpro quantitation

Proteins from the mouse brains were isolated using 200 µL 5% SDC in 50 mM HEPES, pH 8.5 by probe-sonication (3x20 min at 60% amplitude (Q125 probe-sonicator)). After sonication, the solutions were centrifuged at 14000g for 20 min. The supernatant was transferred to a low binding Eppendorf tube and the protein concentration was measured using a Implen NanoPhotometer N60 (Implen, Germany). A total of 100 µg of protein from four replicates of WT and db/db brain was transferred to new tubes and proteins were reduced, alkylated, and digested with Endoprotease Lys-C and trypsin as described above. The four replicates of WT and db/db brains were labeled with TMTpro (WT (126, 127N, 128N and 129N); db/db (130N, 131N, 132N and 133N)). After labeling the samples were pooled and the de-palmitoylated peptides isolated as described above. The purified de-palmitoylated peptides were lyophilized prior to high pH RP separation.

### High pH Reversed-phase separation

The high pH RP separation was performed essentially as described in[71]. The de-palmitoylated peptides from the 2 replicates of the mouse brain membrane preparation were separated into 12 concatenated fractions directly into a microtiter plate. The de-palmitoylated peptides from the mouse tissue TMT experiment were separated into 20 concatenated fractions and the de-palmitoylated peptides from the db/db mouse brain TMT experiment were separated into 12 concatenated fractions. All the fractions were lyophilized prior to LC-MS/MS.

### Liquid chromatography tandem mass spectrometry (LC-MS/MS)

#### Mouse brain membrane de-palmitoylated peptides

The 12 concatenated fractions from each high pH separation (replicates) were resolubilized in 5µL 0.1% Formic acid and 3 µL was loaded onto a nano-Easy LC (Thermo Fisher Scientific) coupled to an Orbitrap Eclipse mass spectrometer (Thermo Fisher Scientific). Samples were loaded onto a 20 cm 100 ID column containing C18 material (Reprosil 1.9 µm, Dr. Maisch, Ammerbuch-Entringen, Germany). Replicate 1 and 2 was analyzed using a 60-90 min LC gradient. Briefly, the peptides were eluted with a gradient 2-25% buffer B (90% ACN in 0.1% FA) over 50-70 min, 25-40% buffer B over 10-20 min and finally 40-95% buffer B in 1 min. Peptides were selected for 3 second cycle time for MS/MS using higher energy collision dissociation (HCD) with normalized collision energy (NCE) setting as 32-33, resolution of 15-30.000 FWHM, Automatic Gain Control (AGC) target value of 300% and maximum injection time of 100 ms.

#### Mouse tissue TMTpro experiment

The 20 concatenated fractions from each high pH separation were resolubilized in 3.5 µL 0.1% Formic acid and 3 µL was loaded onto a 20 cm 100 ID column containing C18 material (Reprosil 1.9 µm, Dr. Maisch, Ammerbuch-Entringen, Germany) on a nano-Easy LC (Thermo Fisher Scientific) coupled to an Orbitrap Exploris 480 mass spectrometer (Thermo Fisher Scientific). The peptides were eluted using a 90 min gradient as described above. Peptides were selected for 3 second cycle time for MS/MS using isolation window of 0.7 Da, HCD with NCE 33, a resolution of 45.000 FWHM, AGC target value of 300% and maximum injection time of 100 ms.

#### Mouse WT and db/db TMTpro experiment

The 12 concatenated fractions from each high pH separation were resolubilized in 3.5 µL 0.1% Formic acid and 3 µL was loaded onto a 20 cm 100 ID column containing C18 material (Reprosil 1.9 µm, Dr. Maisch, Ammerbuch-Entringen, Germany) on a nano-Easy LC (Thermo Fisher Scientific) coupled to an Orbitrap Exploris 480 mass spectrometer (Thermo Fisher Scientific). The peptides were eluted using a 120 min gradient (2-30% buffer B (90% ACN in 0.1% FA) over 100 min, 30-50% buffer B over 20 min and finally 50-95% buffer B in 1 min). Peptides were selected for 3 second cycle time for MS/MS using isolation window of 0.7 Da, HCD with NCE 33, a resolution of 45.000 FWHM, AGC target value of 300% and maximum injection time of 100 ms.

### Peptide identification and quantitation

All raw files for each replicate for the mouse brain palmitoylome mapping were searched in Proteome Discoverer (PD) 2.5.0.400 (Thermo Fisher Scientific, USA) using both the SEQUEST HT (EMBL_mouse.fasta database (21.816 sequences), and Mascot (Uniprot database of *Mus Musculus,* 25.775 sequences) search algorithms . Search parameters were as following: cleavage specificity trypsin/P; precursor mass tolerance of 10 ppm, and fragment mass tolerance of 0.05 Da. Variable modifications: Methionine oxidation, N-terminal Acetylation, N-terminal Met-loss, N-terminal Met loss + Acetyl, NEM(C) and carbamidomethyl (C). The percolator software in PD was used for filtering for false discovery rate (FDR) of <1% for peptides. All data sets from PD were exported to Excel (Microsoft) for further processing.

For the TMTpro experiments, the raw files were searched in PD 2.5.0.400 using both the SEQUEST HT and Mascot search algorithms as above. Search parameters were as following: cleavage specificity trypsin/P; precursor mass tolerance of 10 ppm, and fragment mass tolerance of 0.05 Da. Variable modifications: NEM(C) and carbamidomethyl (C). Fixed modifications: TMTpro (K and N-term). The percolator software in PD was used for filtering for false discovery rate (FDR) of <1% for peptides. All data sets from PD were exported to Excel (Microsoft) for further processing.

The mass spectrometry proteomics data have been deposited to the ProteomeXchange Consortium via the PRIDE partner repository.

### Bioinformatics

Venn diagrams were created using the Venn.diagram package in R. All datasets containing unique sites from the SwissPalm database were downloaded 31-05-2023 (Release 4, 2022-09-03).

Functional analysis of identified de-palmitoylation sites in the mouse brain samples were performed using the Cytoscape Stringapp (ver. 2.0.0) in Cytoscape (Ver.3.10.1) [72, 73]. Prior to functional analysis, data were clustered using Cytoscape ClusterMaker2 (ver. 2.3.2) [74]. Functional analysis was performed separately on each cluster against the entire dataset as background. Redundant terms were filtered out and Only Gene Ontology (GO) terms (cellular compartment, molecular function and biological processes) and Kyoto Encyclopedia of Genes and Genomes (KEGG) pathways were considered. To space nodes in the string map for better visualization, we used the yfiles Layout Algorithm (ver. 1.1.2) organic layout function. For comparing and removing disulfide bonds, we used Uniprot disulfide annotation data (downloaded 12-02-2024) and filtered our data using a homemade R script. Analysis of mouse organ tissues were performed in R with kmeans clustering and using R and Stringapp for GO/KEGG analysis and visualization. Statistical test of db/db mouse brain palmitoylome were performed using the PolySTest [75]. S-palmitoylated peptides with a p-value < 0.05 were considered statistically significant. Volcano plot was generated using R. For string analysis of db/db mouse brain samples, we used the OmicsVisualizer app for plotting quantitative values (log_2_ fold changes) [76]. Graphical structure of WDFY1 were created from icn3D accessed at National Center for Biotechnology Information [77].

## Supplementary figure legends

**Figure S1:**
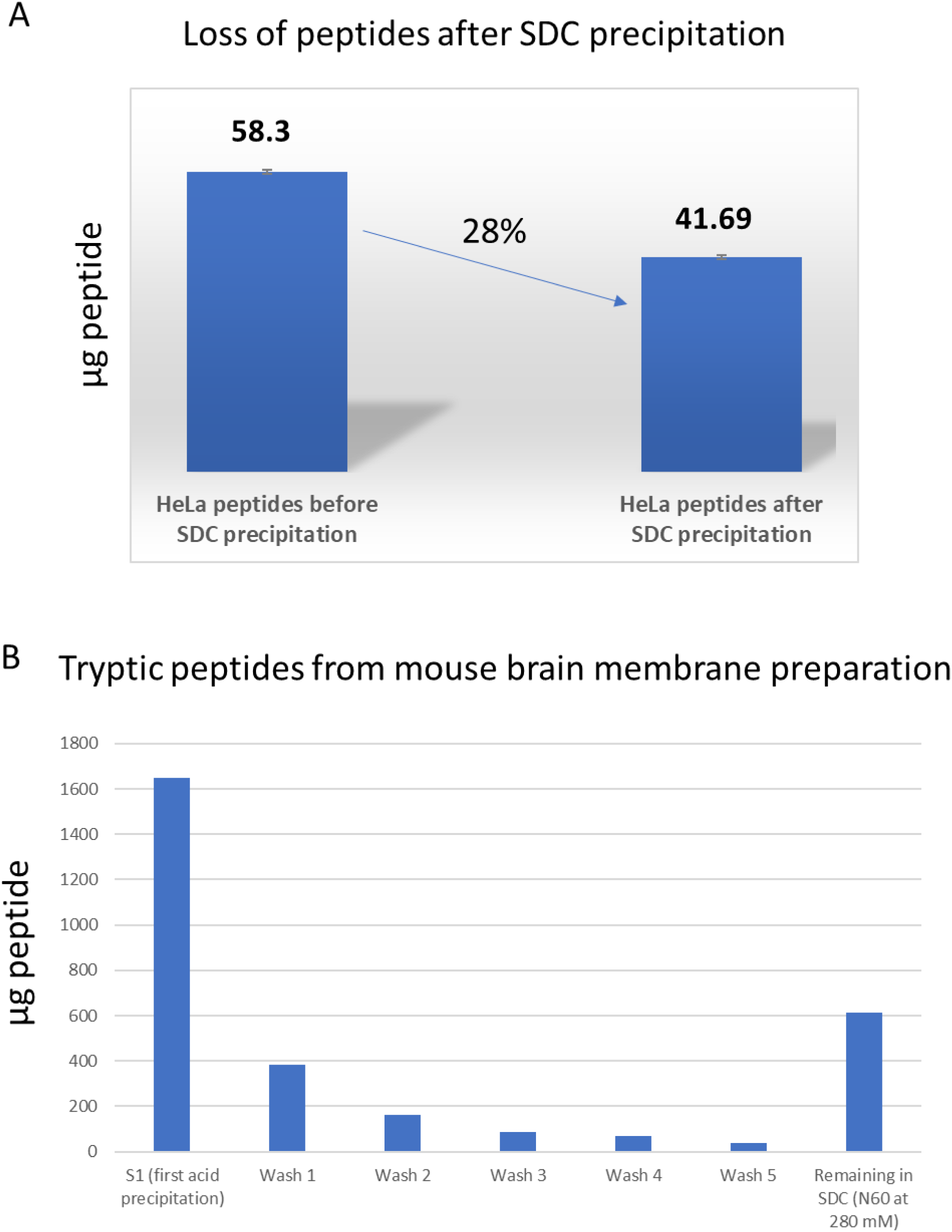
SDC acid precipitation of HeLa cells and mouse brain membrane preparation. A) An aliquot corresponding to 58.3 µg of peptides originating from tryptic digestion of a HeLa cell lysate was taken out and added SDC to a total of 1% SDC. After SDC acid precipitation a loss of 28% of the peptides was lost in the SDC pellet. Amounts of peptides was determined by amino acid composition analysis (AAA). B) Proteins from a mouse brain membrane preparation was digested with trypsin in 5% SDC. The figure shows the amounts of peptides present in the first acid precipitation (S1), the 5 subsequent washes (see methods), and the final solubilized SDC pellet. The peptide/protein was measured by nanodrop at 280 nm.

**Figure S2:**
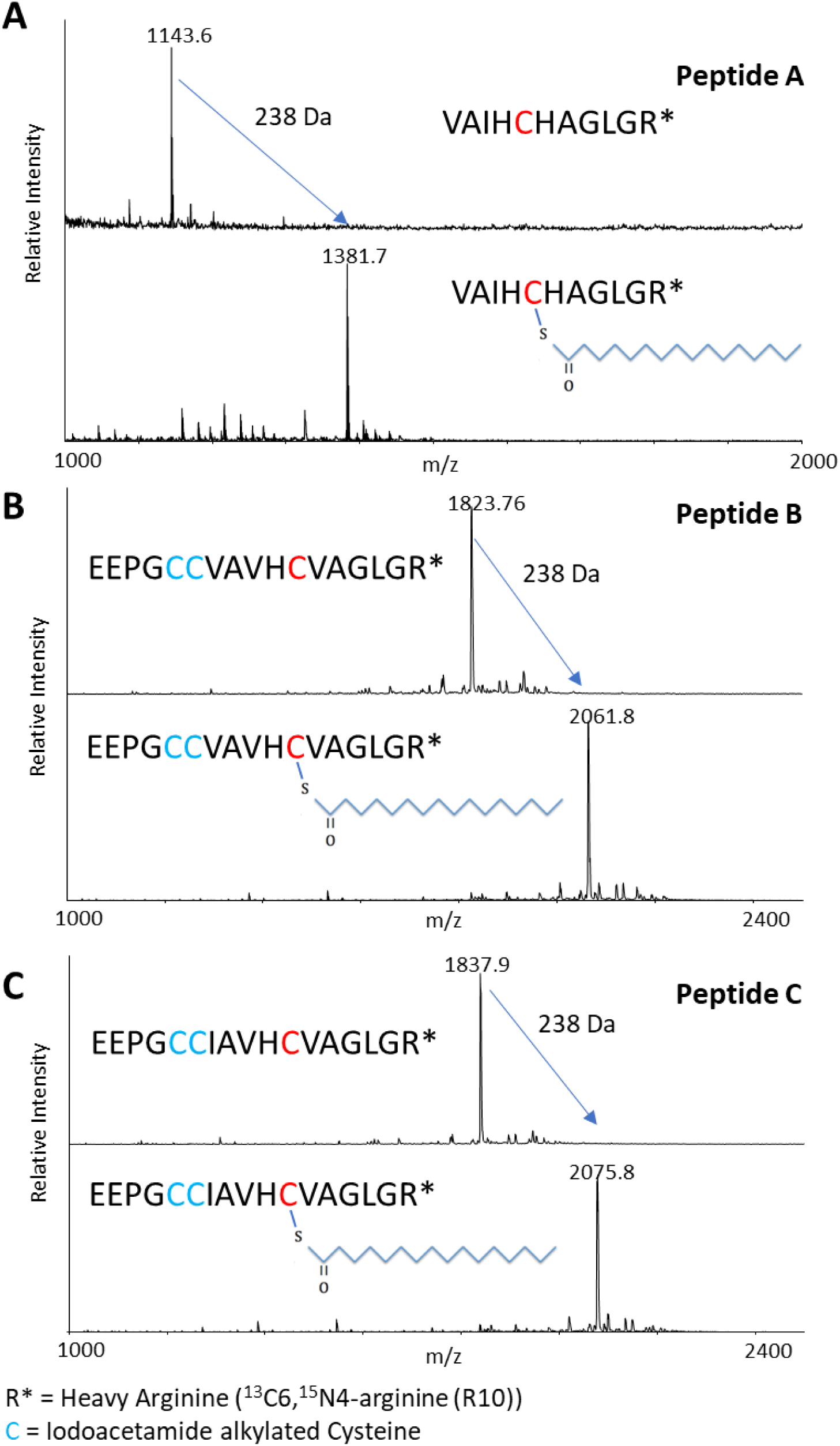
MALDI-MS analysis of S-palmitoylated synthetic peptides. MALDI-MS spectra showing three synthetic peptides (A-C) before and after the S-palmitoylation reaction, resulting in an increase in 238 Da for each cysteine-containing peptide.

**Figure S3:**
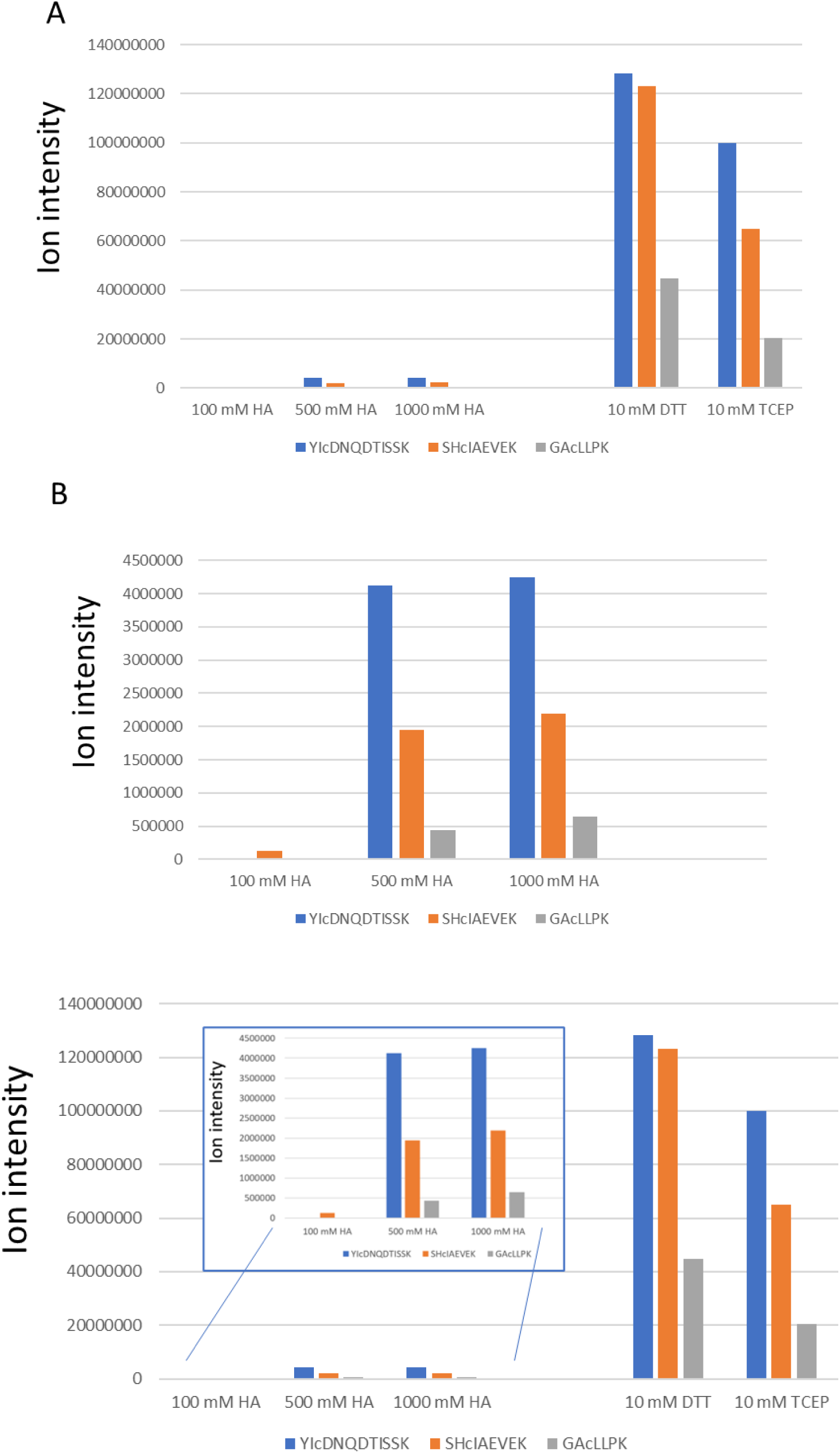
Assessment of disulfide reduction by HA. To assess the specificity of HA, we applied bovine serum albumin proteins, alkylating free cysteines with NEM followed by tryptic digestion and reduction using HA, DTT or TCEP.

**Figure S4:**
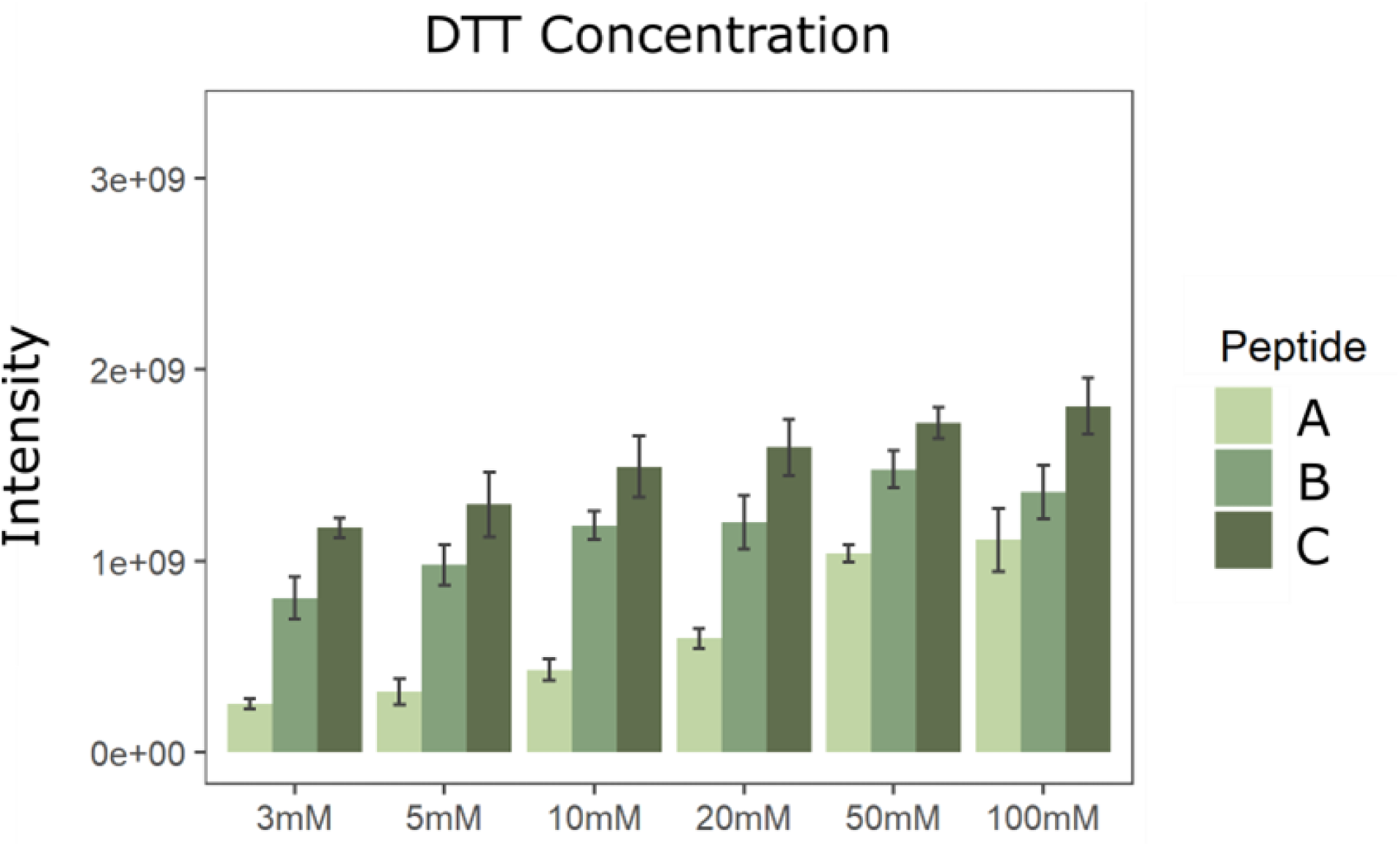
Optimizing the DTT concentration for de-palmitoylation. For testing the optimal concentration of DTT for de-palmitoylation, the three S-palmitoylated synthetic peptides were mixed and divided into equal volumes. Each sample received one of the following concentrations of DTT; 3mM, 5mM, 10mM, 20mM, 50mM or 100mM. Peptide mix were incubated for 1 hour at room temperature with DTT. The results are average of triplicate experiment.

**Figure S5:**
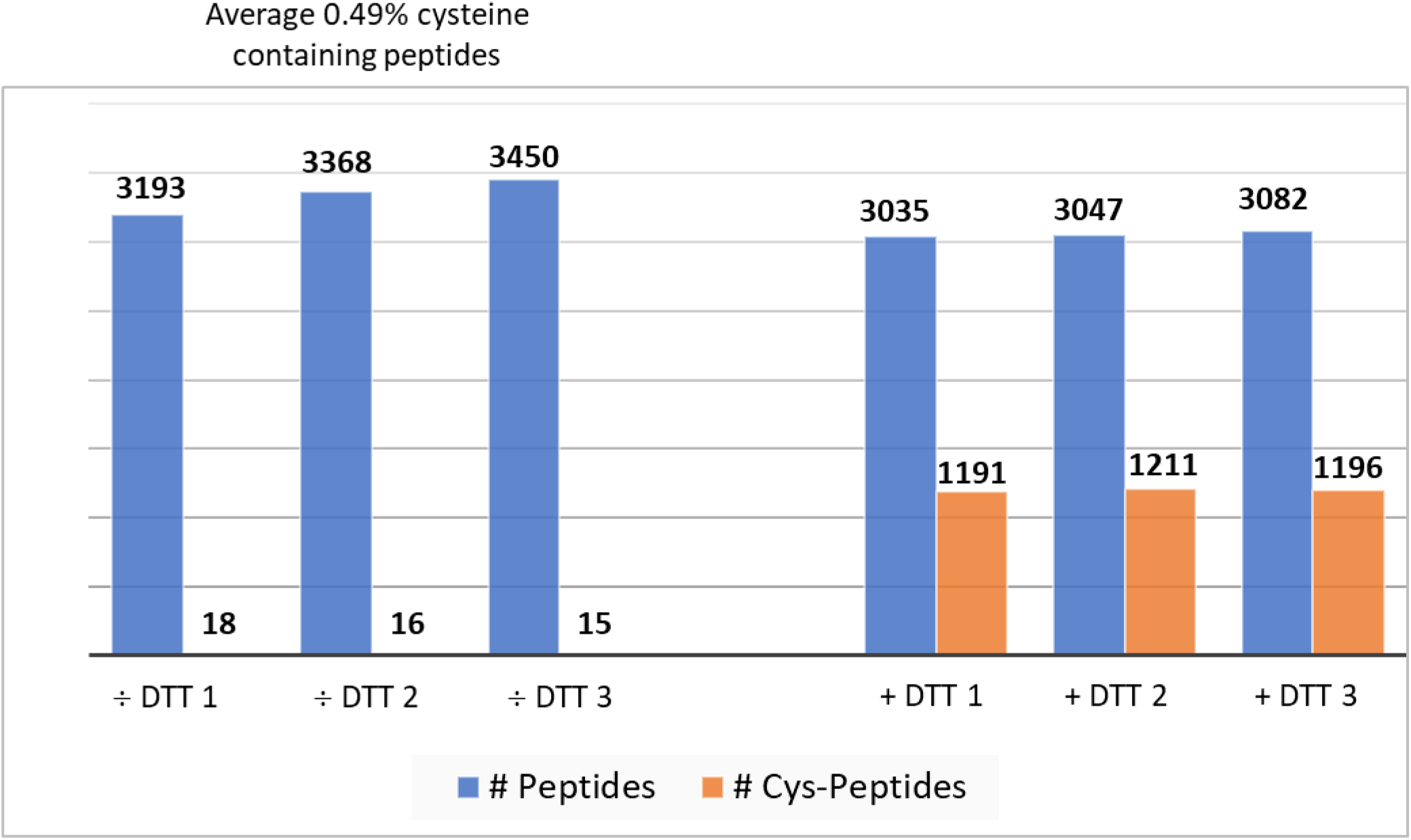
Mouse brain control test with or without DTT. Aliquots of the mouse brain membrane preparation digested with trypsin in 5% SDC were aliquoted and treated with or without 20mM DTT in triplicates. From the triplicates without DTT (÷ DTT 1-3) only an average of ≈ 0.49 % cysteine containing peptides were identified in the eluate compared to ≈ 39% cysteine containing peptides from the samples with DTT (+ DTT 1-3).

**Figure S6:**
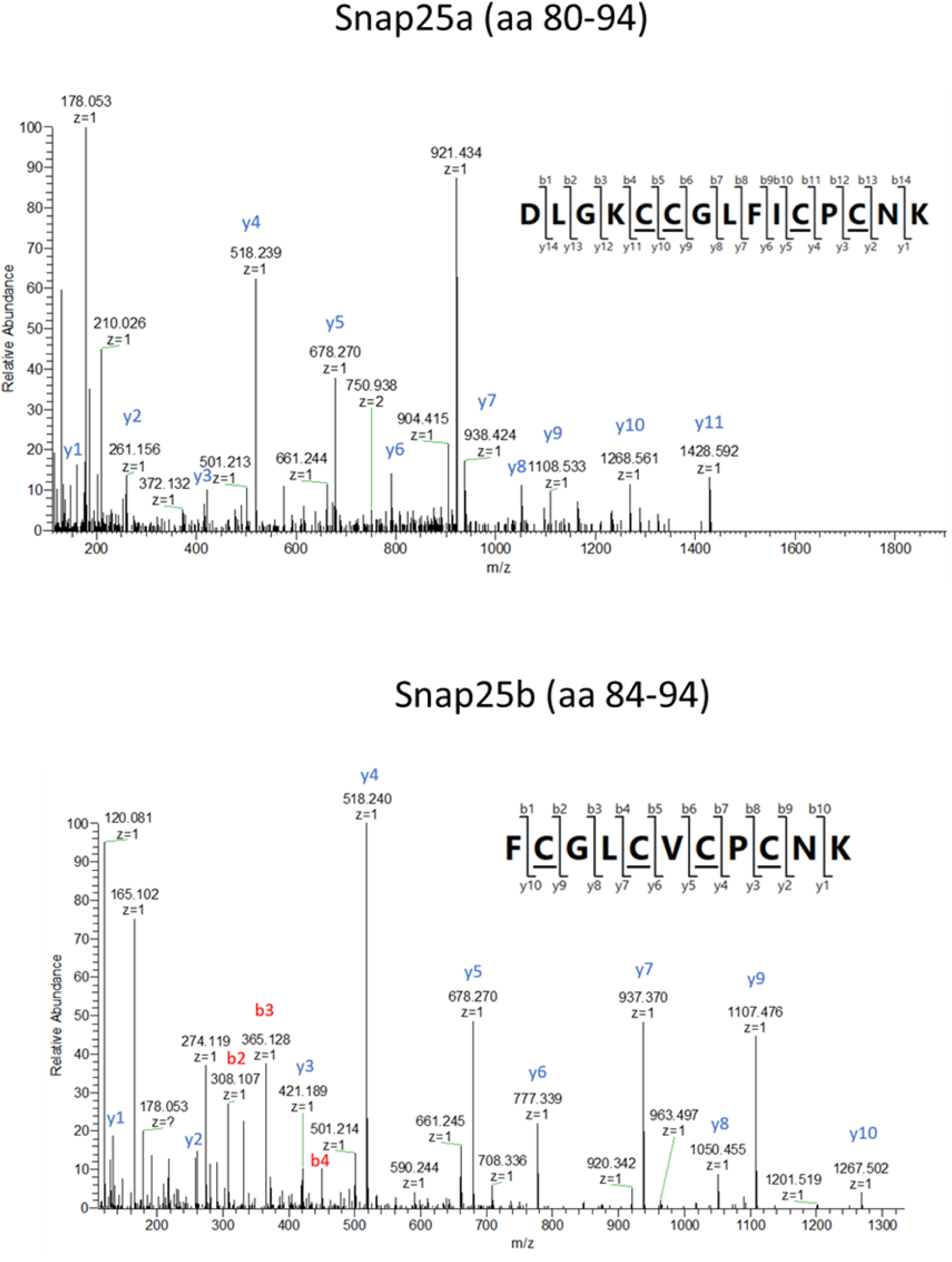
Annotated tandem mass spectra for the peptides containing the S-palmitoylated sites in Snap25 in mouse brains. A well-known S-palmitoylated protein, Snap25, were identified as S-palmitoylated in both splicing variants (Snap25a and Snap25b). S-palmitoylated sites identified are underscored in the peptide sequence.

**Figure S7:**
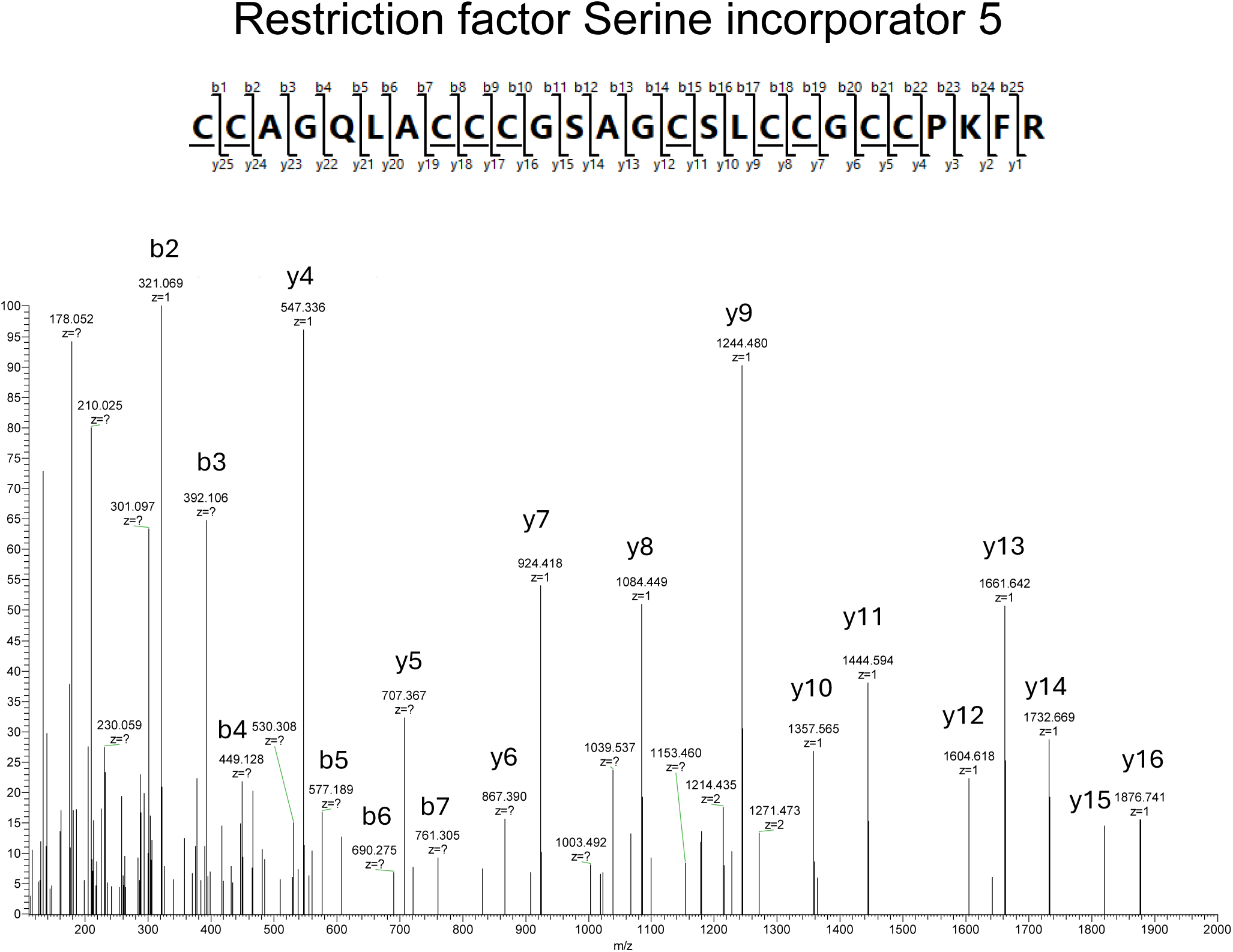
S-palmitoylation of Serine Incorporator 5. Annotated tandem mass spectrum of a peptide from the Serine Incorporator 5 restriction factor containing 10 S-palmitoylation sites on a single peptide sequence. S-palmitoylated cysteines are underscored in the peptide sequence.

**Figure S8:**
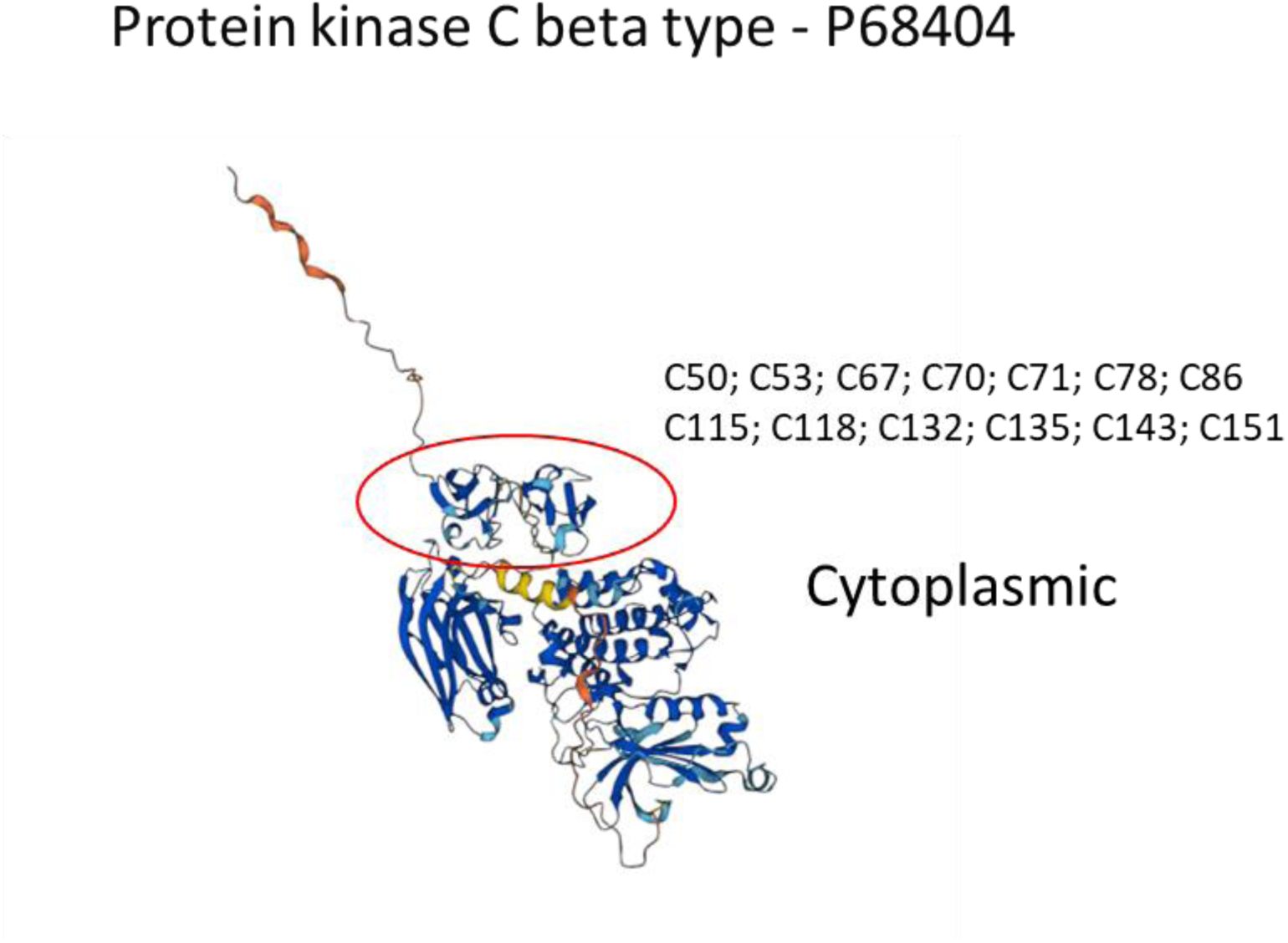
3D structure of PKC beta type. 3D structure of the conventional PKC beta type, red circle illustrating N-terminal domain which contains several loops and disordered regions with S-palmitoylation identified in our study.

**Figure S9:**
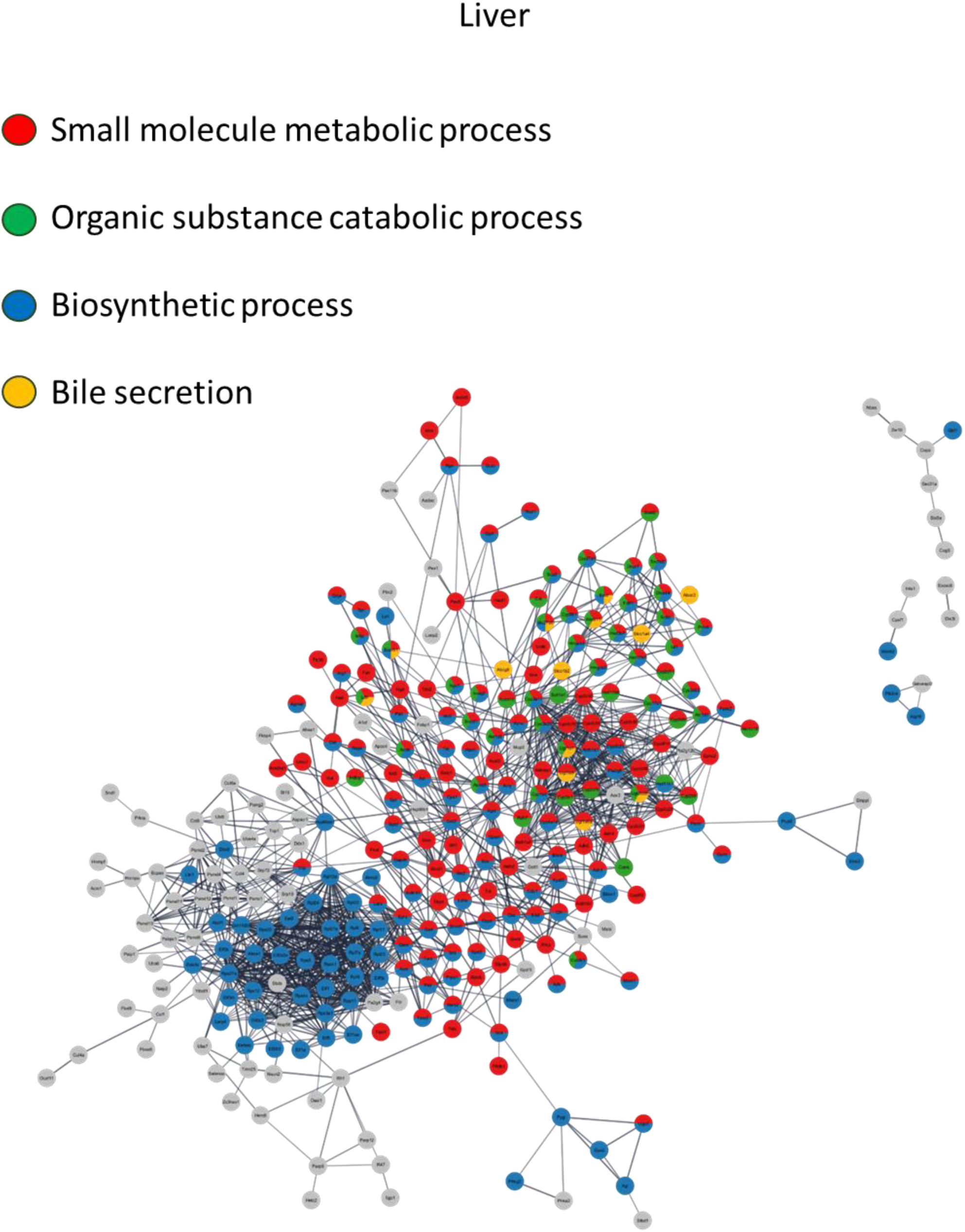
Palmitoylome of mouse liver tissue. GO/KEGG enrichment of cluster 3 following protein filtering for upregulated features with a log_2_ value above 2 in at least one replicate across all conditions. Analysis was performed with Cytoscape Stringapp, string confidence: 0.7.

**Figure S10:**
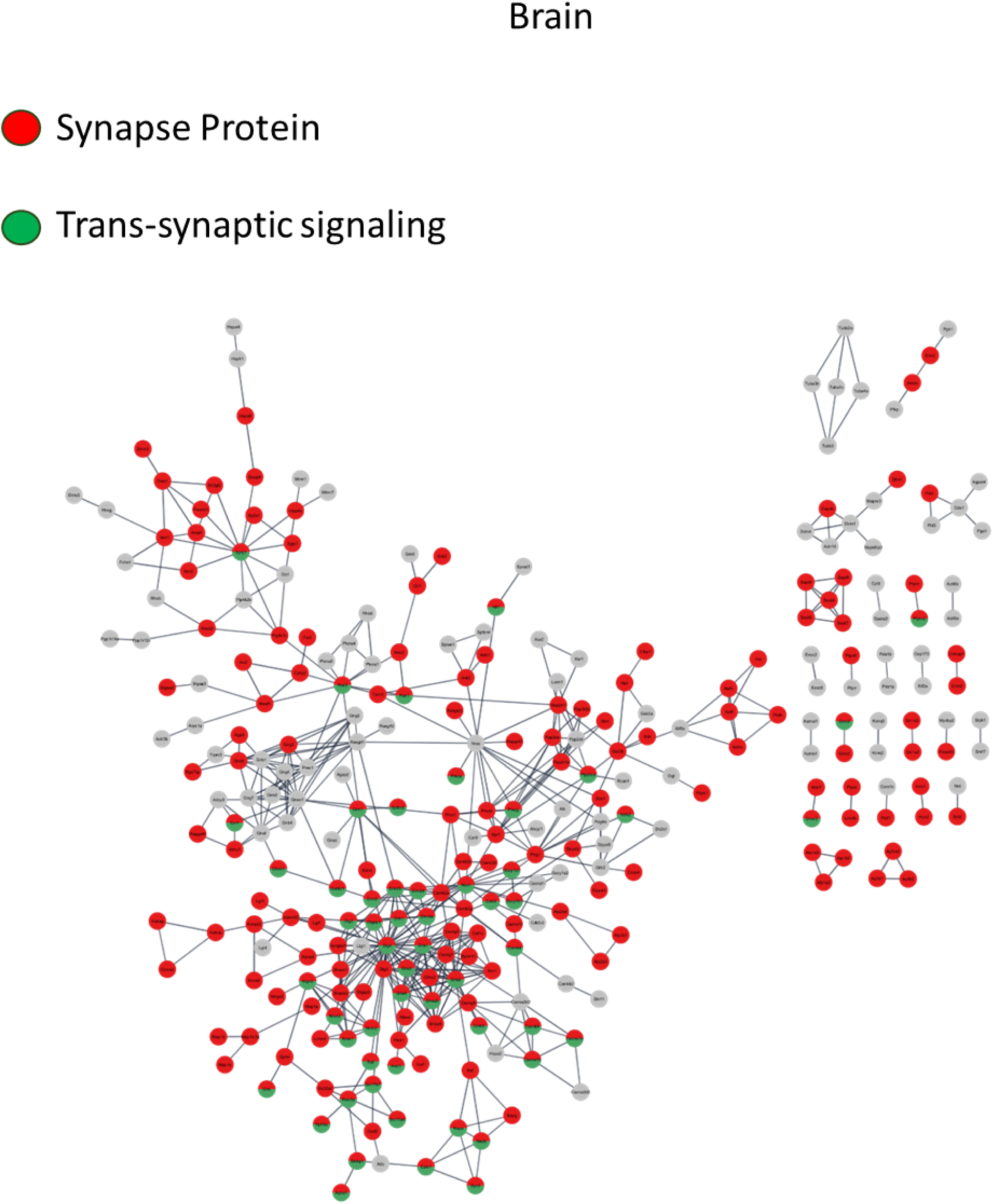
Palmitoylome of mouse brain tissue. GO/KEGG enrichment of cluster 4 protein filtering for upregulated features with a log_2_ value above 2 in at least one replicate across all conditions. Analysis was performed with Cytoscape Stringapp, string confidence: 0.9.

**Figure S11:**
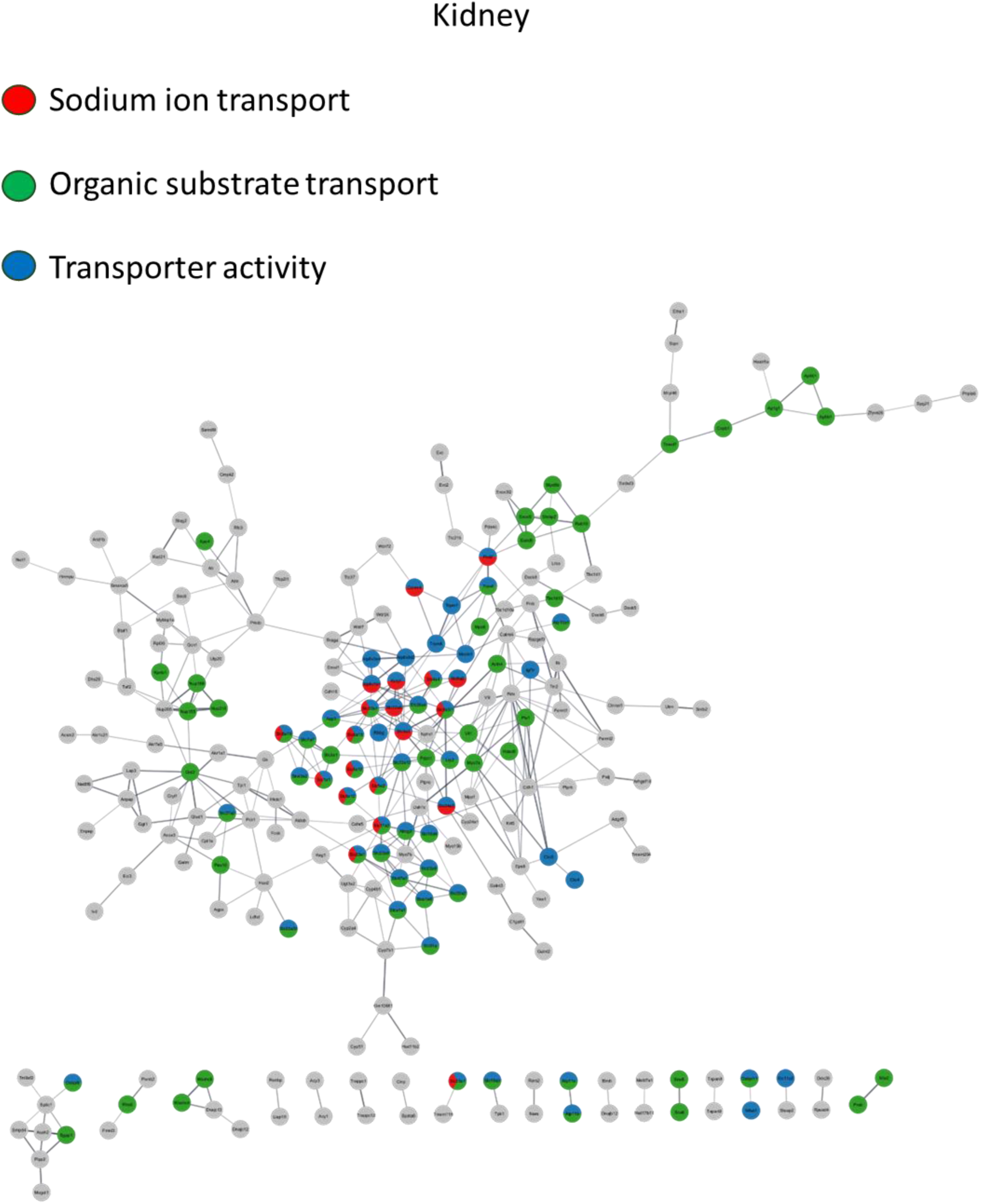
Palmitoylome of mouse kidney tissue. GO/KEGG enrichment of cluster 1 following protein filtering for upregulated features with a log_2_ value above 2 in at least one replicate across all conditions. Analysis was performed with Cytoscape Stringapp, string confidence: 0.5.

## Notes

### Competing Interest Statement

The authors have declared no competing interest.

### Summary of Updates

Manuscript is updated and revised and new data included

