## Supplementary zDHHC 3D structures for "Enrichment of Cysteine S-palmitoylated peptides using Sodium Deoxycholate Acid Precipitation - SDC-ACE"

### Slide 1
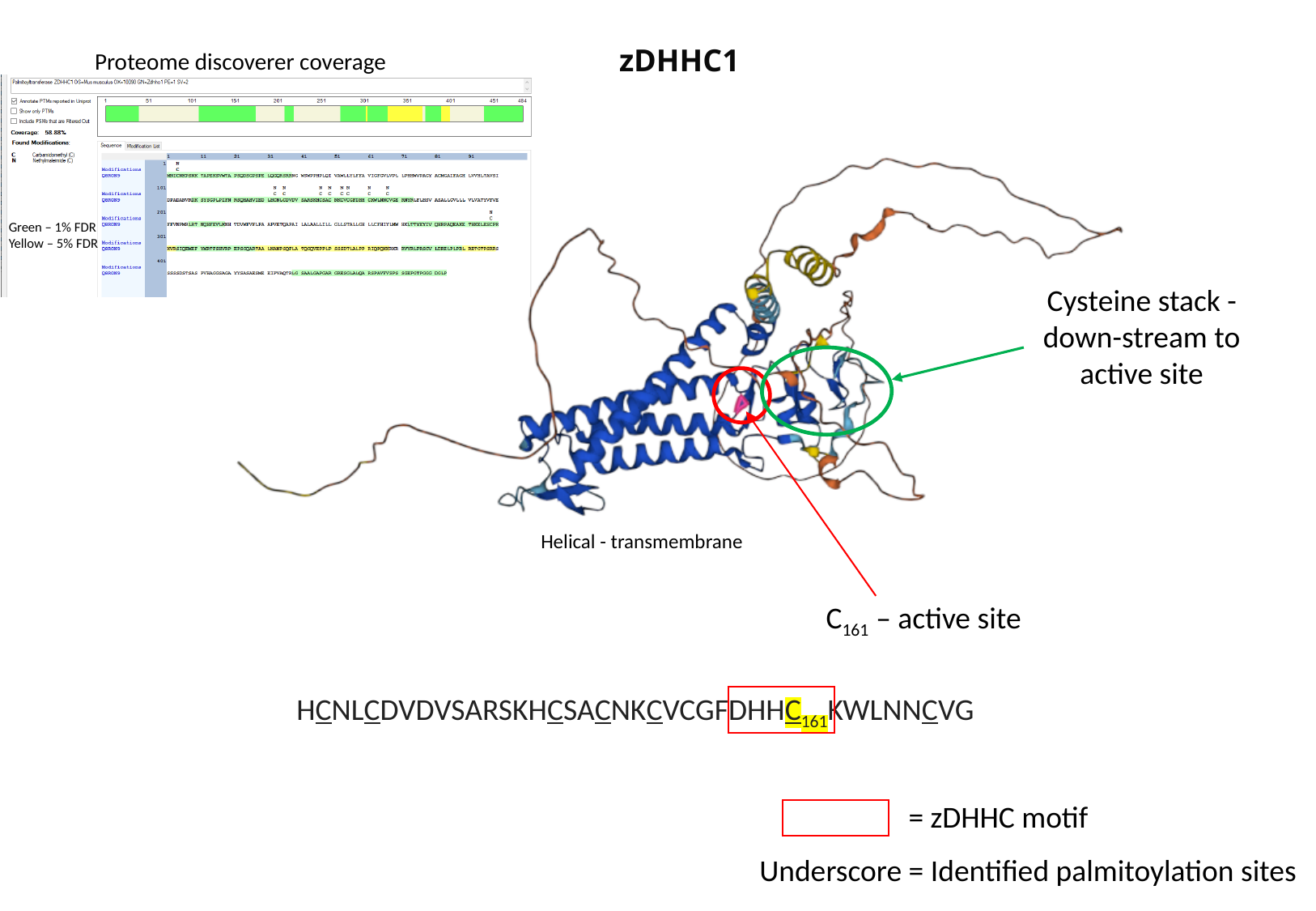

zDHHC1
Proteome discoverer coverage
Green – 1% FDR
Yellow – 5% FDR
Cysteine stack - down-stream to active site
Helical - transmembrane
C_161_ – active site
HCNLCDVDVSARSKHCSACNKCVCGFDHHC_161_KWLNNCVG
= zDHHC motif
Underscore = Identified palmitoylation sites

### Slide 2
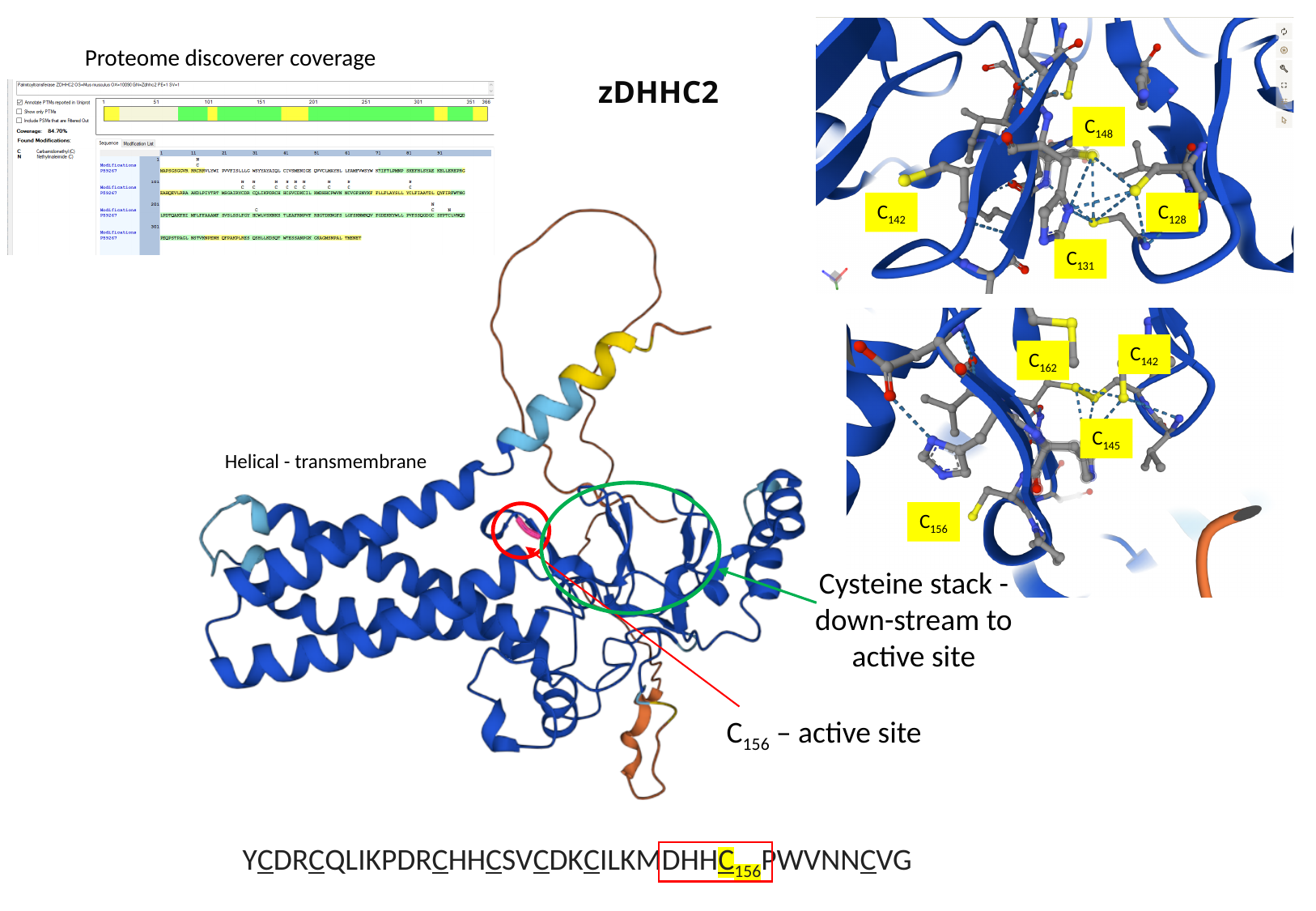

Proteome discoverer coverage
zDHHC2
C_148_
C_142_
C_128_
C_131_
C_142_
C_162_
C_145_
Helical - transmembrane
C_156_
Cysteine stack - down-stream to active site
C_156_ – active site
YCDRCQLIKPDRCHHCSVCDKCILKMDHHC_156_PWVNNCVG

### Slide 3
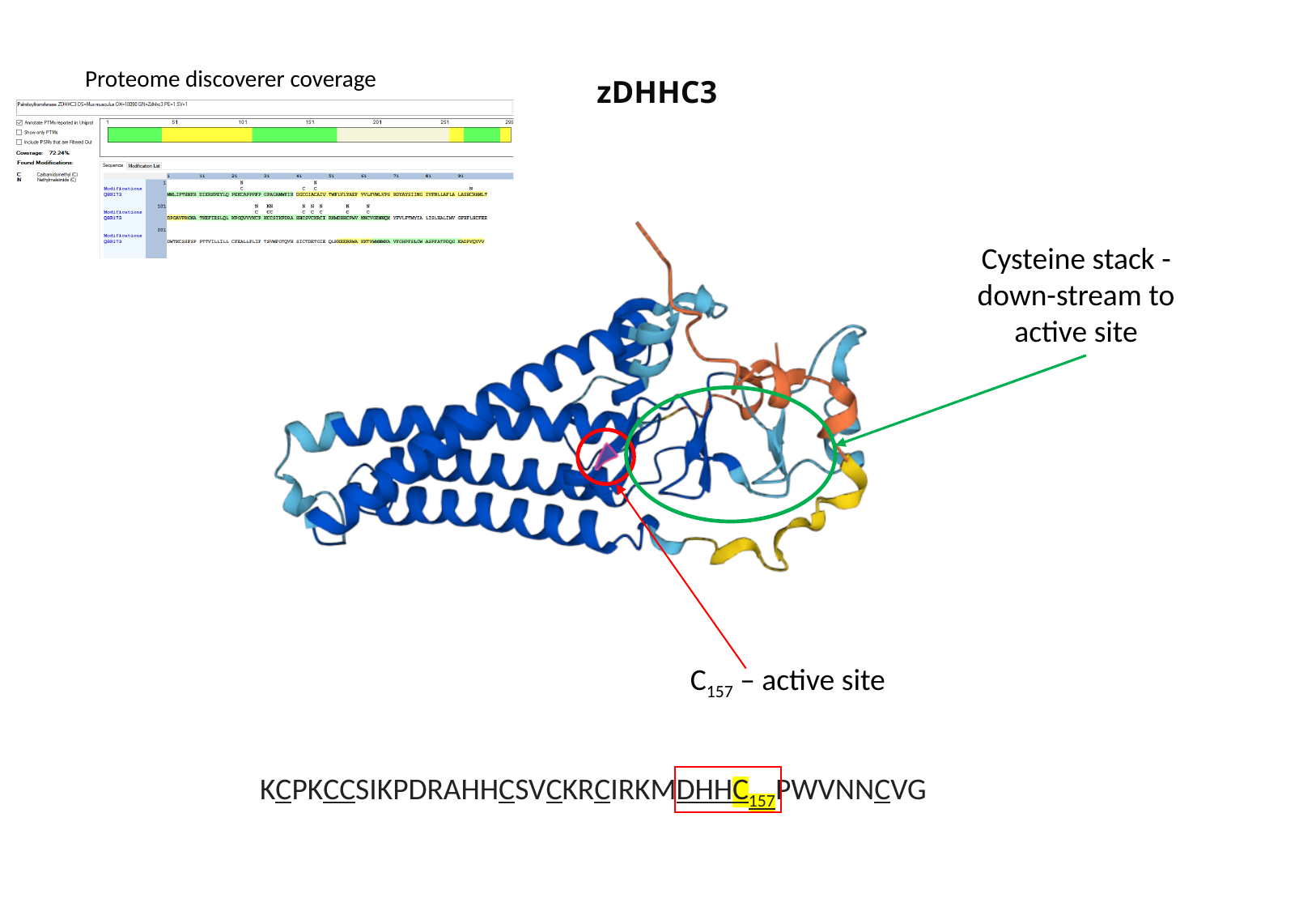

Proteome discoverer coverage
zDHHC3
Cysteine stack - down-stream to active site
C_157_ – active site
KCPKCCSIKPDRAHHCSVCKRCIRKMDHHC_157_PWVNNCVG

### Slide 4
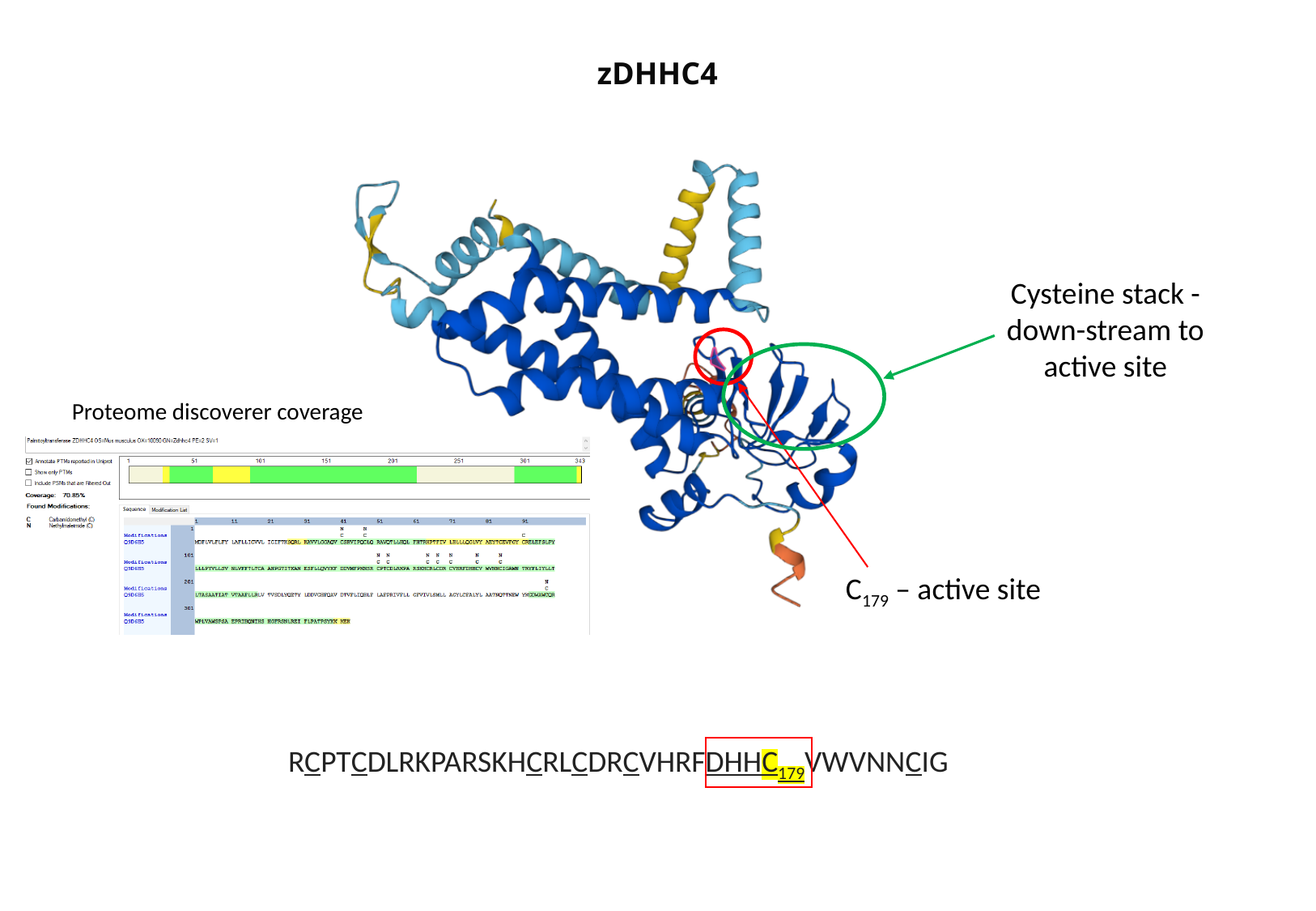

zDHHC4
Cysteine stack - down-stream to active site
Proteome discoverer coverage
C_179_ – active site
RCPTCDLRKPARSKHCRLCDRCVHRFDHHC_179_VWVNNCIG

### Slide 5
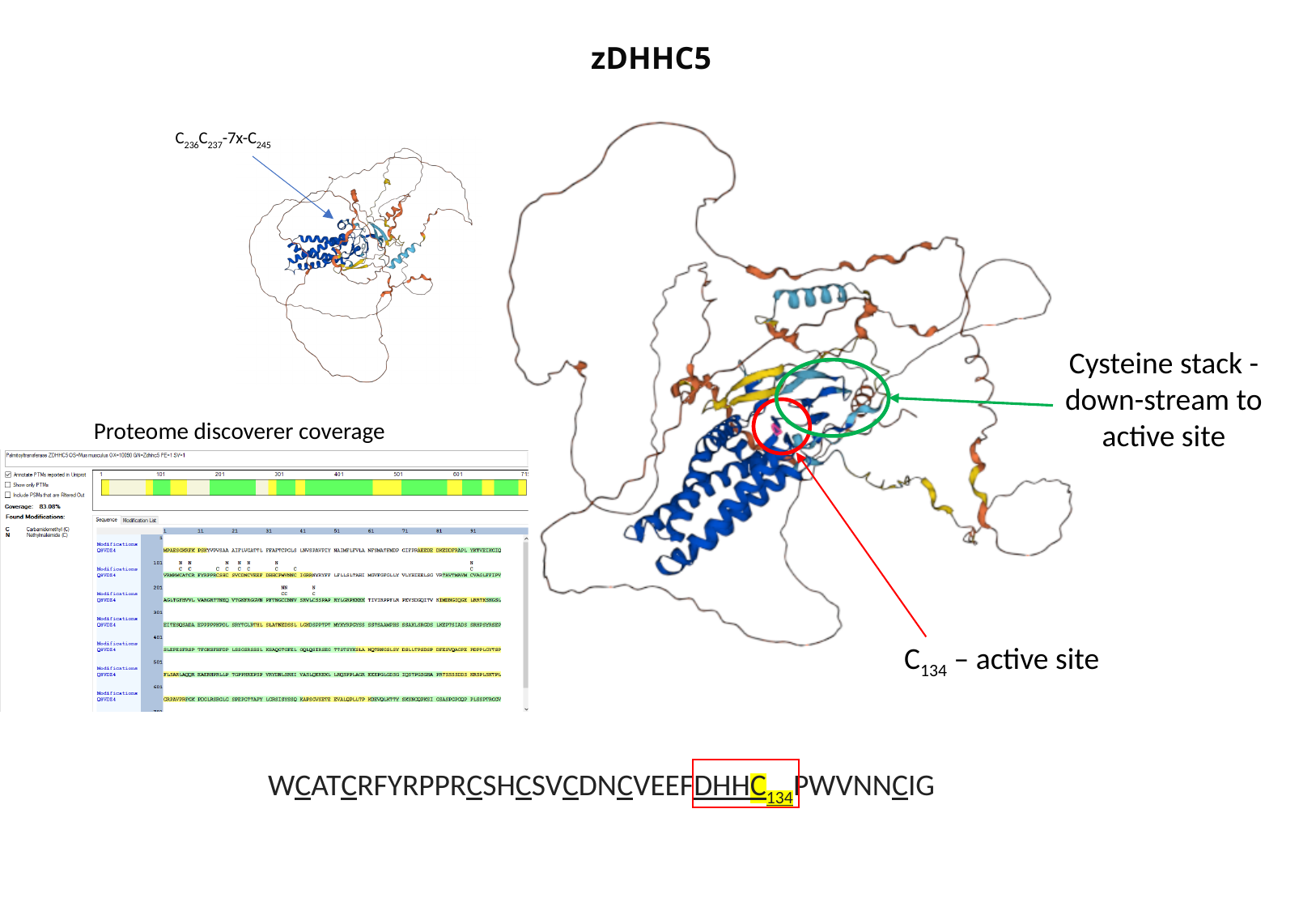

zDHHC5
C_236_C_237_-7x-C_245_
Cysteine stack - down-stream to active site
Proteome discoverer coverage
C_134_ – active site
WCATCRFYRPPRCSHCSVCDNCVEEFDHHC_134_PWVNNCIG

### Slide 6
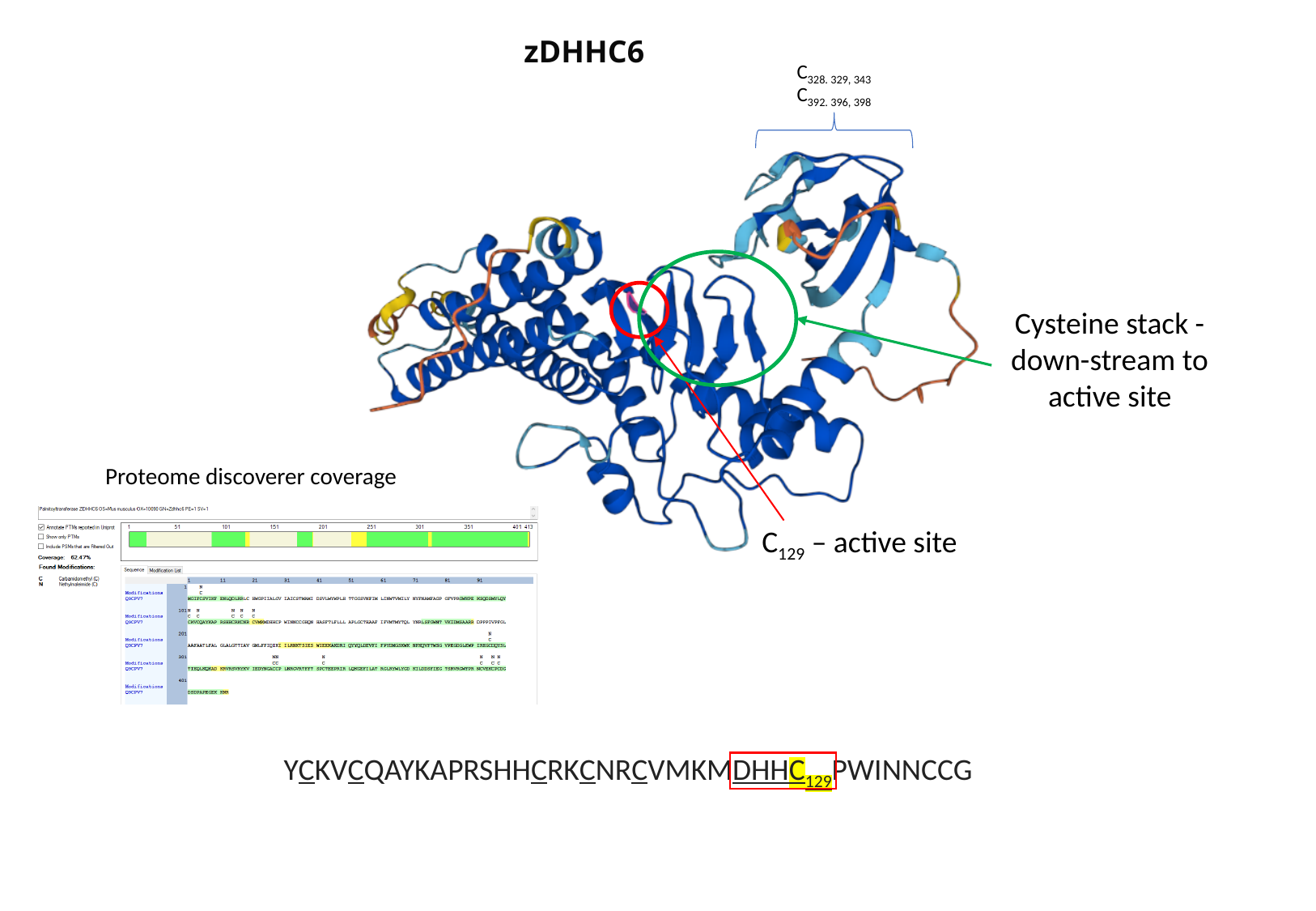

zDHHC6
C_328. 329, 343_
C_392. 396, 398_
Cysteine stack - down-stream to active site
Proteome discoverer coverage
C_129_ – active site
YCKVCQAYKAPRSHHCRKCNRCVMKMDHHC_129_PWINNCCG

### Slide 7
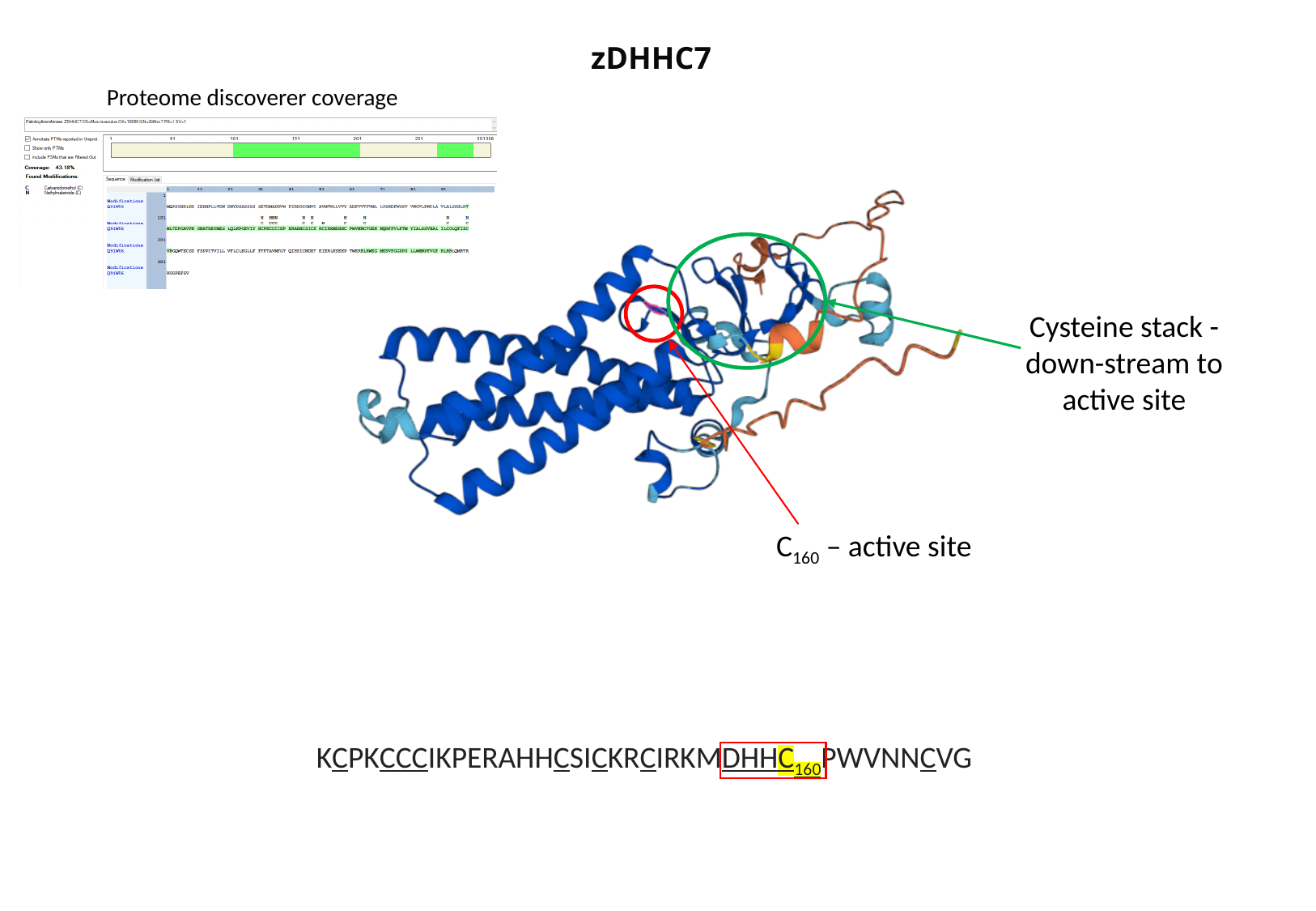

zDHHC7
Proteome discoverer coverage
Cysteine stack - down-stream to active site
C_160_ – active site
KCPKCCCIKPERAHHCSICKRCIRKMDHHC_160_PWVNNCVG

### Slide 8
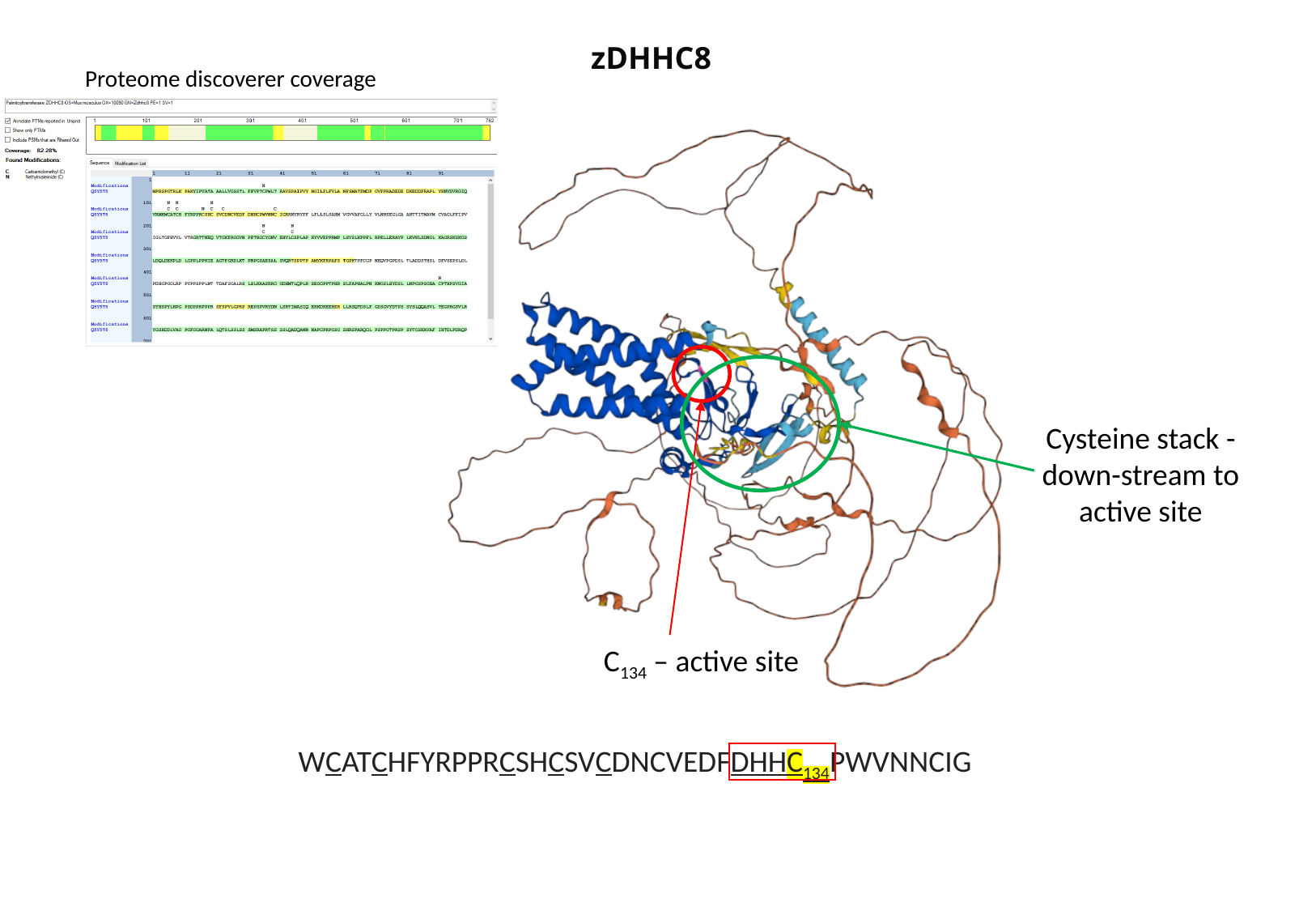

zDHHC8
Proteome discoverer coverage
Cysteine stack - down-stream to active site
C_134_ – active site
WCATCHFYRPPRCSHCSVCDNCVEDFDHHC_134_PWVNNCIG

### Slide 9
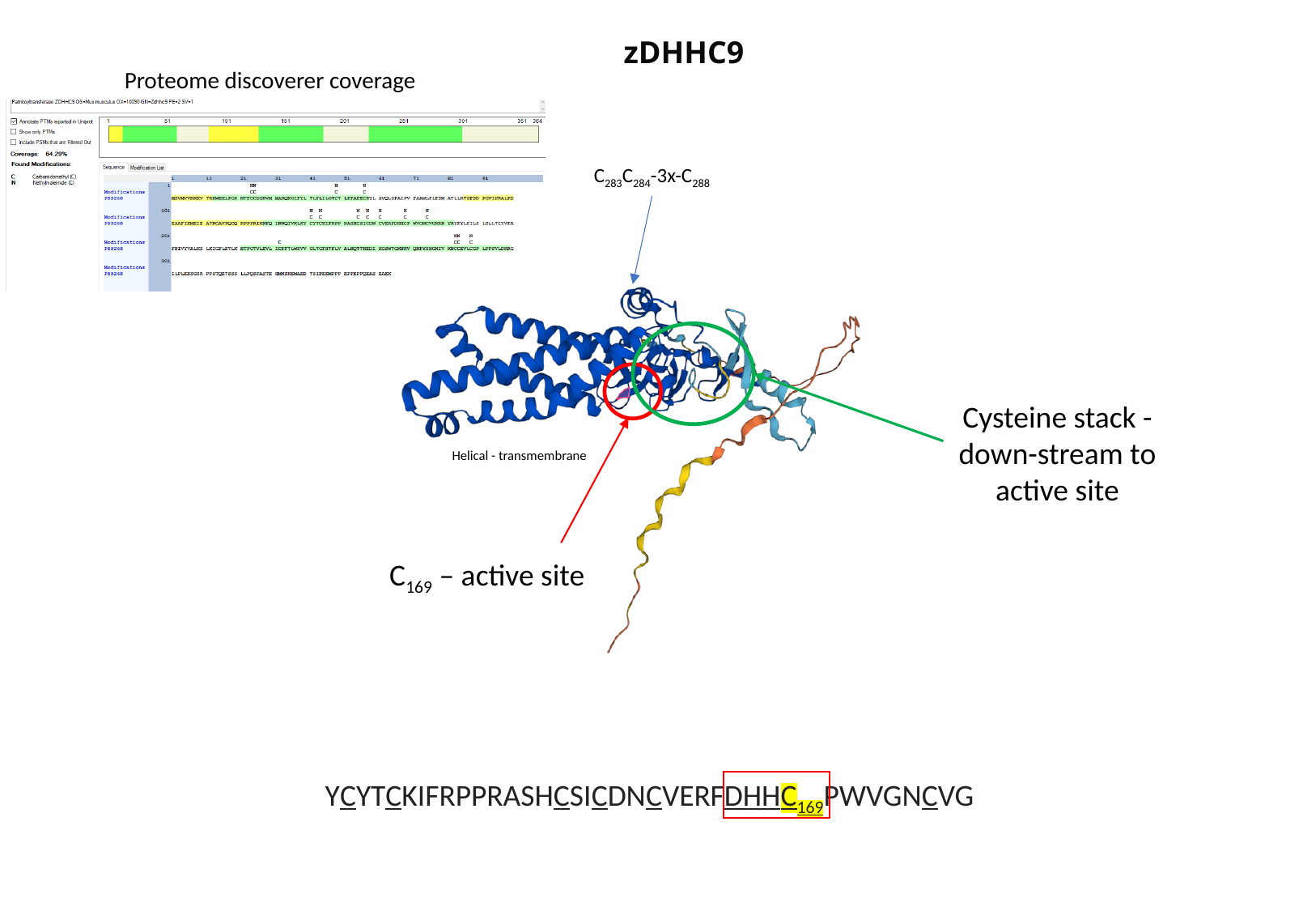

zDHHC9
Proteome discoverer coverage
C_283_C_284_-3x-C_288_
Cysteine stack - down-stream to active site
Helical - transmembrane
C_169_ – active site
YCYTCKIFRPPRASHCSICDNCVERFDHHC_169_PWVGNCVG

### Slide 10
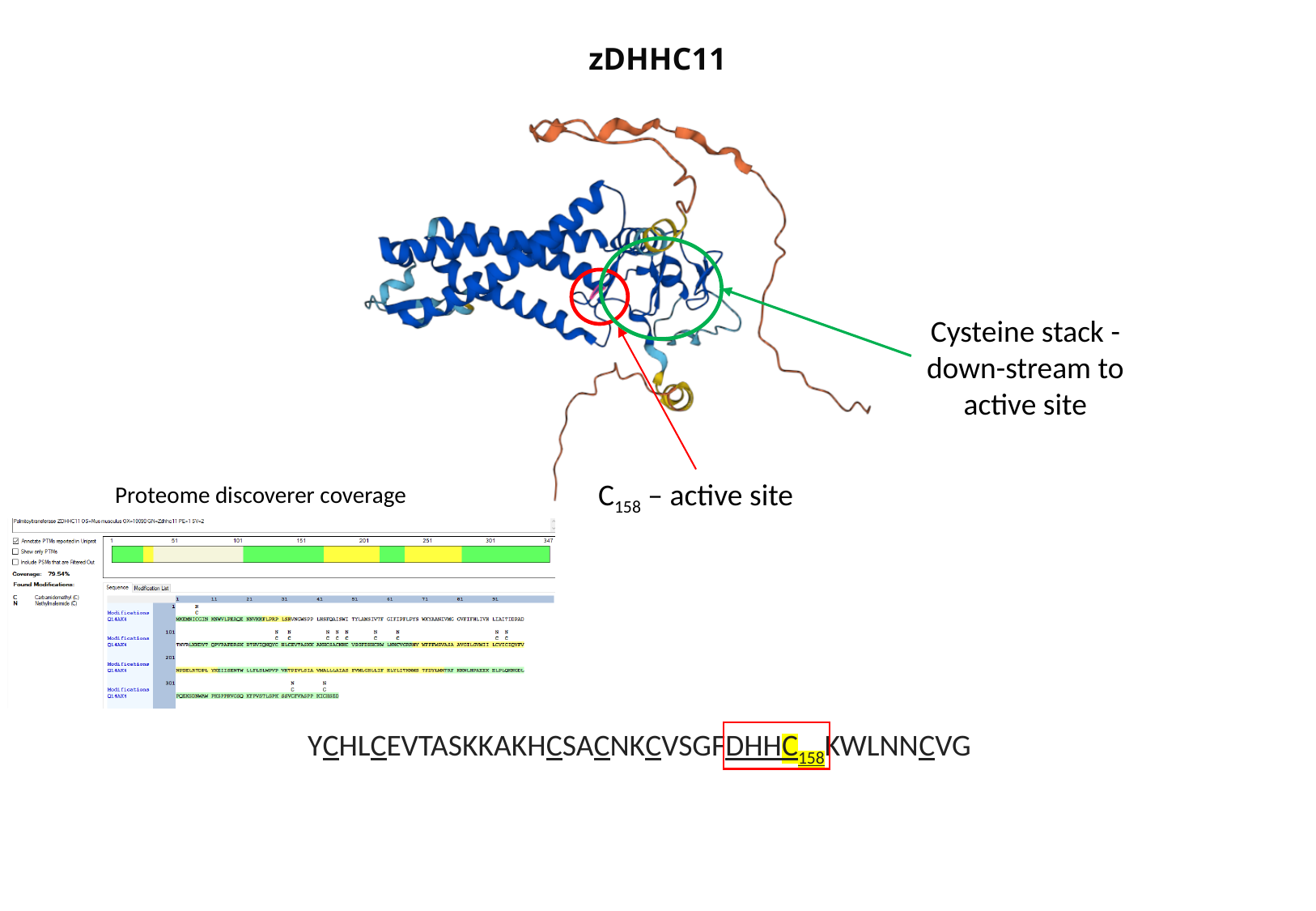

zDHHC11
Cysteine stack - down-stream to active site
C_158_ – active site
Proteome discoverer coverage
YCHLCEVTASKKAKHCSACNKCVSGFDHHC_158_KWLNNCVG

### Slide 11
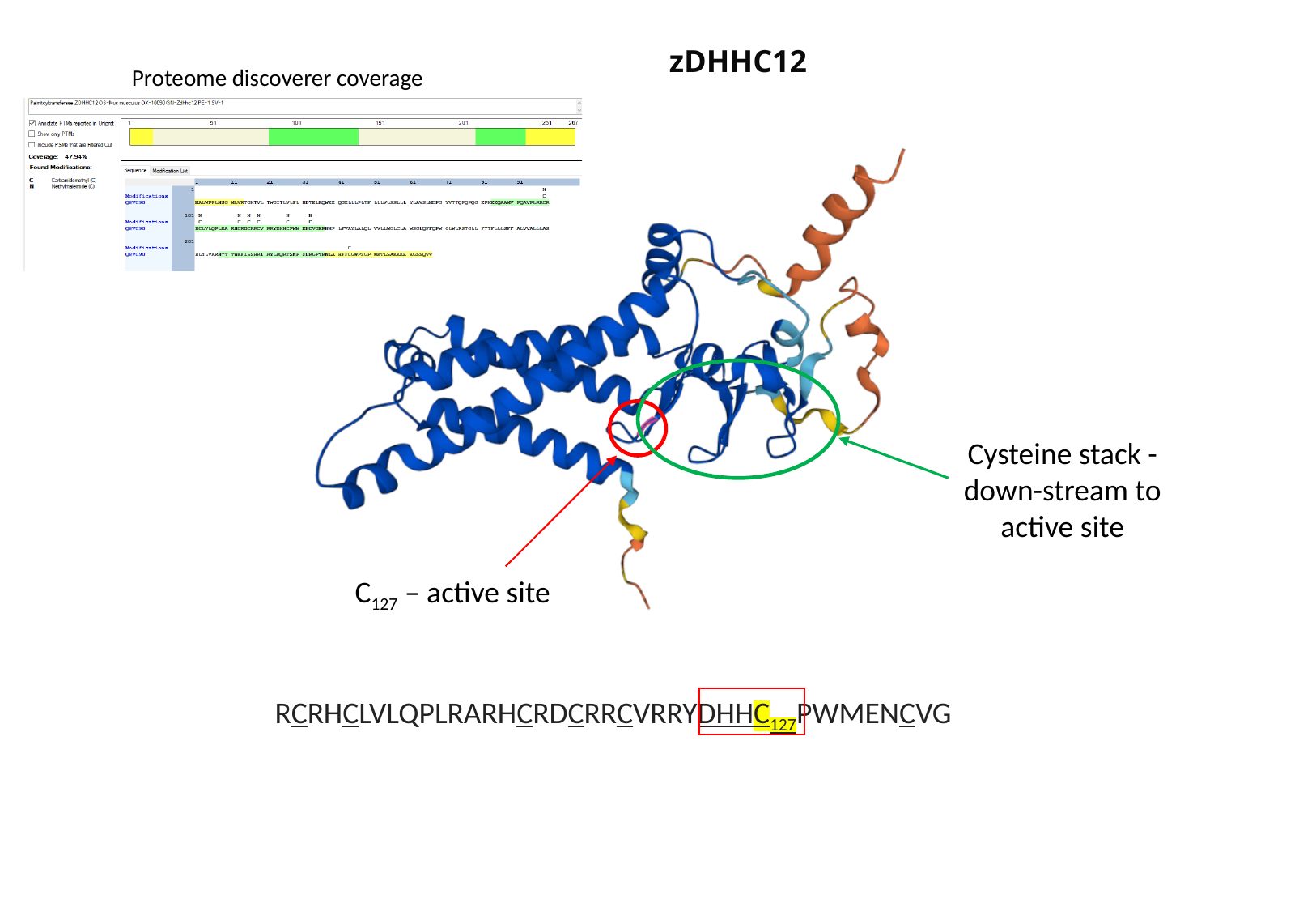

zDHHC12
Proteome discoverer coverage
Cysteine stack - down-stream to active site
C_127_ – active site
RCRHCLVLQPLRARHCRDCRRCVRRYDHHC_127_PWMENCVG

### Slide 12
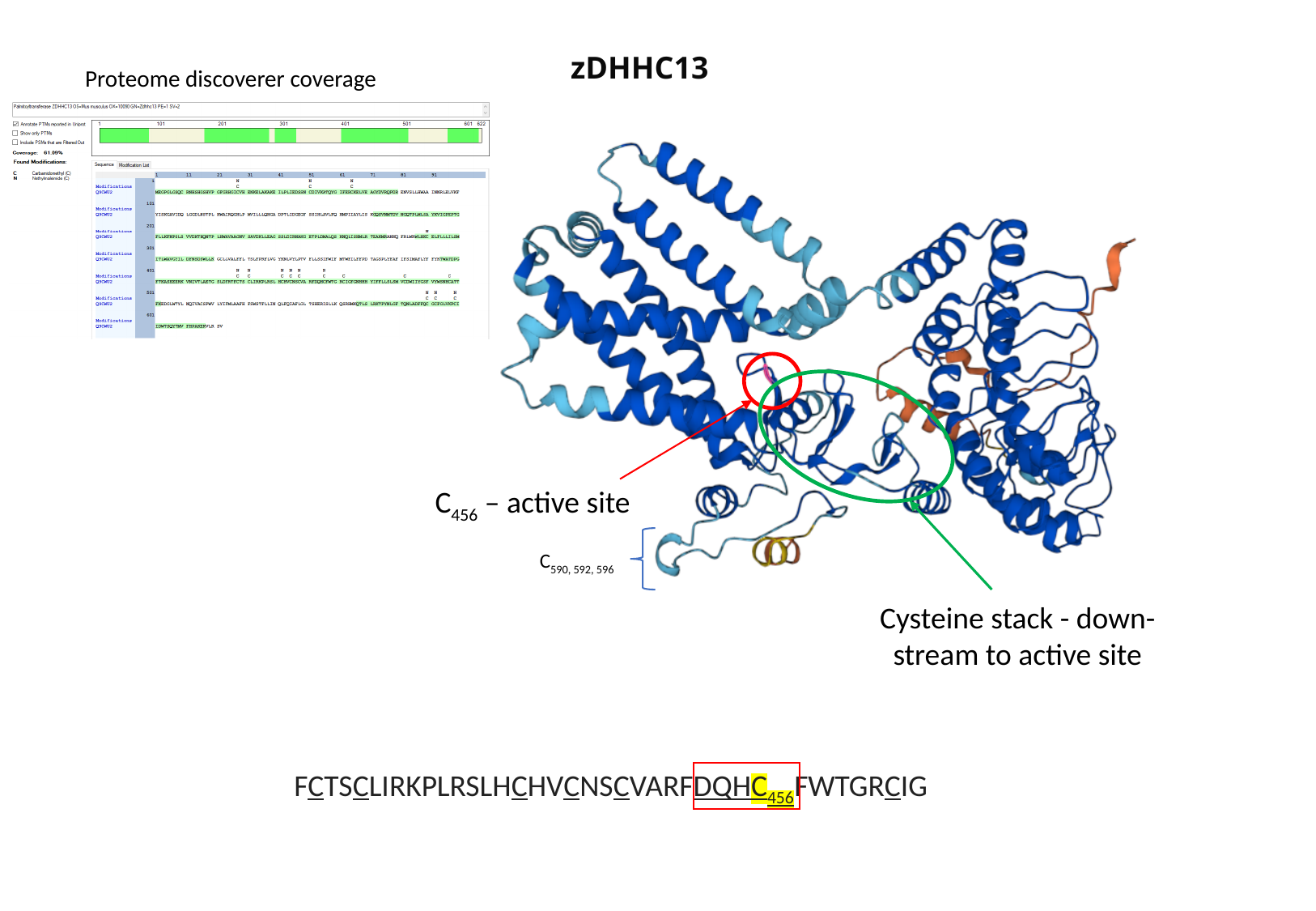

zDHHC13
Proteome discoverer coverage
C_456_ – active site
C_590, 592, 596_
Cysteine stack - down-stream to active site
FCTSCLIRKPLRSLHCHVCNSCVARFDQHC_456_FWTGRCIG

### Slide 13
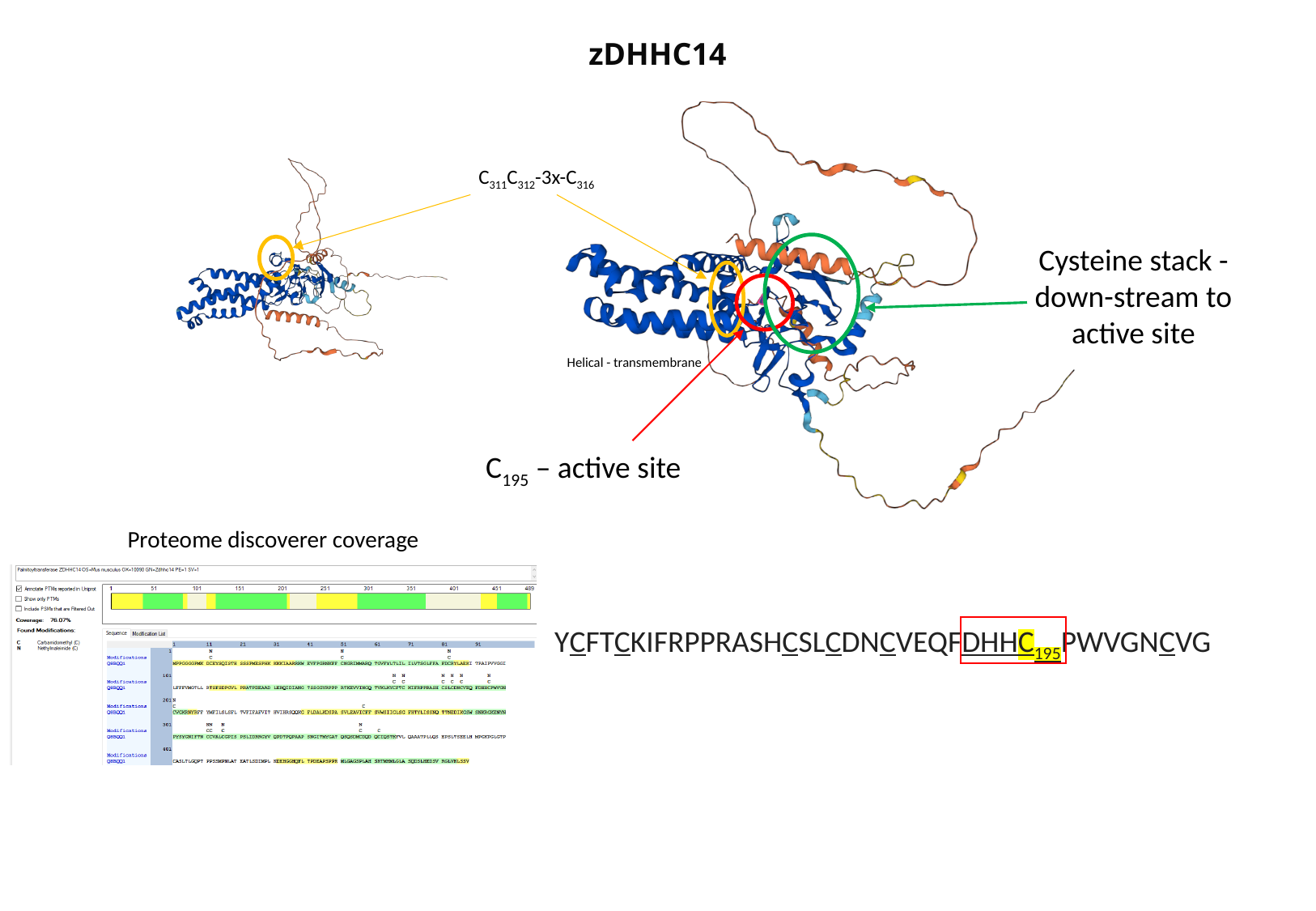

zDHHC14
C_311_C_312_-3x-C_316_
Cysteine stack - down-stream to active site
Helical - transmembrane
C_195_ – active site
Proteome discoverer coverage
YCFTCKIFRPPRASHCSLCDNCVEQFDHHC_195_PWVGNCVG

### Slide 14
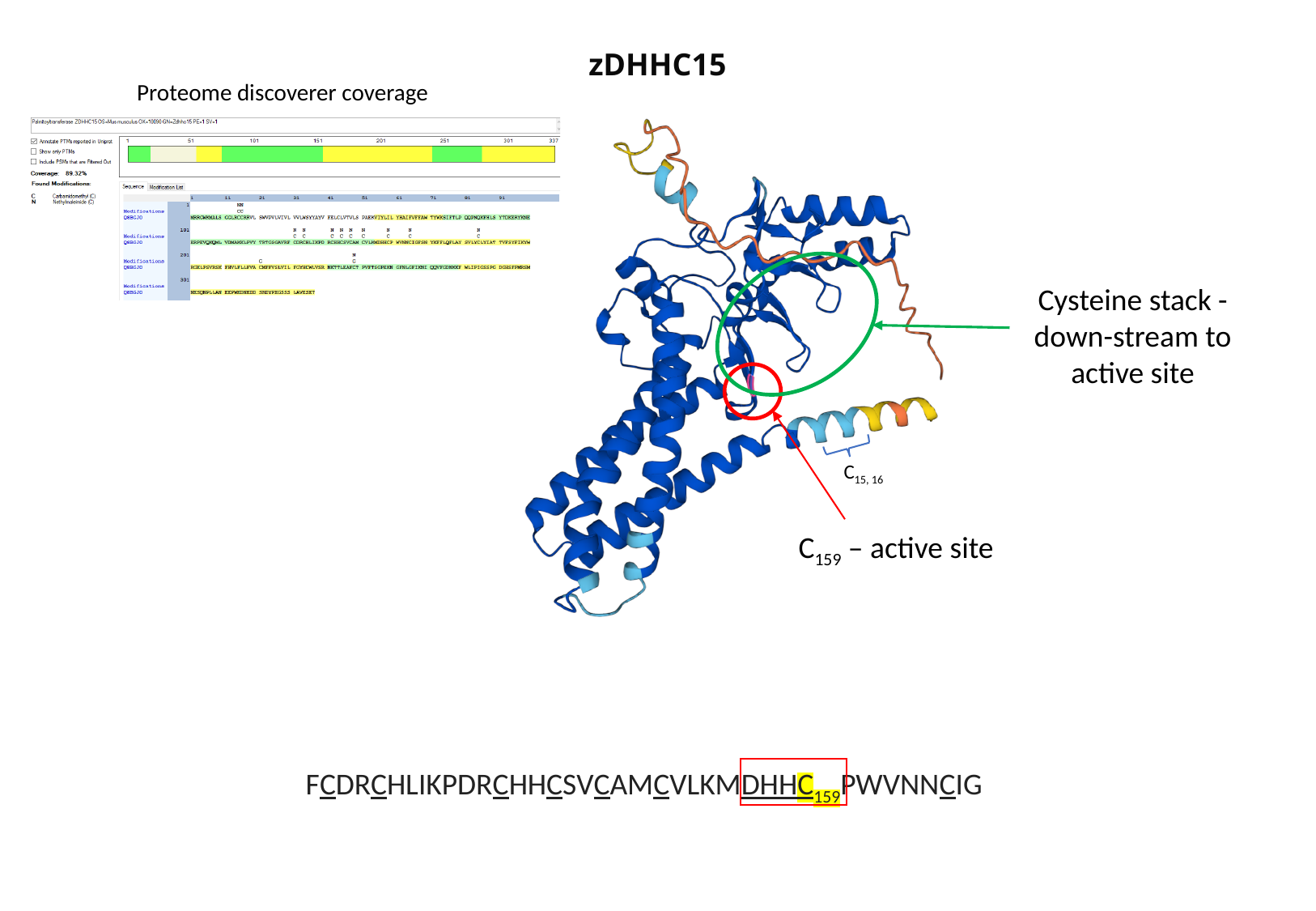

zDHHC15
Proteome discoverer coverage
Cysteine stack - down-stream to active site
C_15, 16_
C_159_ – active site
FCDRCHLIKPDRCHHCSVCAMCVLKMDHHC_159_PWVNNCIG

### Slide 15
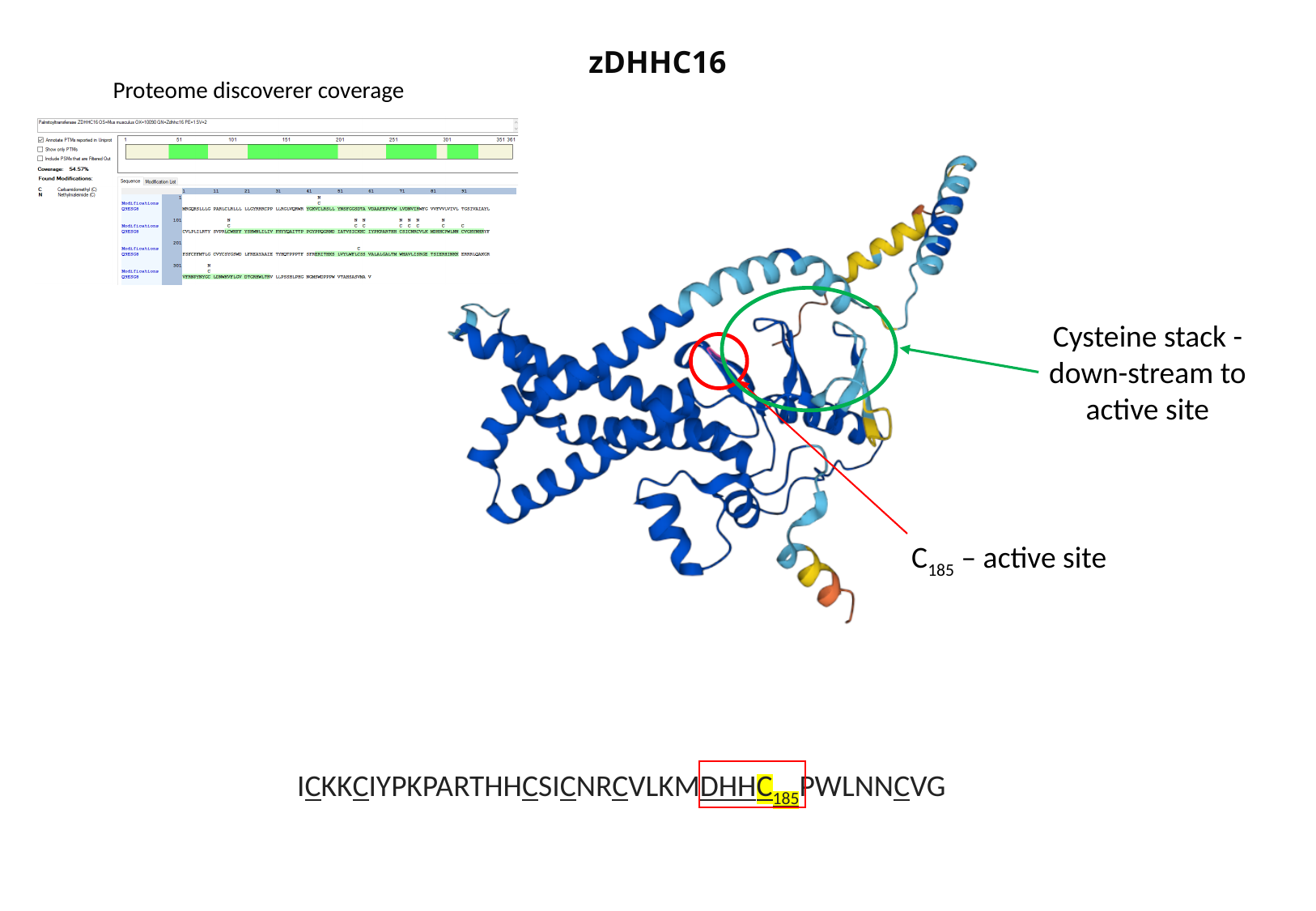

zDHHC16
Proteome discoverer coverage
Cysteine stack - down-stream to active site
C_185_ – active site
ICKKCIYPKPARTHHCSICNRCVLKMDHHC_185_PWLNNCVG

### Slide 16
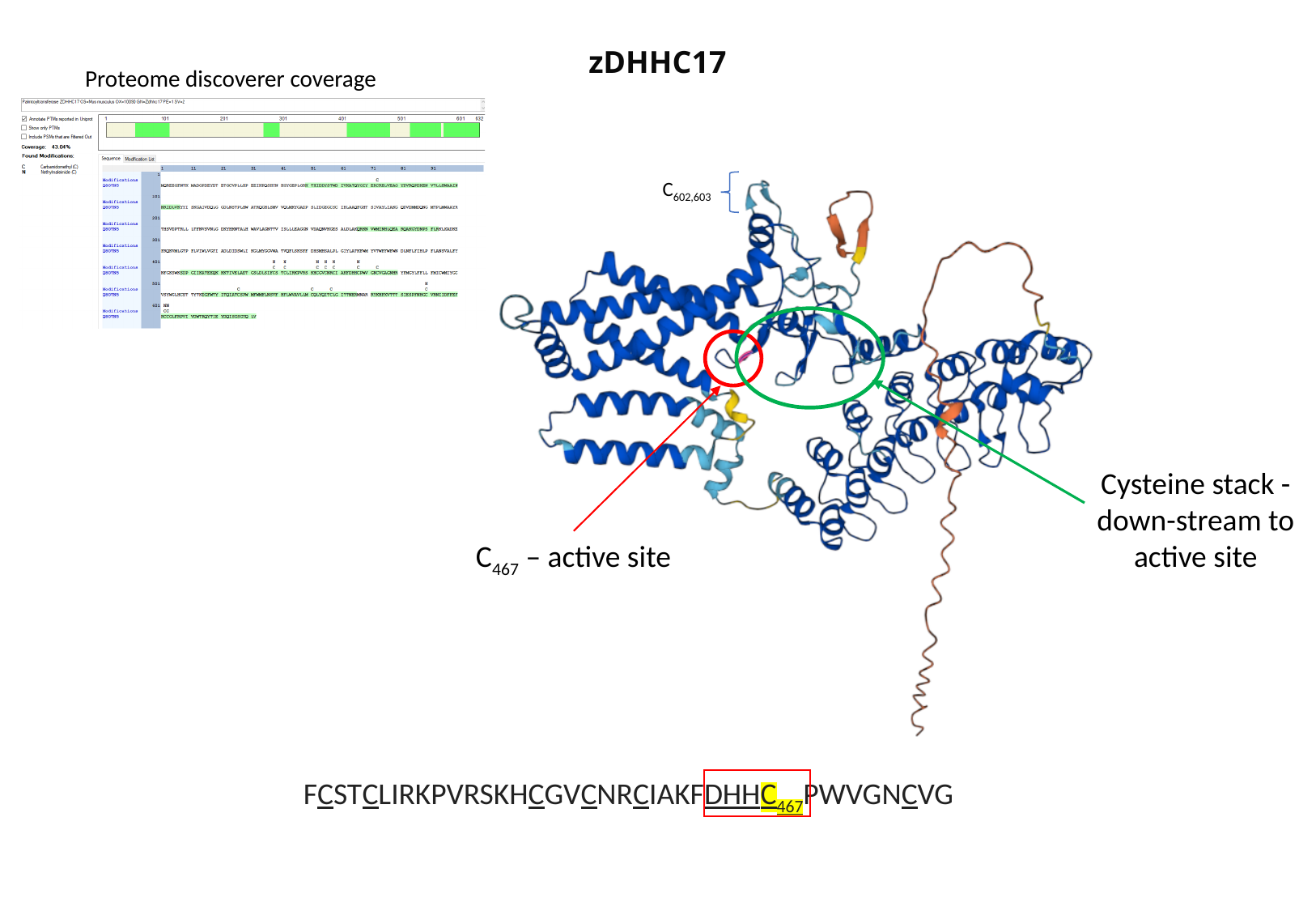

zDHHC17
Proteome discoverer coverage
C_602,603_
Cysteine stack - down-stream to active site
C_467_ – active site
FCSTCLIRKPVRSKHCGVCNRCIAKFDHHC_467_PWVGNCVG

### Slide 17
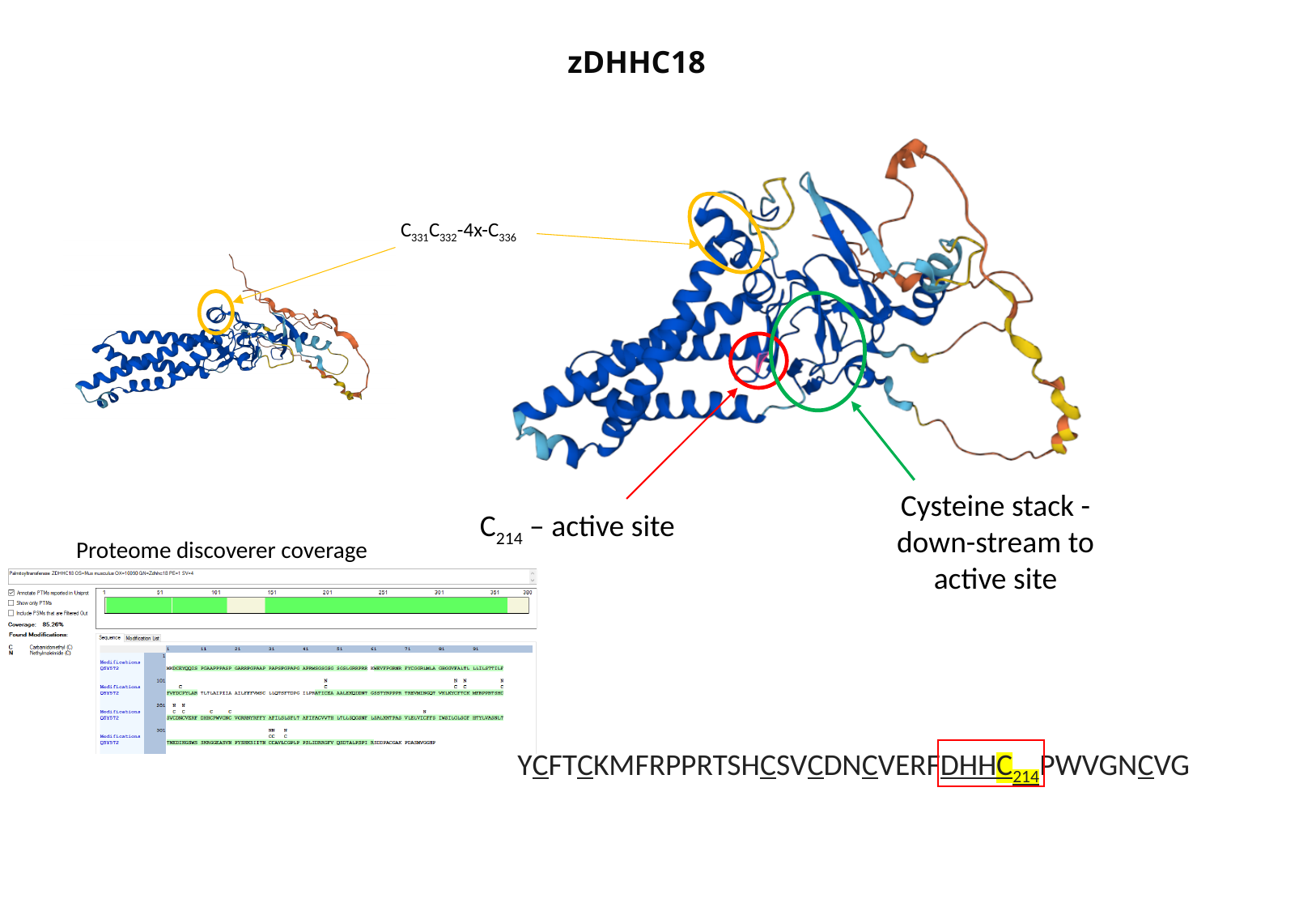

zDHHC18
C_331_C_332_-4x-C_336_
Cysteine stack - down-stream to active site
C_214_ – active site
Proteome discoverer coverage
YCFTCKMFRPPRTSHCSVCDNCVERFDHHC_214_PWVGNCVG

### Slide 18
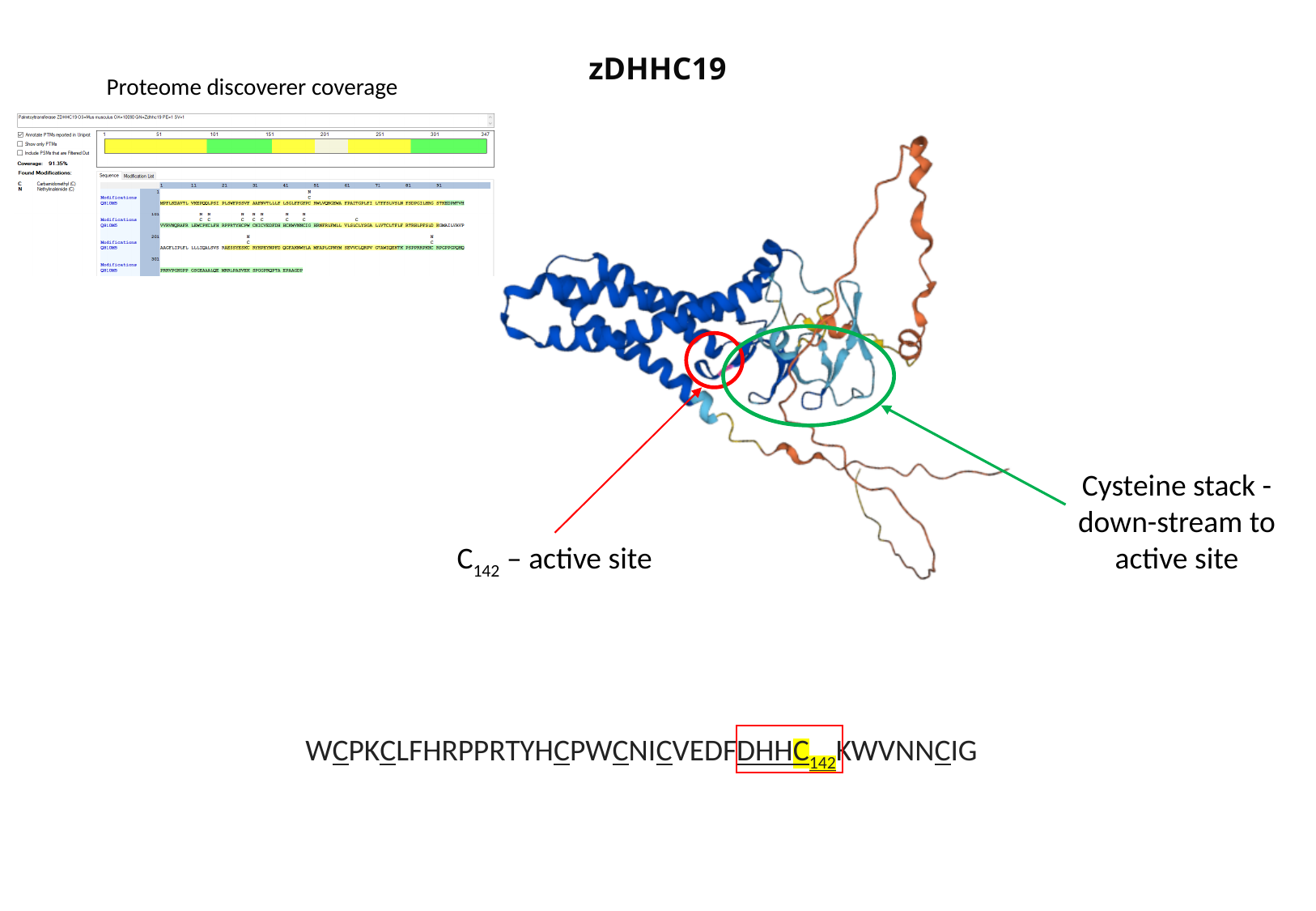

zDHHC19
Proteome discoverer coverage
Cysteine stack - down-stream to active site
C_142_ – active site
WCPKCLFHRPPRTYHCPWCNICVEDFDHHC_142_KWVNNCIG

### Slide 19
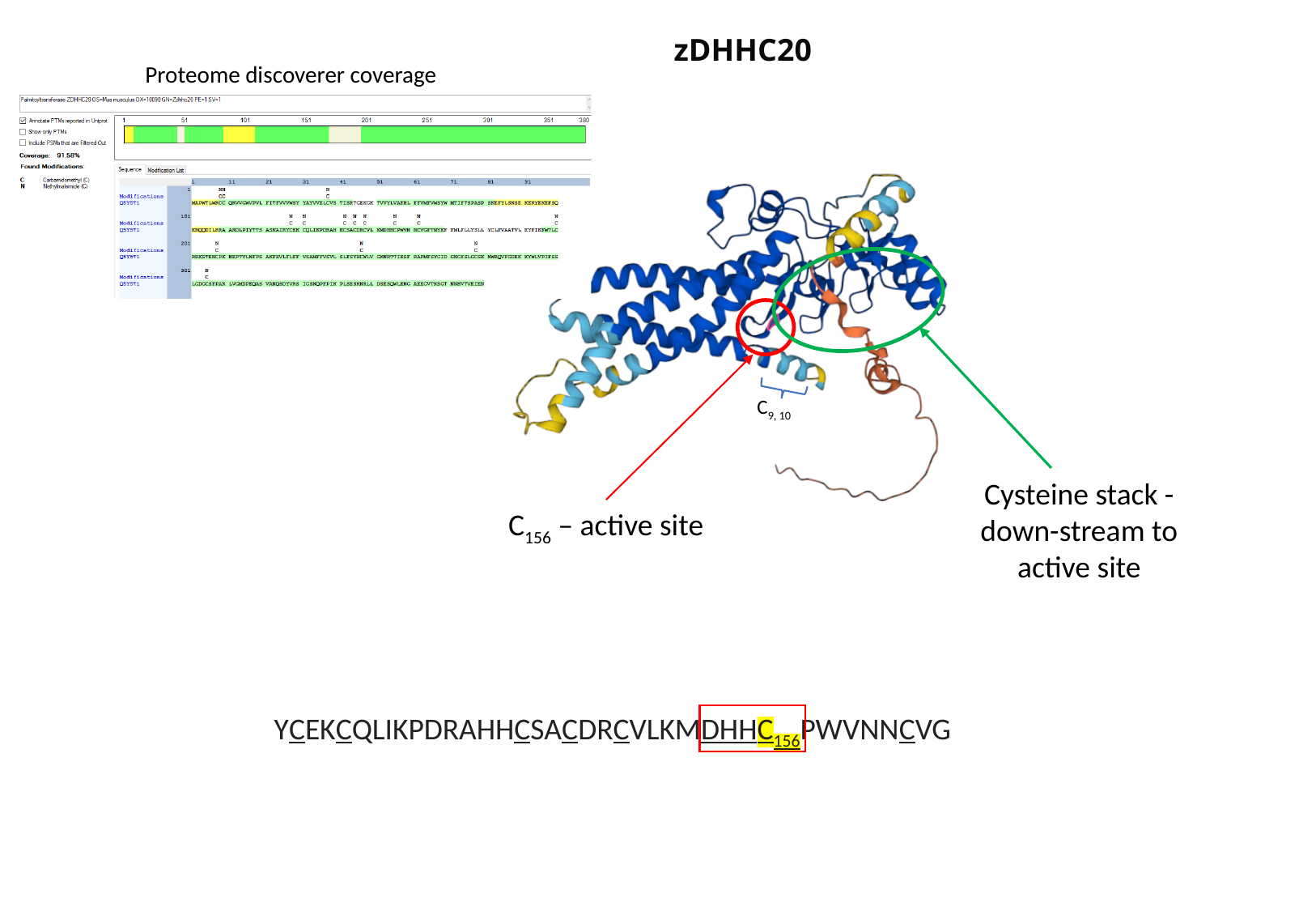

zDHHC20
Proteome discoverer coverage
C_9, 10_
Cysteine stack - down-stream to active site
C_156_ – active site
YCEKCQLIKPDRAHHCSACDRCVLKMDHHC_156_PWVNNCVG

### Slide 20
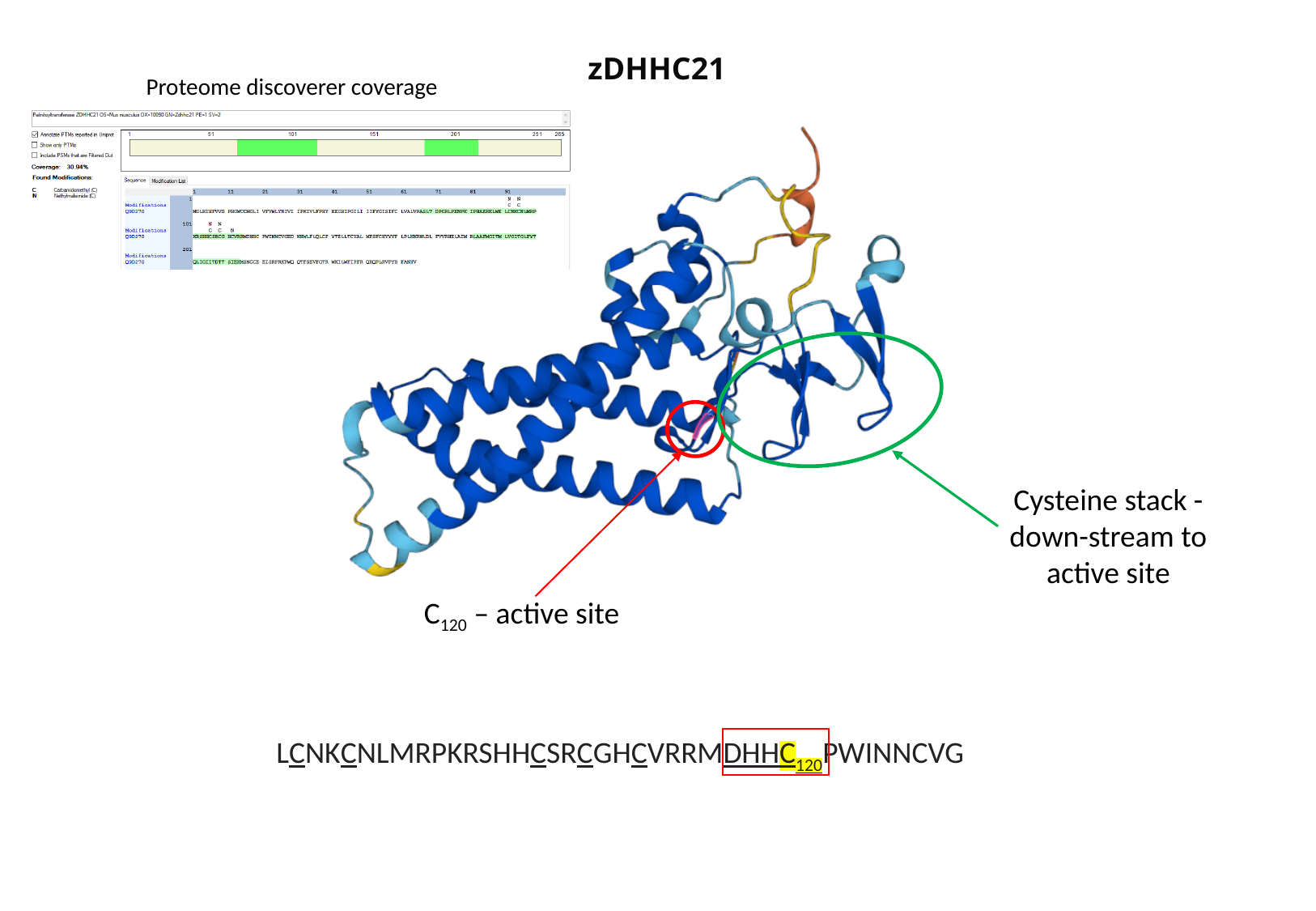

zDHHC21
Proteome discoverer coverage
Cysteine stack - down-stream to active site
C_120_ – active site
LCNKCNLMRPKRSHHCSRCGHCVRRMDHHC_120_PWINNCVG

### Slide 21
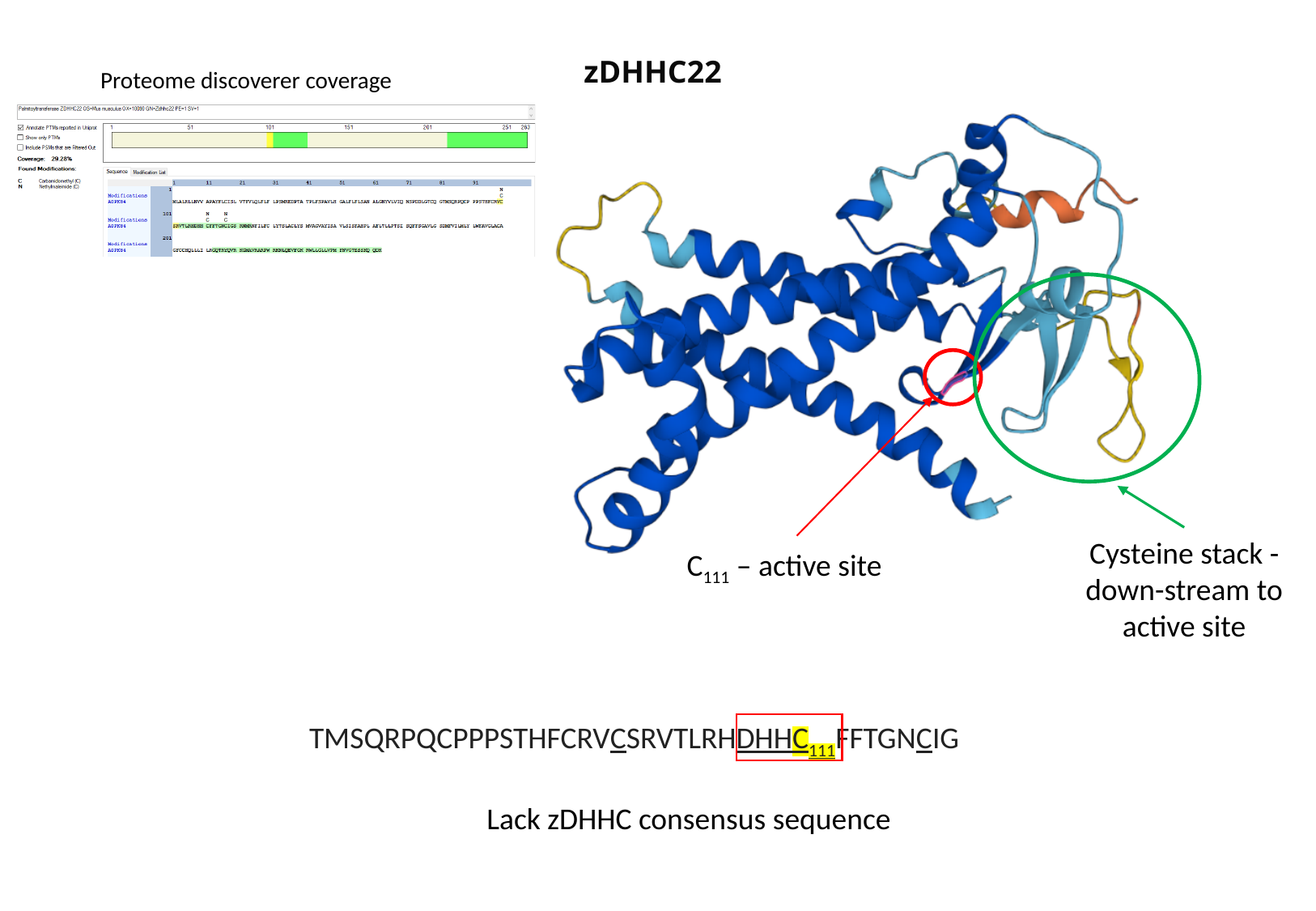

zDHHC22
Proteome discoverer coverage
Cysteine stack - down-stream to active site
C_111_ – active site
TMSQRPQCPPPSTHFCRVCSRVTLRHDHHC_111_FFTGNCIG
Lack zDHHC consensus sequence

### Slide 22
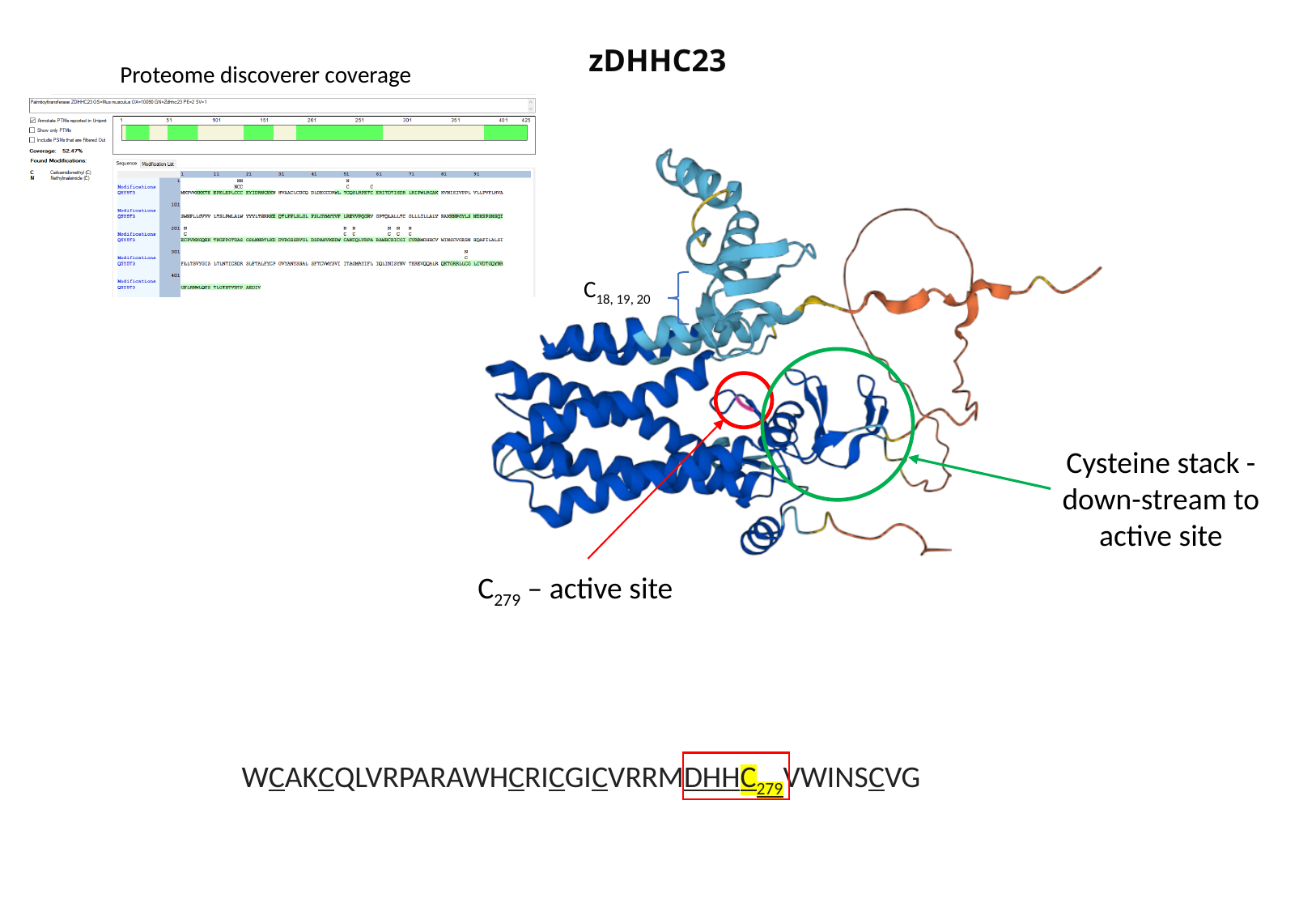

zDHHC23
Proteome discoverer coverage
C_18, 19, 20_
Cysteine stack - down-stream to active site
C_279_ – active site
WCAKCQLVRPARAWHCRICGICVRRMDHHC_279_VWINSCVG

### Slide 23
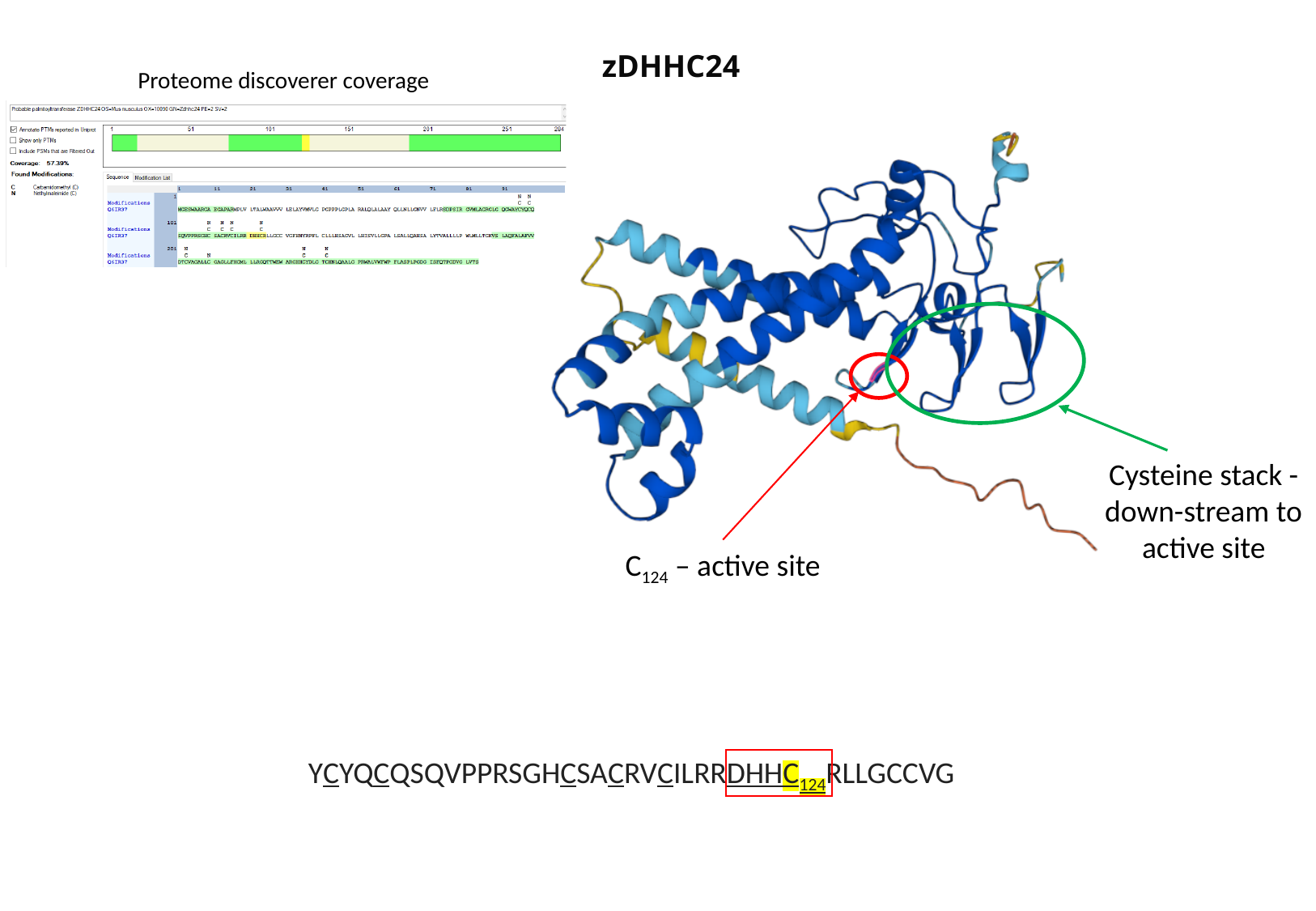

zDHHC24
Proteome discoverer coverage
Cysteine stack - down-stream to active site
C_124_ – active site
YCYQCQSQVPPRSGHCSACRVCILRRDHHC_124_RLLGCCVG

### Slide 24
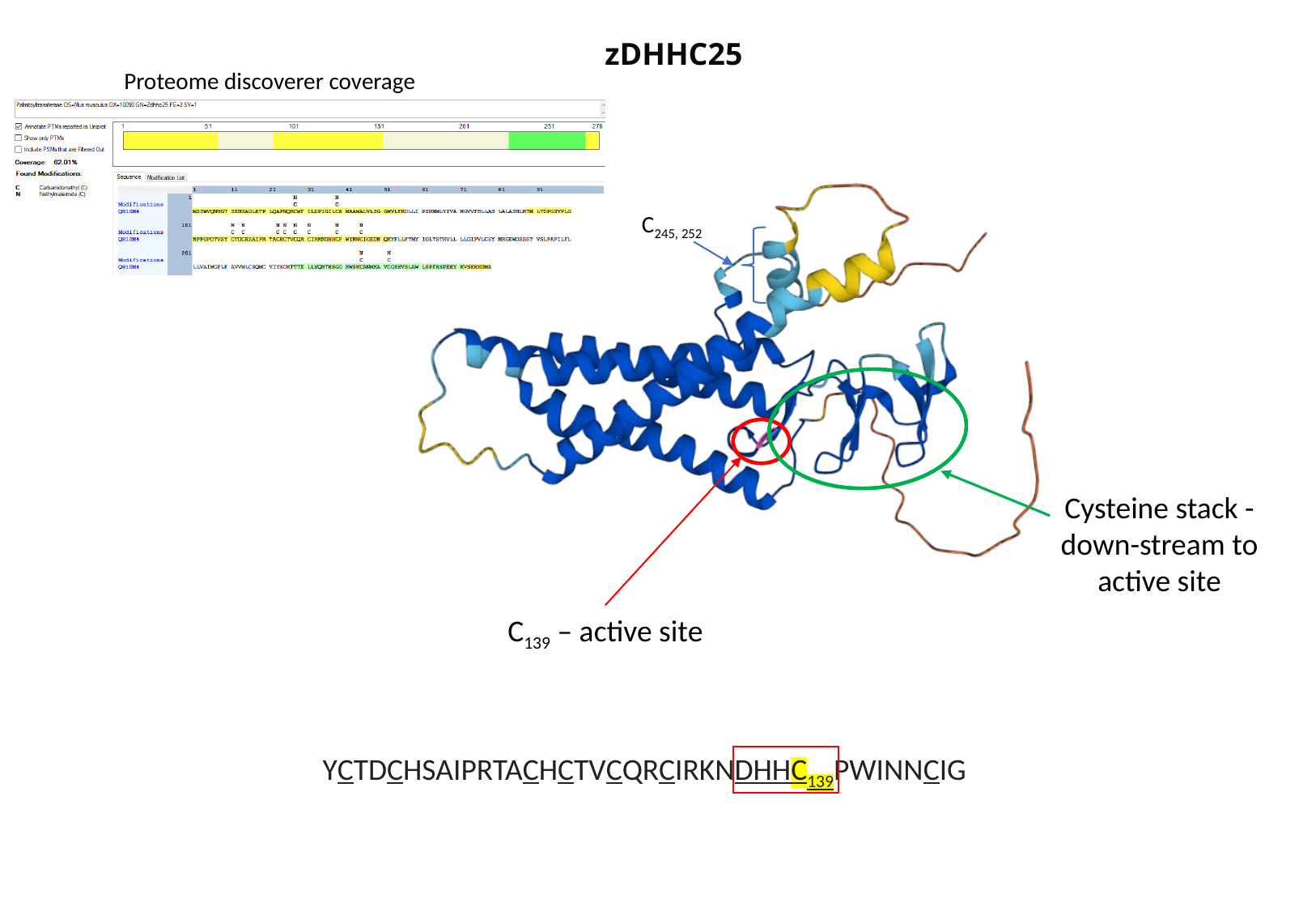

zDHHC25
Proteome discoverer coverage
C_245, 252_
Cysteine stack - down-stream to active site
C_139_ – active site
YCTDCHSAIPRTACHCTVCQRCIRKNDHHC_139_PWINNCIG
